# KCNQ1 regulates human neuronal development through mitochondrial and insulin signalling pathways

**DOI:** 10.64898/2025.12.04.692265

**Authors:** Dorothea Schall, Baran E. Güler, Siamand Alibrahim, Mahmudul Hasan, Joanna Widomska, Claudio Acuna, Mairéad Sullivan, Ralph Röth, Chang Liu, Linda Blaier, Katalin Vincze, Karin Burau, Daniela Mauceri, Marcin Luzarowski, Ágota Apáti, János M. Réthelyi, Geert Poelmans, Gudrun A. Rappold, Barbara Franke, Jeffrey C. Glennon, Simone Berkel

**Author notes:** Corresponding author: Dr. Simone Berkel, Im Neuenheimer Feld 366, 69120 Heidelberg.

## Abstract

*KCNQ1* encodes a voltage-gated potassium channel implicated in various peripheral and neurological disorders, yet its role during human neuronal development remains unclear. To investigate this, we generated *KCNQ1* knockouts (KO) in human induced pluripotent stem cell lines and differentiated them into neural stem cells (NSCs) and cortical neurons. *KCNQ1*-deficient NSCs showed impaired neurite outgrowth, linked to reduced cell adhesion and disrupted neural cell adhesion molecule (NCAM) signalling. This phenotype was reproduced in wild-type and heterozygous lines by pharmacological KCNQ1 inhibition. Whole transcriptome, proteome, and follow-up analyses revealed mitochondrial dysfunction in KO NSCs, including reduced mitochondrial copy number and ATP synthase expression. Additionally, evidence was obtained for an impairment of insulin signalling in NSCs and neurons, with diminished insulin receptor gene expression and perturbation of key downstream signalling pathways (RAS-MAPK, PI3K-AKT). In neurons, *KCNQ1* loss resulted in decreased synaptic activity and a more immature gene expression profile. Overall, our work reveals a novel role for KCNQ1 in human neurodevelopment by regulating cell adhesion, mitochondrial function, and insulin signalling. This work increases our understanding of KCNQ1 function in neurons and its contribution to neurological phenotypes observed in patients with *KCNQ1*-related diseases.

## Introduction

KCNQ1 is the pore-forming alpha subunit of a voltage-gated potassium channel complex. It interacts with KCNE family members, which act as beta subunits, to assemble the functional Kv7.1 potassium channel^1,2^. This channel generates a slow delayed rectifier current^3^.

KCNQ1 is ubiquitously expressed in humans, though its function varies in different tissues^2^. In the heart, it is responsible for the main repolarizing potassium current, and for the regulation of the cardiac action potential duration^3^. In peripheral organs such as the stomach, intestines, kidney, thyroid gland, and the respiratory system, KCNQ1 is expressed in epithelial cells. Here, it facilitates the transport of ions, hormones, and water, maintaining potassium homeostasis^2^. In the pancreas, it modulates insulin secretion from beta cells by regulating potassium channel currents^4^. KCNQ1 is also expressed in multiple regions of the developing and adult brain in mice and humans. In neuronal tissues, it interacts with KCNE1 and is hypothesized to influence neuronal excitability^5^. A recent study showed Kcnq1 expression in rat brain microvessels and primary endothelial cells, suggesting a role in regulating blood-brain barrier permeability^6^. However, the precise function of KCNQ1 in the brain and neurons remains largely unknown.

*KCNQ1* is an imprinted gene, with only the maternal transcript expressed in most human tissues. The paternal transcript is silenced by the non-coding *KCNQ1 overlapping transcript 1* (*KCNQ1OT1*) which is located in antisense orientation within intron 10 of the *KCNQ1* gene^7^. *KCNQ1* has been linked to various diseases, primarily those affecting the heart, such as familial atrial fibrillation^8^, long^9^ and short^10^ QT syndrome, Romano-Ward syndrome^11^, and Jervell and Lange-Nielsen syndrome; the latter presents with deafness^4^. Additionally, *KCNQ1* is recognized as a susceptibility gene for diabetes mellitus type 2 (DM2)^12,13^ in genome-wide association studies (GWAS). Genetic variants in *KCNQ1* have been identified in patients with epilepsy and can increase risk for sudden unexplained death^5^. Moreover, the integration of GWAS findings on obsessive compulsive disorder (OCD) has highlighted insulin-related signalling as a key pathway in OCD etiology and identified KCNQ1 as a crucial molecule in this process^14,15^.

Notably, comorbidities exist among the various diseases in which *KCNQ1* is involved. Genetic variants in *KCNQ1* have been identified in patients with long QT syndrome (LQTS), who also experience epilepsy^16–18^. Individuals with LQTS are at increased risk of developing DM2, hearing loss, epilepsy, and psychiatric disorders compared to the general population^19,20^. Patients with DM2 were found to exhibit mild reductions in cognitive function, and an elevated risk for dementia (50–100%) compared to matched non-diabetic controls^21–23^. They performed worse on tests assessing verbal and visual memory, attention, processing speed, executive function, and motor control^21,23^. DM2 is caused by dysfunctional insulin signalling leading to insulin resistance, and is linked to multiple co-morbidities including LQTS^19^, epilepsy^24^, neuropsychiatric^25,26^ and neurodegenerative disorders^23^. Recent studies provide first evidence for a shared genetic etiology of these multi-morbid conditions, with mounting evidence pointing to disturbed insulin signalling as a common mechanism^27,28^. Given *KCNQ1*’s involvement in LQTS, DM2, epilepsy, and OCD, it may also play a role in comorbidities between diseases of peripheral organs and the central nervous system. As KCNQ1 regulates insulin secretion in pancreatic cells^4^, and contributes to DM2 and related conditions, we hypothesize that it may also influence insulin signalling in the brain and impacts disease comorbidities.

In this study, we investigated KCNQ1 in human neuronal stem cells (NSCs) and cortical neurons to better understand its functional role, and to unravel its impact on insulin signalling. The majority of pathogenic variants in *KCNQ1* are loss-of-function; therefore, genetic deletion of *KCNQ1* helps to assess its physiological role, and to elucidate disease mechanisms. To achieve this, we generated knockouts (KOs) of *KCNQ1* in human induced pluripotent stem cells (iPSCs) that were subsequently differentiated into NSCs and cortical neurons. The loss of *KCNQ1* impaired early neuronal differentiation by reducing neurite outgrowth and affected neuronal function. Transcriptome and proteome analyses revealed disruptions in gene transcription, protein translation, mitochondrial function and insulin signalling, involving key downstream effectors such as RAS, mitogen-activated protein kinase (MAPK, also known as “extracellular signal-regulated kinase” (ERK)), phosphatidylinositol 3-kinase (PI3K), and Akt serine/threonine kinase (AKT). Overall, our findings demonstrate for the first time a role for KCNQ1 in neuronal development and function.

## Results

### Generation and differentiation of *KCNQ1*-KO lines

To investigate KCNQ1 function in human neuronal cells, we used an iPSC line from a healthy human donor^29^, and generated *KCNQ1*-KO lines with the CRISPR/Cas9 system by deleting exon 4 and 5 with two different guide RNA (gRNA) combinations (Fig. 1a). We obtained one homozygous and three heterozygous *KCNQ1*-KO lines, as well as two unedited wildtype (WT) isogenic control lines (Fig. 1b, Suppl. Fig. 1). The loss of exons 4 and 5 leads to a frameshift and a premature stop codon, and subsequently to the loss of *KCNQ1*. To confirm the gene KO, we quantified *KCNQ1* expression by qPCR and found reduced expression in two heterozygous (hetKO 1, hetKO 2) and in the homozygous KO (homKO 1) iPSC lines when compared to the respective controls (WT 1, WT 2) (Fig. 1c). *KCNQ1* expression was not decreased in the heterozygous KO line hetKO 3. Since *KCNQ1* is an imprinted gene for which only the maternal allele is actively transcribed in most tissues, we hypothesized that a deletion on different alleles is the reason for different *KCNQ1* expression levels in different hetKO lines. Imprinting of *KCNQ1* is generally not affected by the reprogramming of somatic cells into iPSCs^30^. We distinguished the two *KCNQ1* alleles based on heterozygous SNPs (Suppl. Fig. 2a) and found that the hetKO 1 and hetKO 2 lines carry the deletion on allele A, whereas in hetKO 3, the deletion is on allele B (Suppl. Fig. 2b). Combining these results with *KCNQ1* expression levels, we conclude that allele A is the active allele (since *KCNQ1* expression is reduced in the heterozygous KOs carrying the deletion on this allele), and allele B is the inactive allele (since hetKO 3 does not show a reduction in *KCNQ1* expression). We confirmed good iPSC quality by analysing chromosomal stability by karyotyping (Suppl. Fig. 3a), excluded off-target editing by sequencing of putative unspecific gRNA binding sites (Suppl. Fig. 3b), and confirmed pluripotency by trilineage differentiation and pluripotency marker protein expression analysis (Suppl. Fig. 3c, d).

**Figure 1:**
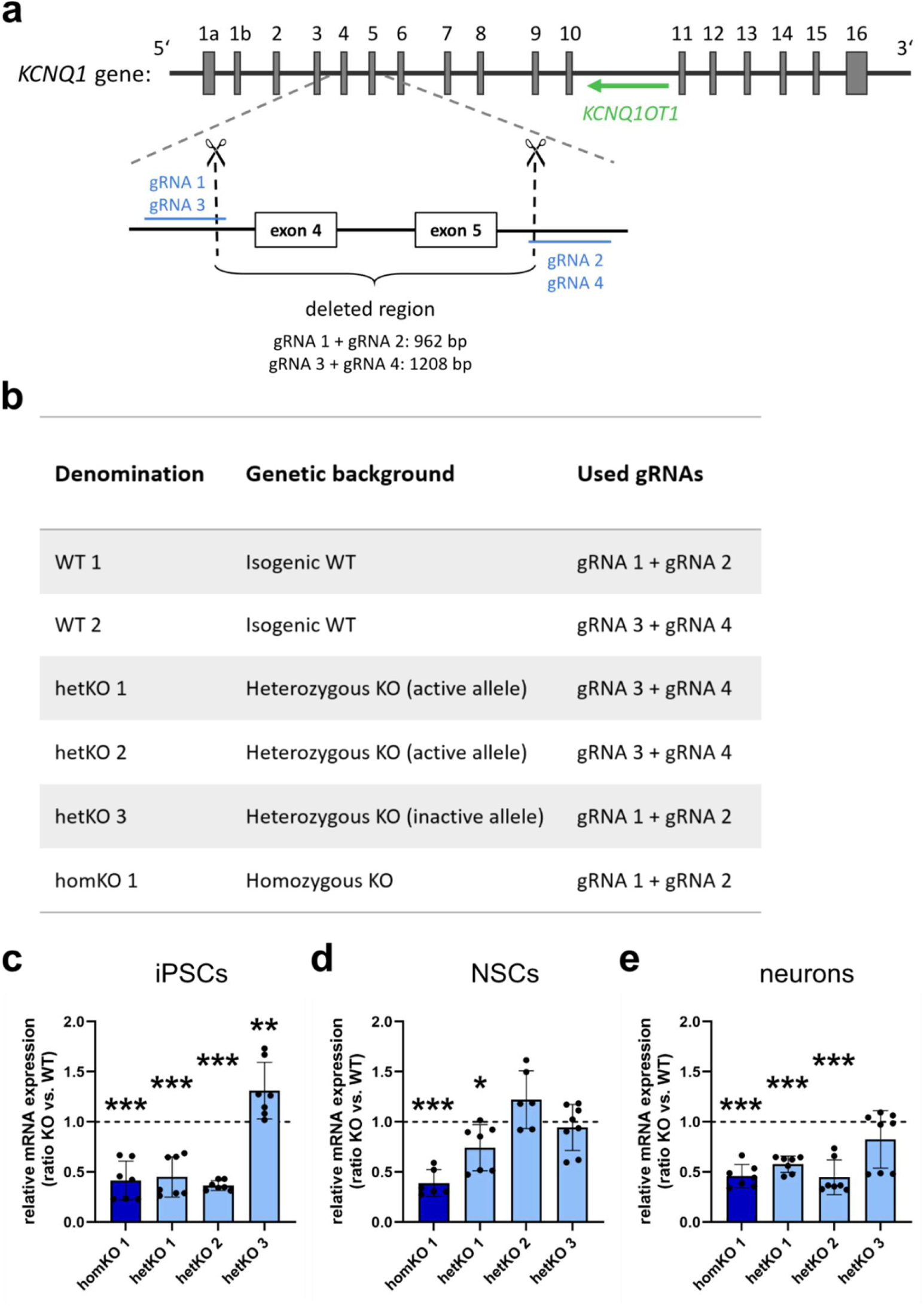
Generation of *KCNQ1*-KO cell lines and relative *KCNQ1* expression in iPSCs, NSCs and neurons. **a** Illustration of the *KCNQ1* gene and the induced deletion. Exons 4 and 5 were deleted by using two gRNAs flanking both exons. *KCNQ1OT1* is the non-coding *KCNQ1* overlapping transcript 1 in antisense orientation. **b** List of generated *KCNQ1*-KO cell lines and isogenic controls. **c-e** *KCNQ1* expression on RNA level in iPSCs (c), NSCs (d), and neurons (e). KO lines were always normalized and compared to their respective WT control (treated with the same gRNA, see 1b), indicated as dotted line. One-way ANOVA with Bonferroni-adjusted post hoc test, mean ± standard deviation (SD), n=3. **c** *P*<0.0001, F=43.33, post hoc test: homKO 1, hetKO 1, hetKO 2: *P*<0.001; hetKO 3: *P*=0.0033. **d** *P*<0.0001, F=15.23, post hoc test: homKO 1: *P*<0.001, hetKO 3: *P*>0.99, hetKO 1: *P*=0.0385, hetKO 2: *P*=0.1222. **e** *P*<0.0001, F=20.79, post hoc test: homKO 1, hetKO 1, hetKO 2: *P*<0.001; hetKO 3: *P*=0.1315.

For the investigation of KCNQ1 function in neuronal cells, we differentiated the iPSCs into NSCs and cortical neurons^31^. The differentiated NSCs at passages 5-10 expressed proteins typical for NSCs, such as Nestin, PAX6, and SOX2 (Suppl. Fig. 4 a, b). NSCs were then further differentiated into neurons and quality was assessed focusing on morphology, neuronal activity, and protein expression. Patch clamp electrophysiology analysis revealed that most neurons showed spontaneous firing of action potentials (APs) under physiological conditions (2 mM Ca++/1 mM Mg++ in the external solution) and all neurons fired APs upon positive current injections (Suppl. Fig. 5a-d). Differentiated neurons also displayed prominent excitatory postsynaptic currents (EPSCs) under physiological conditions (Suppl. Fig. 5e, f). Immunofluorescence microscopy of the differentiated neuronal cultures revealed β3-tubulin, Tau, and MAP2 positive neurons accompanied by a few GFAP positive glial cells (Suppl. Fig. 5g).

*KCNQ1* expression was analysed in NSCs and neurons of all cell lines. In NSCs, significantly diminished *KCNQ1* expression was detected in homKO 1 and hetKO 1 (Fig. 1d). Neurons showed reduced *KCNQ1* expression in homKO 1, hetKO 1, and hetKO 2 compared to their respective controls (Fig. 1e), which is in line with the expression pattern in iPSCs (Fig. 1c). HetKO 3 did not show reduced *KCNQ1* expression in iPSCs, NSCs, or neurons (Fig. 1c-e), as it carries the deletion on the inactive allele.

Notably, hetKO 2 NSCs did not exhibit reduced *KCNQ1* expression and hetKO 1 showed only some reduction, which could be caused by a loss of imprinting and expression of the second allele. To elucidate imprinting differences between cell types, we performed RT-PCR to find out which allele (WT or KO) is expressed in iPSCs, NSCs, and neurons for all different genotypes. In iPSCs, the heterozygous *KCNQ1*-KO lines exhibited monoallelic expression: hetKO 3 expressed the WT allele, and hetKO 1 and hetKO 2 the KO allele, respectively (Suppl. Fig. 6a). In NSCs and neurons, we detected biallelic expression (WT and KO) across all three heterozygous *KCNQ1*-KO lines (Suppl. Fig. 6a). These findings suggest that *KCNQ1* imprinting is at least partially lost during neural induction, leading to biallelic expression in NSCs, which persists through subsequent neuronal differentiation. To exclude the possibility that genome editing affected imprinting, we assessed the expression of the long non-coding RNA *KCNQ1OT1* — a known imprinting regulator of *KCNQ1* and additional genes — and found no evidence of differential expression. Therefore, we can rule out that differential *KCNQ1OT1* expression is the cause of the biallelic expression in NSCs and neurons (Suppl. Fig. 6b). Additionally, we identified a shorter transcript lacking exons 3-6 and 8 (NM_001406838.1) (Suppl. Fig. 6a), without existing evidence for translation into a functional protein. The presence of this transcript likely accounts for the residual *KCNQ1* expression observed in homKO 1, hetKO 1, and hetKO 2 cell lines by qPCR (Fig. 1c-e), in which exon 10-11 was targeted.

HetKO 3 represents a distinct case, as the KO was introduced on the inactive allele which is silenced by imprinting. Consequently, no reduction in *KCNQ1* expression was observed in iPSCs, where only the active WT allele is expressed, rendering hetKO 3 indistinguishable from WT controls at this stage. However, in NSCs and neurons, the KO allele was transcribed, making hetKO 3 molecularly distinct from WT cells, even though *KCNQ1* expression was not found reduced in general. Given this imprinting-dependent shift in expression, hetKO 3 was not treated as a WT control, and was included in subsequent analyses.

After generation and differentiation of *KCNQ1*-KO NSCs and neurons and their quality assessment, we next investigated KCNQ1 function at the NSC stage.

### The role of KCNQ1 in early neuronal development

#### KCNQ1 loss and inhibition reduced neurite outgrowth during early neuronal differentiation

To investigate whether KCNQ1 has an impact on early neuronal differentiation, we focused on the critical stage of neurodevelopment marked by the emergence of immature neurites that subsequently differentiate into axons and dendrites. Hence, we measured neurite length of NSCs during the initial 48h of neuronal differentiation using live cell imaging with the Incucyte system. We selected homKO 1, hetKO 1, and their WT controls for these experiments, as we expected the strongest phenotypes in these KO cell lines. Compared to the respective isogenic controls, significantly shorter neurite length was observed in both homKO 1 and hetKO 1 lines (Fig 2a, b), suggesting that KCNQ1 promotes early neuronal differentiation. To validate this finding and ensure reproducibility, we employed an independent differentiation protocol^32^. Consistent with previous findings, both KO lines displayed reduced neurite length relative to controls (Suppl. Fig. 7). This data shows that KCNQ1 is involved in early neuronal development.

**Figure 2:**
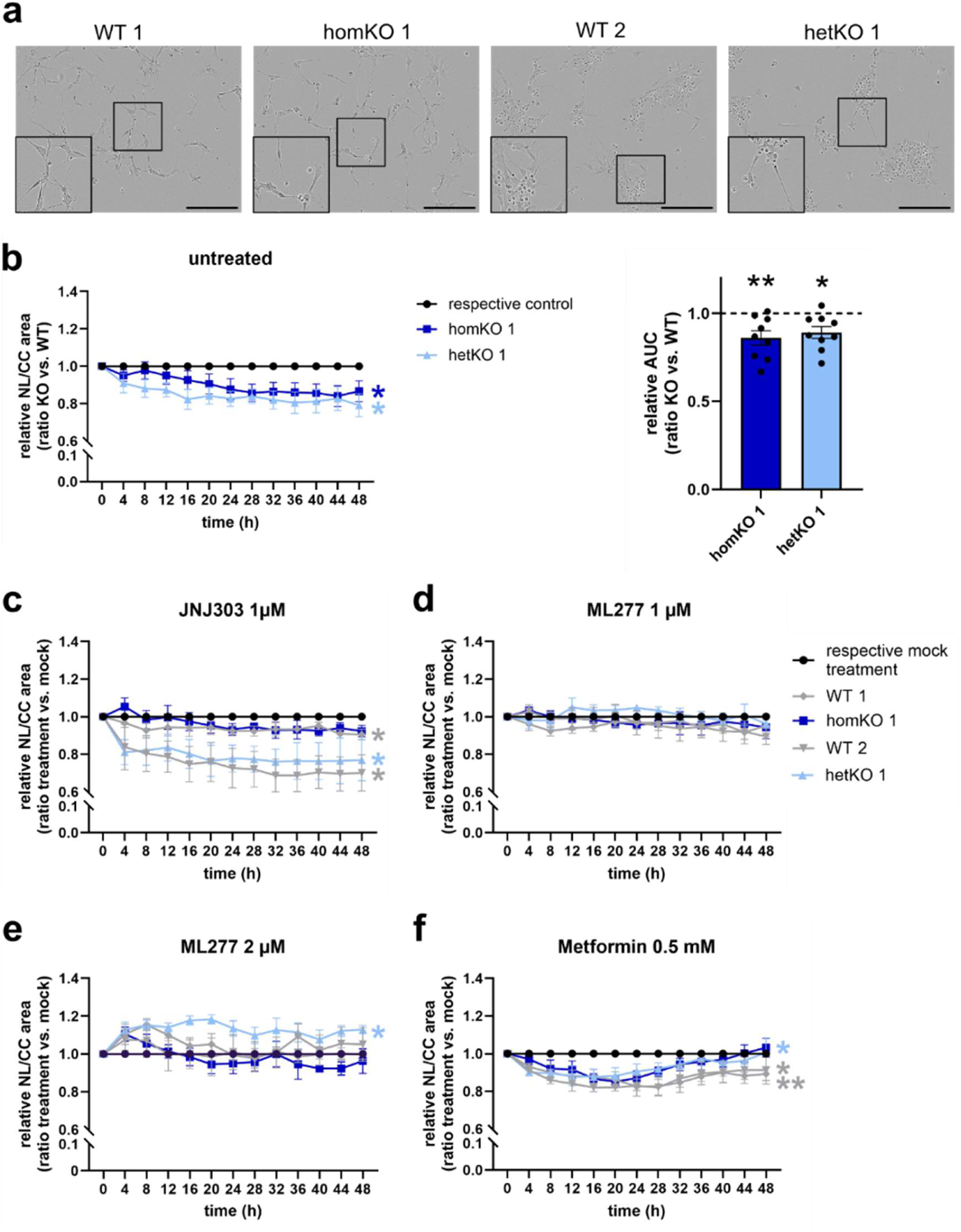
Neurite outgrowth measurement during early neuronal differentiation. **a** Representative pictures of differentiating NSCs 48h after differentiation start. Scale bar: 200 µm. **b** Neurite outgrowth was measured using the IncuCyte system within the first 48h after starting neuronal differentiation. The relative neurite length per cell body cluster area (NL/CC) for each KO line was compared to its respective control (homKO 1 vs. WT 1, hetKO 1 vs WT 2). Left: Two-way ANOVA (time x cell line) with Bonferroni-adjusted post hoc test, mean ± standard error of the mean (SEM), n=9. Asterisks indicate significances of ANOVA for factor cell line: homKO 1: *P*=0.0382, F=5.103; hetKO 1: *P*=0.004, F=11.29. Post hoc test: homKO 1: all *P*>0.05, hetKO 1: 24h: *P*=0.0316, 28h: *P*=0.0336, all others *P*>0.05. Right: The area under curve (AUC) was calculated, followed by a one-way ANOVA with Bonferroni-adjusted post hoc test, mean ± SEM, n=9, *P*=0.0006, F=7.608, post hoc test: homKO 1: *P*=0.0014, hetKO 1: *P*=0.0129. Both KO lines are normalized and compared to their respective WT control, indicated as dotted lines. **c-f** Treatment of NSCs with KCNQ1 antagonist JNJ303 (IC50= 64 nM) (c), agonist ML277 (EC50= 270 nM) (d: 1 µM, e: 2µM), and metformin (f). NSCs were treated with different compounds at the start of neuronal differentiation, and neurite outgrowth was measured for 48h. Treated cell lines were compared to their respective mock treatment (DMSO or water). Two-way ANOVA (time x treatment) with Bonferroni-adjusted post hoc test, mean ± SEM, n=4. Asterisks indicate significances of ANOVA for factor cell line: **c** WT 1: *P*=0.0336, F=13.71; homKO 1: *P*=0.3068, F=1.247; WT 2: *P*=0.032, F=7.723; hetKO 1: *P*=0.0413, F=6.703. Post hoc test: all *P*>0.05. **d** WT 1: *P*=0.2347, F=1.745; homKO 1: *P*=0.4198, F=0.7499; WT 2: *P*=0.4882, F=0.545; hetKO 1: *P*=0.5443, F=0.4128. Post hoc test: all *P*>0.05. **e** WT 1: *P*=0.1358, F=3.473; homKO 1: *P*=0.6623, F=0.2217; WT 2: *P*=0.71, F=0.1521; hetKO 1: *P*=0.0005, F=47.73. Post hoc test: hetKO 1: 4h: *P*=0.0120, 8h: p=0.0012, 12h: *P*=0.0040, 16h: *P*=0.0001, 20h: *P*<0.0001, 24h: *P*=0.0062, 32h: *P*=0.0133, 36h: *P*=0.0376, 44h: *P*=0.0190, 48h: *P*=0.0106, 0h/28h/40h: *P*>0.05; WT 1/homKO 1/WT 2: all *P*>0.05. **f** WT 1: *P*=0.002, F=27.0; homKO 1: *P*=0.0966, F=3.874; WT 2: *P*=0.0225, F=9.309; hetKO 1: *P*=0.0346, F=7.402. Post hoc test: WT 1: 24h: *P*=0.0460; WT 2: 20h: *P*=0.0356; all others *P*>0.05.

As the loss of *KCNQ1* resulted in reduced neurite outgrowth, we hypothesized that blocking of the Kv7.1 channel might have similar effects. Therefore, we treated the homKO 1, WT 1, hetKO 1, and WT 2 NSC lines with the KCNQ1 antagonist JNJ303 and measured neurite outgrowth as above. We observed a reduced neurite length compared to mock treatment for both WT cell lines and for hetKO 1. Differentiating homKO 1 NSCs (Fig. 2c) served as a negative control, as *KCNQ1* is entirely lost in this line, and no effect of JNJ303 was observed. Taken together, we discovered that both, inhibition and loss of KCNQ1, led to reduced neurite outgrowth.

To explore if activation of KCNQ1 also impacts on neurite outgrowth, we treated the same NSC lines with the specific KCNQ1 agonist ML277 and measured no effect at a concentration of 1 µM (Fig. 2d). A concentration of 2 µM, however, selectively enhanced neurite outgrowth in hetKO 1 NSCs (Fig. 2e), rescuing the neurite growth deficit observed in this line (Fig. 2b). These findings suggest that KCNQ1 activation is sufficient to restore early neurite extension in heterozygous KO NSCs, which retain one functional allele and express residual WT *KCNQ1*. In contrast, homozygous *KCNQ1*-KO cells, lacking both alleles, did not respond to ML277 treatment, indicating that the compound’s effect is specific to KCNQ1 activity and not due to off-target actions on other ion channels. To determine, whether loss of KCNQ1 or pharmacological modulation with an antagonist or agonist impacts cellular proliferation or induces cytotoxicity, we assessed proliferation and apoptosis rates in untreated and treated NSCs. No significant differences were observed under any condition (Suppl. Fig. 8), indicating that KCNQ1 manipulation does not affect NSC viability or proliferative capacity under these conditions.

#### Homozygous *KCNQ1*-KO NSCs are unresponsive to metformin treatment

*KCNQ1* is a known susceptibility gene for DM2, a chronic metabolic disorder characterized by insulin resistance and elevated blood glucose levels. Metformin is a commonly used drug for treating insulin resistance and DM2. In neurons, metformin is known to have neuroprotective and anti-inflammatory effects by reducing oxidative stress, to enhance insulin sensitivity, and to activate AMP-activated protein kinase (AMPK) which is a critical energy sensor^33^.

As *KCNQ1* has been strongly linked to DM2, we aimed to elucidate whether it influences the mode of action of metformin in differentiating NSCs. Treatment with 0.5 mM metformin had no effect on proliferation and apoptosis (Suppl. Fig. 8). However, it led to a significant reduction in neurite length with low effect size for both WT lines and hetKO 1, compared to mock treatment (Fig. 2f). No significant effect of metformin was seen for homKO 1, suggesting that the decrease of neurite length after metformin treatment is at least partially dependent on proper KCNQ1 function.

#### KCNQ1 impacts on multiple signalling pathways in NSCs

To elucidate cellular and molecular alterations in NSCs carrying a homozygous *KCNQ1*-KO, we performed transcriptomic and proteomic analysis. RNA sequencing was performed on WT 1 and homKO 1 cell lines at passages 6 and 7 from two independent NSC inductions, yielding four replicates per line. Principal component analysis (PCA) revealed separation between homKO 1 and WT 1 NSCs, indicating distinct differences in gene expression profiles (Suppl. Fig. 9a). A total of 13,820 protein coding genes were identified for each sample, and DESeq2 was used to identify differentially expressed genes (DEGs), as described in the “Methods” section. Using a cutoff of a FDR < 0.05, we identified 72 protein coding DEGs in homKO 1 NSCs compared to WT 1; 68 genes decreased in the KO, 4 increased (Fig. 3a, Supplementary Table 1 (Tab1)). The processed and raw (FASTQ) gene expression data for the NSCs is available through the gene expression omnibus (GSE number to be allocated). *KCNQ1* and other members of the KCNQ gene family (KCNQ2-5), as well as genes coding for KCNQ1 interaction partners, such as *KCNE1* and *KCNE2,* were not significantly differentially expressed. They had a low read coverage and were excluded from downstream analysis.

**Figure 3:**
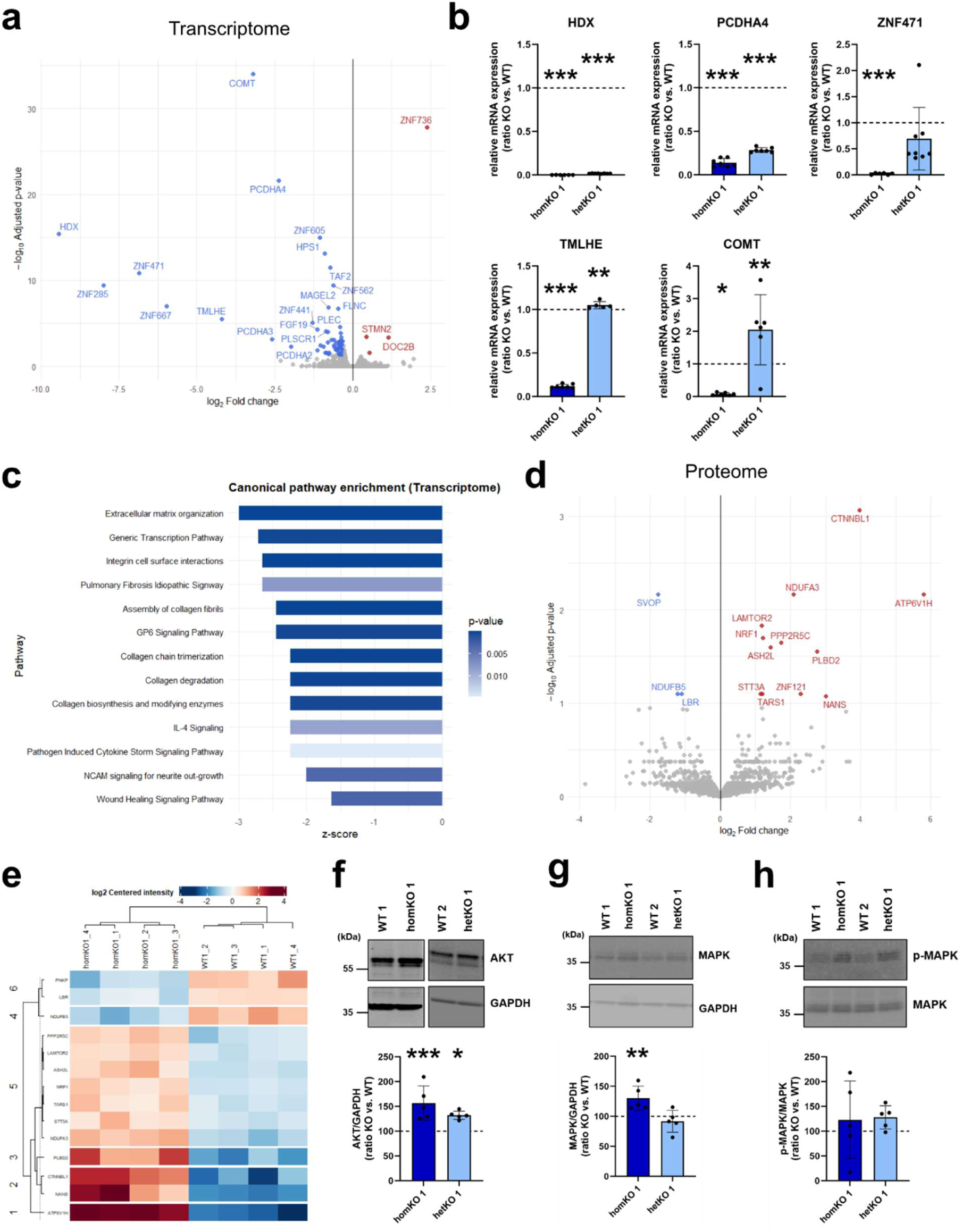
Transcriptome and proteome analysis of homozygous *KCNQ1*-KO and WT NSCs. **a** Volcano plot of transcriptome analysis from NSCs homKO 1 compared to WT 1. Genes with an adjusted *P*-value FDR < 0.05 and a log2fold change > 1.3 are shown. **b** Validation of transcriptome analysis by analysing the expression of the most significantly dysregulated genes in homKO 1 and additionally hetKO 1 compared to their respective WT control. One-way ANOVA with Bonferroni-adjusted post hoc test, mean ± SD, n=3. *HDX*: *P*<0.0001, F=1276293, post hoc test: homKO 1, hetKO 1: *P*<0.0001; *PCDHA4*: *P*<0.0001, F=1924, post hoc test: homKO 1, hetKO 1: *P*<0.0001; *ZNF471*: *P*<0.0001, F=12.79, post hoc test: homKO 1: *P*<0.0001, hetKO 1: 0.1416; *TMLHE*: *P*<0.0001, F=2710, post hoc test: homKO 1: *P*<0.0001, hetKO 1: *P*=0.0036; *COMT*: *P*<0.0001, F=13.41, post hoc test: homKO 1: *P*=0.0154, hetKO 1: 0.00641. **c** Bar graph with significant canonical pathways from transcriptomic data. All pathways with enrichment FDR *P* < 0.05 and with z-scores assigned are shown and sorted by z-score magnitude. All these pathways have negative z-scores, indicating that, based on the profile of differentially expressed genes (at FDR *P* < 0.05), they are predicted to be inhibited. **d** Volcano plot of proteome analysis from NSCs homKO 1 compared to WT 1. Gene names are shown for proteins (for better comparison) with an adjusted *P*-value FDR < 0.10 and a log2fold change > 1.3. **e** Heatmap showing significant DEPs (FDR < 0.10) for individual biological replicates. **f-h** Western blot analysis of AKT (f), MAPK (g), and p-MAPK (h) (top: representative membrane image, bottom: quantification). One-way ANOVA with Bonferroni-adjusted post hoc test, mean ± SD, n=5. AKT: *P*=0.0002, F=11.95, post hoc test: homKO 1: *P*=0.0002, hetKO 1: *P*=0.0224; MAPK: *P*=0.0022, F=7.631, post hoc test: homKO 1: *P*=0.0061, hetKO 1: *P*=0.7512; p-MAPK: *P*=0.5911, F=0.6555, post hoc test: homKO 1: *P*=0.7791, hetKO 1: *P*=0.5927. In all bar graphs KO values are normalized to their respective WT control, which are set to the value 1 or 100 and indicated by the dotted line.

Among the ten most significantly downregulated DEGs, we identified three zinc finger protein coding genes (*ZNF285, ZNF471, ZNF667*) and three protocadherin A genes (*PCDHA2, PCDHA3, PCDHA4*), pointing towards impairments in gene transcription and cell adhesion (labelled in yellow in Suppl. Table 1, Tab 1). From this top ten list and based on their function, we selected the following genes for a follow-up by qPCR analysis: *HDX (*highly divergent homeobox)*, PCDHA4* (protocadherin α4), *ZNF471* (zinc finger protein 471*), TMLHE* (trimethyllysine hydroxylase ε), and *COMT* (catechol-o-methyltransferase). To investigate whether heterozygous *KCNQ1*-KOs exhibit responses comparable to the homozygous KO, we included hetKO 1 and WT 2 NSCs alongside the previously analysed homKO 1 and WT 1 NSCs in the qPCR analysis. We confirmed downregulation of all five genes identified by RNA sequencing for homKO 1 NSCs compared to WT 1, revealing high reproducibility of the RNA sequencing results (Fig. 3b). *HDX* and *PCDHA4* were also downregulated in the hetKO 1 NSCs compared to WT 2 controls, emphasizing reduced transcriptional regulation and impaired cell adhesion as a consequence of *KCNQ1* loss (Fig. 3b).

Ingenuity Pathway Analysis (IPA) was used to identify enriched canonical pathways within the transcriptomic data, and this revealed, among others, the inhibition of extracellular matrix organization and interaction, NCAM signalling, and the generic transcription pathway (Fig. 3c, Suppl. Fig. 1 (Tab 2)). The IPA network enrichment analysis pointed towards an involvement of insulin signalling as well as to signalling cascades including RAS – MAPK/ERK, and PI3K (Suppl. Fig. 9b). RAS and PI3K signalling operate downstream of insulin receptor signalling but are also involved in many other signalling cascades. Moreover, protein kinases C and A (PKC; PKA) were affected, which are signalling molecules with a function in cell growth that are also known to regulate KCNQ1 activity^34,35^.

In addition, we performed proteomic analysis of WT 1 and homKO 1 NSCs generated in parallel with the RNA-seq samples (passages 6-7; four replicates per line). Using LC–MS, we identified 2,758 proteins in total, and differentially expressed proteins (DEPs) were determined using DEP2 as described in Methods. PCA revealed that 22.5% of the proteome-wide variance can be explained by genotype (WT versus homKO), and correlation analysis showed clear clustering of both groups (Suppl. Fig. 9c, d). Applying a lenient cutoff (FDR < 0.10) to capture biologically relevant differences, we identified 15 DEPs between homKO 1 and WT 1 NSCs, of which 3 were downregulated and 12 were upregulated (Fig. 3d; Suppl. Table 2, Tab 1). Among these, three proteins were identified with a mitochondrial function: NDUFA3 (Complex I subunit; NADH ubiquinone oxidoreductase, log₂FC = 2), NDUFB5 (Complex I subunit, log₂FC = –1.2), and NRF1 (Nuclear respiratory factor 1, a transcription factor regulating mitochondrial biogenesis and respiration, log₂FC = 1.2). The strongest expression changes were observed for ATP6V1H (log₂FC = 5.78) and CTNNBL1 (log₂FC = 3.9). ATP6V1H, a subunit of the V-ATPase proton pump, regulates organelle acidification in lysosomes and endosomes, implicating KCNQ1 in lysosomal and endosomal pH regulation. CTNNBL1, a nuclear RNA-splicing factor and NLS-binding protein, regulates gene expression and cell-cycle progression via spliceosome activation and nuclear transport, and may influence splicing of potassium channel transcripts such as the KCNQs. Additionally, enzymes linked to lipid and carbohydrate metabolism, including NANS (N-acetylneuraminic acid synthase, sialic acid biosynthesis; log₂FC = 3) and PLBD2 (phospholipase domain-containing protein 2, lipid hydrolysis; log₂FC = 2.76), were upregulated in homKO 1 NSCs. Pathway enrichment analyses using IPA and gene set enrichment analysis (GSEA) identified disruptions overlapping with the transcriptomic data (cell adhesion, gene expression regulation), and pointed to additional impairments affecting protein translation, metabolism, and mitochondrial function (Suppl. Fig. 10, 11; Suppl. Table 2, Tabs 3, 4). These findings are consistent with transcriptomic signatures indicating perturbations in insulin signalling, including the RAS–MAPK and PI3K–AKT pathways, which regulate cell adhesion, differentiation, transcription, and mitochondrial function^36,37^. Given the observed involvement of PKA and PKC pathways at the transcript level, gene ontology analysis further revealed enrichment of upregulated proteins associated with PKC signalling (Table 1; Supplementary Table 2, Tab 3). In summary, these multi-omics data point to impairments in NSCs which affect cell adhesion, transcription, translation, insulin signalling, and mitochondrial function. Based on these insights from the omics data, we conducted further analyses of AKT and MAPK expression and their phosphorylation, alongside with assessments of mitochondrial function.

**Table 1:** Overview of enriched canonical pathways and biological processes identified in homozygous *KCNQ1*-KO NSCs and neurons by transcriptomic and proteomic analyses.

| *Cell type* | **Transcriptome**<br>*canonical pathways, network enrichment (IPA)* | **Proteome**<br>*canonical pathways (IPA)\* & GSEA#* |
| --- | --- | --- |
| NSCs | Integrin cell surface interaction<br>Generic transcription pathway<br>NCAM signalling for neurite outgrowth<br>Protein kinase A & C signalling<br>Insulin signalling (RAS/ERK, PI3K) | Cell adhesion #<br>Gene expression regulation *,#<br>Protein kinase C signalling #<br>Insulin-like growth factor receptor signalling pathway #<br>Protein translation #<br>Mitochondria function *,#<br>Metabolism * |
| neurons | Generic transcription pathway<br>Insulin signalling - ERK - AKT | Regulation of transcription #<br>Regulation of insulin secretion #<br>Insulin responsive compartment #<br>Vesicle trafficking/recycling (GARP complex)#<br>Protein translation *,#<br>Mitochondria function #<br>Layer formation in the cerebral cortex #<br>Cellular growth, proliferation and development *<br>Cell cycle #<br>Metabolism of RNA and proteins * |

#### Loss of KCNQ1 affects expression of AKT and MAPK in NSCs

Our previous results revealed reduced neurite outgrowth in both heterozygous and homozygous *KCNQ1*-KO NSCs, and enrichment analysis of transcriptomic and proteomic data pointed towards involvement of insulin signalling in homKO 1 NSCs (Table 1), comprising key molecules in downstream signalling cascades, such as AKT and MAPK. Therefore, we followed up if these key regulatory proteins were affected by KCNQ1 loss, by analysing their expression with western blotting. Interestingly, an increased AKT expression was observed for homKO 1 and hetKO 1 compared to their respective controls (Fig. 3f), whereas phosphorylated AKT (Ser473) could not be detected (data not shown). Furthermore, we found increased MAPK expression in the homKO 1 NSCs, but equal phosphorylation levels between the cell lines (Thr202/Tyr204, Fig. 3g, h), indicating increased levels of active MAPK in the homKO 1 line. Taken together, these results show elevated AKT and MAPK signalling in homozygous *KCNQ1*-KO NSCs.

#### Loss of KCNQ1 impairs mitochondrial biogenesis in NSCs

To follow-up potential impairments in mitochondrial function as indicated by the proteomics data analysis, we analysed mitochondrial DNA copy number and deletion. We quantified two genes encoded on the mitochondrial DNA (mtDNA), NADH dehydrogenase 2 (*ND2*) and cytochrome C oxidase 1 (*CO1*), in comparison to the nuclear beta-2-microglobulin gene (*β2M*), to determine the mtDNA copy number. Furthermore, *ND2* and *CO1* were analysed in relation to the mtDNA encoded gene NADH dehydrogenase 4 (*ND4*). The *ND4* gene region is frequently deleted in mitochondrial and other diseases^38^, and therefore can be used to determine mtDNA deletion. In our case, no evidence for a mitochondrial DNA deletion was obtained (Suppl. Fig. 12 b). However, mitochondria copy number was significantly decreased in homKO 1 and hetKO 1 NSCs compared to the respective controls (Fig. 4a), suggesting impaired mitochondrial biogenesis. The proteome analysis revealed mitochondrial dysfunction, affecting different biological processes such as cytochrome c release (GO00018), mitochondrial fusion (GO00080) (GSEA, Suppl. Table 2 (Tab 3)), respiratory electron transport and mitochondrial biogenesis (IPA, Suppl. Fig. 11, Suppl. Table 2 (Tab 4)), among others. In follow-up experiments, we further quantified the mitochondrial protein ATP5α by western blotting. ATP5α is a subunit of the complex V at the inner mitochondrial membrane, the so-called ATP synthase, that produces ATP at the end of the electron transport chain in the process of oxidative phosphorylation. We found lower expression levels of ATP5α in the homKO 1 NSCs compared to WT control (Fig. 4b). In summary, these findings show reduced mitochondria biogenesis, and additionally in homKO 1 a reduced ATP synthase expression.

**Figure 4:**
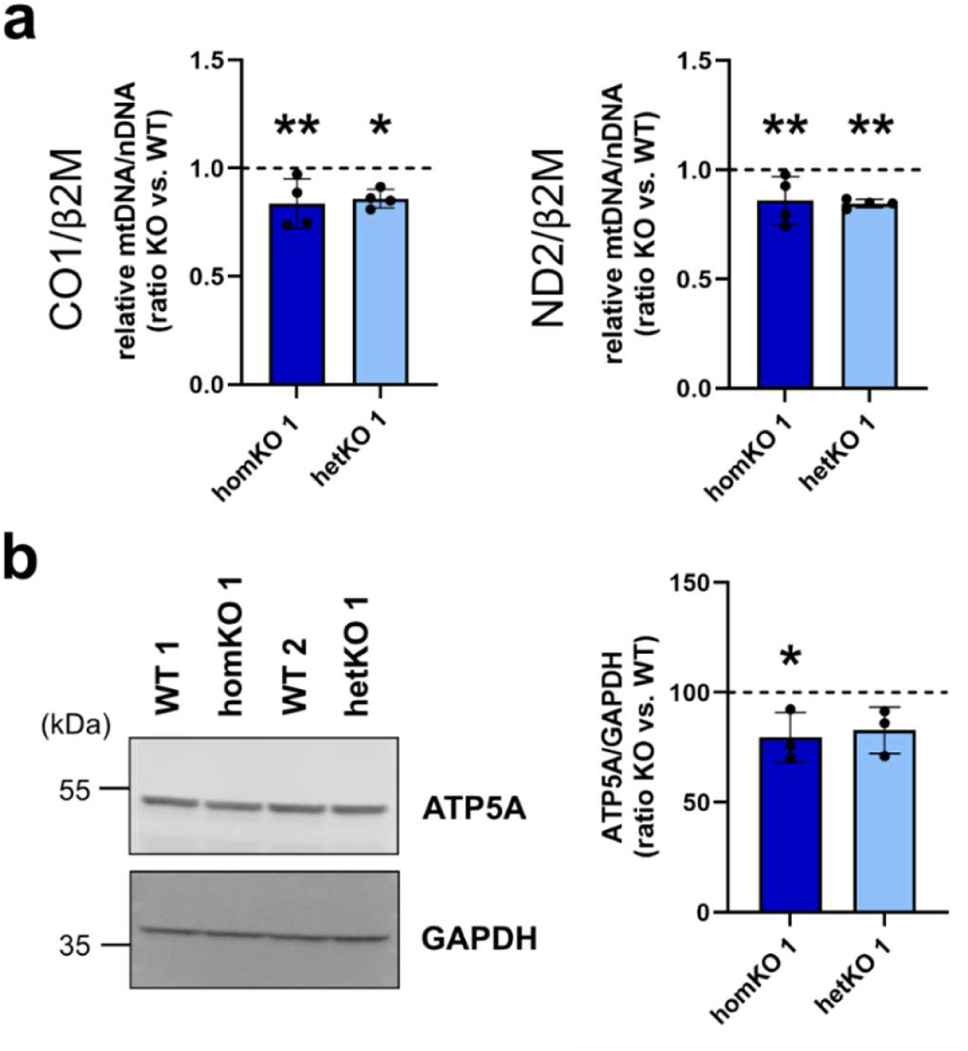
Analysis of mitochondrial copy number and ATP5A expression. **a** The amount of mitochondrial DNA was determined by quantification of two mitochondrial genes (*ND2*, *CO1*) normalized to the amount of DNA of the nuclear gene *β2M*. One-way ANOVA with Bonferroni-adjusted post hoc test, mean ± SD, n=4. *CO1/β2M*: *P*=0.0028, F=8.43, post hoc test: homKO 1: *P*=0.0051, hetKO 1: *P*=0.0134; *ND2/β2M*: *P*=0.002, F=9.195, post hoc test: homKO 1: *P*=0.0082, hetKO 1: *P*=0.0044. **b** ATP5A expression was measured by western blot (left: representative membrane image, right: quantification). One-way ANOVA with Bonferroni-adjusted post hoc test, mean ± SD, n=3, *P*=0.0198, F=5.929, post hoc test: homKO 1: *P*=0.0245, hetKO 1: *P*=0.0542. KO values are normalized to their respective WT control, which are set to the value 1 or 100 and indicated by the dotted line. *ND2*: *NADH dehydrogenase 2*; *CO1*: *cytochrome C oxidase 1*; *β2M*: *beta-2 microglobulin*; ATP5A: ATP synthase F1 subunit alpha.

### The role of KCNQ1 in neurons

#### Loss of KCNQ1 led to reduced neuronal maturity with reduced synaptic activity

To investigate the role of KCNQ1 in cortical neurons, we analysed genes that are typically expressed in neurons and their precursors (Fig. 5a). We focused on the homozygous KO line in comparison to WT 1, as we expected the strongest effects. We discovered a significant reduction of *SYP* (coding for synaptophysin located at the pre-synapsis), *DLG4* (coding for PSD95 located at the post-synapsis), *CUX2* (coding for cut-like homeobox transcription factor 2, expressed in cortical layer II-IV), and the *IR* (coding for the insulin receptor), as well as an increase of *RELN* (coding for reelin, expressed in cortical layer I during early neuronal differentiation) in the homKO 1 compared to WT 1 neurons (Fig. 5a). The increase of *RELN* together with the decrease in *CUX2* suggests that the homKO 1 neurons exhibit delayed neuronal maturation compared to WT 1 neurons, which could be influenced by reduced insulin receptor expression, as proper insulin signalling is needed for neuronal survival and differentiation.

**Figure 5:**
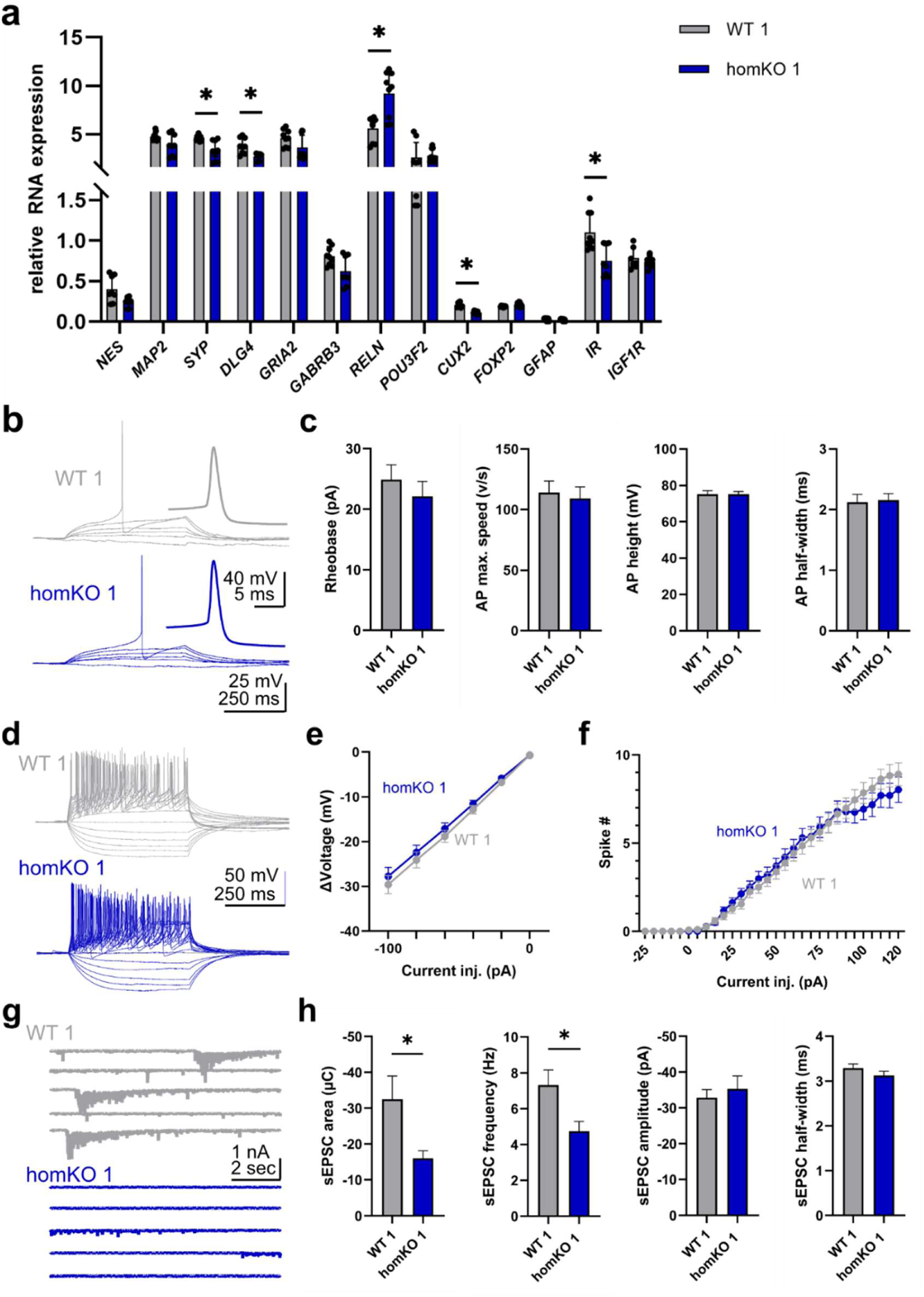
Impact of *KCNQ1* deletion on intrinsic properties and synaptic activity in neurons. **a** Expression analysis of several marker genes in the homozygous KO neurons compared to the isogenic control. Multiple unpaired two-tailed Student’s *t*-test, followed by Holm-Sídák-adjusted post hoc comparison, mean ± SD, n=3. Adjusted *P*-values: *NES*: *P*=0.159609; *MAP2*: *P*=0.186771; *SYP*: *P*=0.013009; *DLG4*: *P*=0.015635; *GRIA2*: *P*=0.272447; *GABRB3*: *P*=0.158609; *RELN*: *P*=0.010376; *POUF3F2*: *P*=0.773693; *CUX2*: *P*=0.000201; *FOXP2*: *P*=0.158609; *GFAP*: *P*=0.773693; *IR*: *P*=0.038275; *IGF1R*: *P*=0.773693. **b, c** Basic properties of neurons. **b** Representative recordings of a control (WT 1) and a mutant (homKO 1) neuron. Cells were kept at near −70mV holding potentials and progressively depolarized using positive current injections (from 0 pA, +5 pA steps for 500 ms). **c** Summary plots of rheobase, as well as action potential (AP) max speed, height, and half-width. Number of experiments: Rheobase WT 1 and homKO 1: 55 cells/3 batches, AP properties (all graphs) WT 1: 43 cells/3 batches, homKO 1: 46 cells/3 batches. Unpaired two-tailed Student’s *t*-test, mean ± SEM. Rheobase: *P*=0.4239; AP max. speed: *P*=0.7198; AP height: *P*=0.9943; AP half-width: *P*=0.8290. **d, e** Neuronal excitability. **d** Representative recordings of a control (WT 1) and a mutant (homKO 1) neuron. Cells were kept at near −70mV holding potentials and subjected to a series of current injection steps (from −100 to +480 pA, 20 pA steps for 500 ms). **e** Summary plot of voltage responses upon injections of hyperpolarizing current injections (from −100 pA to 0 pA). Number of experiments for WT 1 and homKO 1: 45 cells/3 batches for each. Two-way ANOVA (injected current x cell line) with Bonferroni-adjusted post hoc test, mean ± SEM, all *P*>0.05. **f** Summary plot of spiking activity upon injections of depolarizing current injections (from 0 pA to 480 pA). Number of experiments: WT 1 45 cells/3 batches; homKO 1: 44 cells/3 batches. Two-way ANOVA (injected current x cell line) with Bonferroni-adjusted post hoc test, mean ± SEM, all *P*>0.05. **g, h** Spontaneous excitatory postsynaptic currents (sEPSCs). **g** Representative recordings of a control and a mutant neuron at −70 mV holding potentials. **h** Summary graphs of sEPSC integrated charge, sEPSC frequency, amplitude, and half-width. Number of experiments: WT 1 and homKO 1 45 cells/3 batches for each. Unpaired two-tailed Student’s *t*-test, mean ± SEM. sEPSC charge: *P*=0.0183; sEPSC frequency: 0.0123; sEPSC amplitude: *P*=0.5617; sEPSC half-width: *P*=0.2023. *NES*: Nestin; *MAP2*: microtubule associated protein2; *SYP*: synaptophysin; *DLG4*: discs large MAGUK scaffold protein 4; *GRIA4*: glutamate ionotropic receptor AMPA type subunit 4; *GABRB3*: gamma-aminobutyric acid type A receptor subunit beta 3; *RELN*: reelin; *POU3F2*: POU class 3 homeobox 2; *CUX2*: cut like homeobox 2; *FOXP2*: forkhead box P2; *GFAP*: glia fibrillary acidic protein; *IR*: insulin receptor; *IGF1R*: insulin like growth factor 1 receptor.

We additionally assessed the properties and function of the homozygous *KCNQ1*-KO neurons compared to the WT control, using patch-clamp. No differences in basic AP properties (Fig. 5b, c) and neuronal excitability (Fig. 5d-f) were found. However, we observed an overall reduction in excitatory transmission: both the overall charge transfer, as well as the frequency of spontaneous EPSCs were reduced in the homozygous KO neurons compared to WT (Fig. 5g, h). Heterozygous *KCNQ1*-KO neurons (hetKO 1) did not show these alterations, indicating a milder effect on neuronal function (Suppl. Fig. 13a), which could be explained by residual *KCNQ1* expression of the WT allele due to leaky imprinting in neurons (Suppl. Fig. 6a). These data indicate that homozygous loss of *KCNQ1* very likely results in an overall reduced excitatory synaptic function, which could be explained by the fact that they were less mature neurons.

#### KCNQ1 impacts on multiple signalling pathways in neurons

Transcriptomic and proteomic analyses were performed as previously done for the NSCs. RNA sequencing was performed on iPSC-derived neurons from WT 1, WT 2, homKO 1, hetKO 1, hetKO 2, and hetKO 3, each differentiated in four independent rounds from NSCs^31^, yielding four biological replicates per cell line. A total of 15,231 protein-coding genes were detected for all samples, and differential expression analysis was conducted using DESeq2. The PCA plot did not reveal a clear separation between the WT and the KO groups, suggesting minimal differences in the underlying data, and a low impact of the *KCNQ1*-KO on transcription in neurons (Suppl. Fig. 13b). This is in line with the fact that potassium channels mostly influence transcription indirectly after their activation, whereas our analysis was carried out under basic cellular conditions. However, we identified protein coding DEGs using an FDR threshold of < 0.05 (Suppl. Fig. 13c, 14a, Supplementary Table 1 (Tabs 3–6)). As for the NSCs, the processed and raw (FASTQ) gene expression data for the neurons is available through the gene expression omnibus (GSE number to be allocated). Overlapping DEGs in at least three *KCNQ1*-KO cell lines were found with a function in mitochondria (*TXNRD2*), transcriptional regulation (*ZFP3, ZNF471, ZNF585A/B, HOXB9*), cell adhesion (*PCDHA4, PCDHGC3*), neurodevelopment and neuronal survival (*NGFR, NHLH2*), regulation of insulin (*ISL1*), and t-RNA-guanine transglycosylation (*C9orf64*) (Suppl. Fig. 14 a-c, Suppl. Table 1 (Tab 7)). Validation using nCounter analysis confirmed differential expression for 14 of 16 selected genes (Suppl. Fig. 14d) across almost all cell lines (Suppl. Table 1 (Tab 8)), demonstrating high reproducibility of the RNA sequencing data — even for genes with modest effect sizes (e.g., *NHLH2* in hetKO 3: log₂ fold change = 0.636).

IPA identified as an enriched canonical pathway in the homKO 1 neurons the reduction of the generic transcription pathway, caused by a high number of differentially expressed transcriptional regulators. Network enrichment analysis pointed towards an involvement in insulin signalling for homKO 1, hetKO 1, and hetKO 2 neurons, including dysregulation of both downstream effectors AKT and MAPK/ERK (Table 1, Suppl. Table 1 (Tabs 9-12), Suppl. Fig. 15, 16).

We also performed quantitative proteomic analysis by LC mass spectrometry on iPSC-derived neurons from WT 1, WT 2, homKO 1, hetKO 1, hetKO 2, and hetKO 3 lines, differentiated in parallel to the RNA-seq samples across four independent rounds from NSCs to neurons^31^, yielding four biological replicates per genotype. In total, 2,990 proteins were identified. In contrast to the transcriptomic profiles, PCA separated homKO 1, hetKO 1, and hetKO 2 neurons from WT 1 and WT 2, and correlation analysis revealed clustering by genotype (Suppl. Fig. 17), indicating distinct protein expression signatures associated with KCNQ1 loss. Using an FDR cutoff of < 0.10, we identified significantly altered neuronal proteins (Fig. 6a, b; Suppl. Table 2).

**Figure 6:**
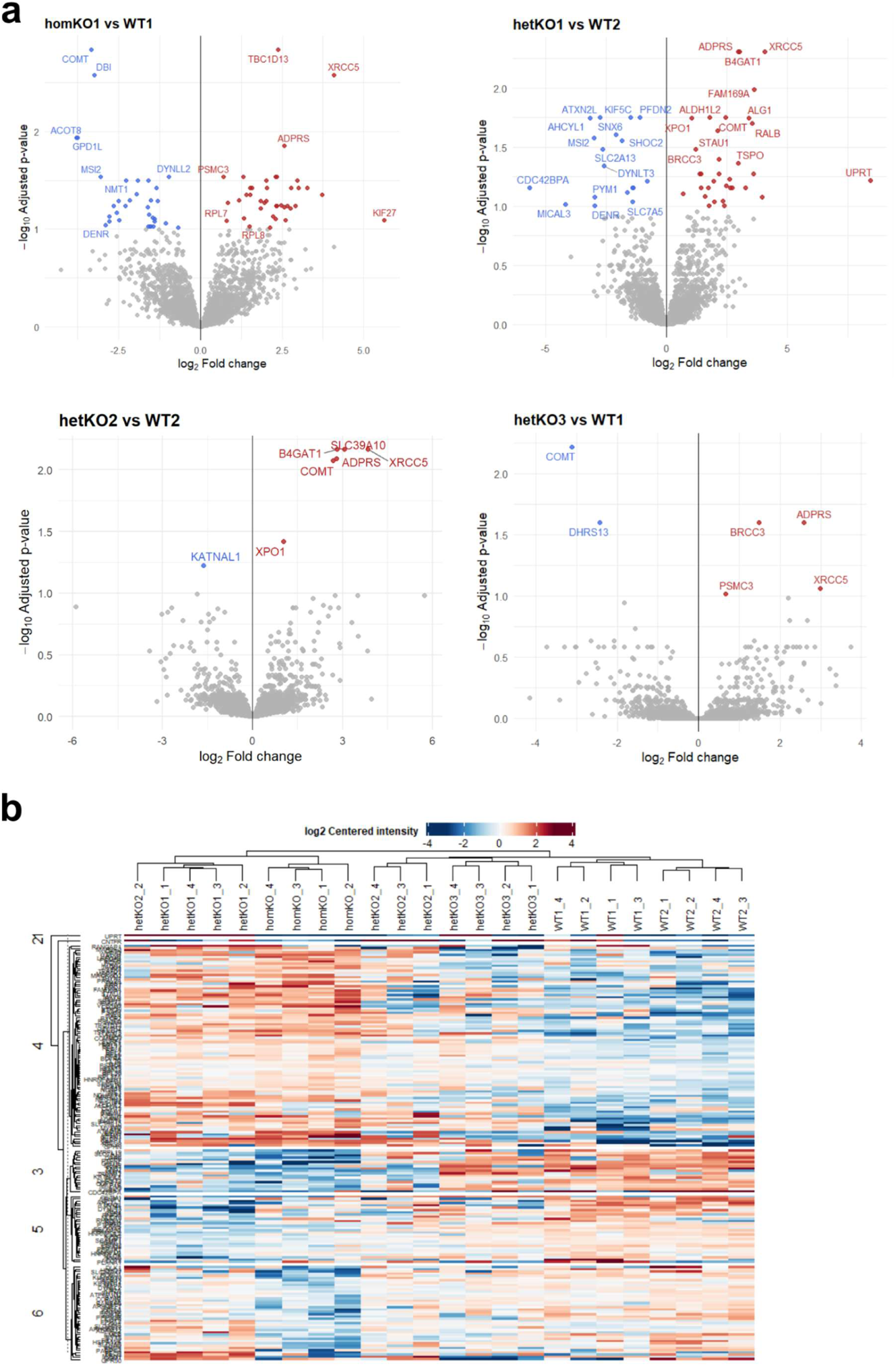
Proteomic data of *KCNQ1*-KO neurons. **a** Volcano plots showing DEPs in neurons (FDR < 0.1, log2fold change > 1.3). Significantly upregulated proteins are shown in red, downregulated proteins in blue. **b** The heatmap shows the log2 centered intensity of all DEPs in homKO 1. The dendrogram indicates separation between WT and KO cell lines, where homKO 1 and hetKO 1 as well as hetKO 2 and hetKO 3 are clustering closer together.

For homKO 1 neurons we identified 29 DEPs, 11 downregulated and 18 upregulated. Among the most significant changes, seven mitochondrial proteins with distinct metabolic functions were differentially expressed in homKO 1 neurons (|log₂FC| > 1.9). ACOT8, GPD1L, DBI, and DHRS13 were downregulated, whereas ATAD3B, MAVS, and TSPO were upregulated, pointing to impairments in mitochondrial lipid metabolism, redox homeostasis and signalling. Beyond metabolism, several proteins involved in RAS–MAPK and PI3K–AKT signalling (KRAS, STRN3, MSI2, ROBO1, NMT1, TBC1D13; |log₂FC| > 1.3) were affected, implicating perturbations in pathways governing cell growth and survival. Additionally, proteins linked to mTOR-dependent translation control (PAIP1, MSI2, RPS2, STAU1, TBC1D13; |log₂FC| > 1.4) were found, suggesting broader effects on protein synthesis and translational regulation. Three of these proteins were also differentially expressed in hetKO 1 and hetKO 2 neurons: TSPO (translocator protein), MSI2 (musashi RNA-binding protein 2), STAU1 (staufen double-stranded RNA-binding protein 1). TSPO and STAU1 were upregulated, whereas MSI2 was downregulated. TSPO is a mitochondrial protein involved in mitochondrial signalling, metabolism, and cell survival. As an RNA-binding protein, STAU1 plays a major role in mRNA transport, localization, and decay in neurons. MSI2 is a post-transcriptional regulator involved in neuronal differentiation.

In addition, enrichment analysis with IPA and GSEA of the homKO 1 neurons pointed to impairments in cell growth, proliferation, and development (Suppl. Fig. 11), supporting our previous results of delayed neuronal maturation. Both analyses further predicted that KCNQ1 impacts on protein translation (Table 1, Suppl. Fig. 11, 18).

As the transcriptomic data analysis revealed an involvement of mRNA transcription and insulin signalling, we searched for additional evidence at the protein level in the GSEA gene ontology (GO) terms (Suppl. Table 2 (Tab 5)) and in the IPA canonical pathways (Suppl. Table 2 (Tab 6)). The GSEA proteome data supported an involvement of these processes. Table 1 summarizes the identified pathways comparing transcriptomic and proteomic data analysis results.

The omics results should be interpreted as exploratory and hypothesis-generating, warranting further experimental validation. Based on the analysis of omics data, the observed alterations in AKT/MAPK signalling and mitochondrial function were expected to persist in KO neurons. However, follow-up experiments using qPCR and western blotting did not provide additional evidence for defects in mitochondrial biogenesis or ATP synthase expression in neurons (mtDNA copy number, ATP5A expression, Suppl. Fig. 12b, c).

#### KCNQ1 affects similar pathways in NSCs and neurons

Several impairments identified by transcriptomic analysis in homozygous *KCNQ1*-KO NSCs persist in the differentiated neurons, particularly in gene transcription and insulin signalling pathways, including AKT and MAPK signal transduction. At the protein level, GSEA and IPA revealed overlaps between NSCs and neurons, with affected signalling pathways in both cell types involving protein translation, gene transcription, mitochondrial function, and cellular metabolism (Table 1). Comparative GSEA proteomic analysis of NSCs and neurons revealed the ATPase complex and mitochondrial protein-containing complexes as among the most significantly affected cellular compartments in both cell types. Accordingly, we focused on ATP- and mitochondria-related GO terms and identified several enriched mitochondrial processes, including mitochondrial transport, the inner mitochondrial membrane, and cytochrome c release. In addition, multiple ATP-related biological processes were consistently enriched, indicating impaired ATPase function which affects ATP hydrolysis and proton transmembrane transport. These findings indicate a clear impact of KCNQ1 loss on energy metabolism in both NSCs and neurons (Suppl. Fig. 19). Overall, these consistencies strengthen our findings and support the robustness of the observed effect.

## Discussion

To investigate the role of KCNQ1 in human neuronal development, we generated *KCNQ1*-KO and isogenic control iPSC lines, which were subsequently differentiated into NSCs and cortical neurons. We observed that KCNQ1 already plays a role in early neuronal differentiation, influencing neurite outgrowth, mitochondrial function, and cellular energy metabolism. In differentiated neurons, *KCNQ1* loss resulted in reduced functional activity, and a more immature gene expression profile. Combined transcriptomic and proteomic analyses revealed that key biological processes — including gene transcription, protein translation, mitochondrial function, and insulin signalling — were disrupted in both NSCs and neurons. These results indicate that molecular dysfunctions originating during early neurodevelopment persist throughout neuronal maturation.

In NSCs, both homozygous and heterozygous *KCNQ1*-KOs showed reduced neurite outgrowth during early neuronal differentiation. Moreover, pharmacological inhibition of KCNQ1 channels with the antagonist JNJ303 resulted in reduced neurite outgrowth in both WT and hetKO 1 NSCs, whereas no effect was observed in homKO 1 NSCs. Notably, the reduced neurite outgrowth in the hetKO 1 NSCs could be rescued by activating the Kv7.1 channel with the agonist ML277. Even though *KCNQ1* is an imprinted gene and the deletion in hetKO 1 affects the active allele, this NSC line also expressed the WT allele which explains the rescue effect. As a possible underlying mechanism for the observed diminished neurite outgrowth, transcriptome and proteome analyses pointed towards impairments in cell adhesion and a decrease in NCAM signalling, a known regulator of neurite outgrowth. Additionally, the omics data revealed that loss of *KCNQ1* affects PKA and PKC signalling pathways in NSCs. While transcriptomic data predicted an inhibition of both PKA and PKC signalling, the PKC signalling seems to be activated at the protein level (GSEA), suggesting the involvement of post-transcriptional regulation or compensatory responses. Importantly, both PKA and PKC modulate KCNQ1 activity through phosphorylation and are known to regulate key cellular processes such as adhesion, proliferation, differentiation, and metabolism^34,35,39^. Therefore, dysregulation of these pathways may also contribute to the observed cellular phenotype.

The loss of *KCNQ1* in heterozygous and homozygous KO NSCs results in mitochondrial dysfunction, characterized by a decrease in mitochondrial biogenesis — as evidenced by reduced mitochondrial DNA copy numbers. Moreover, the expression of a subunit of the ATP-synthase complex (ATP5α), a key component in ATP production, is diminished, suggesting decreased ATP production and a potentially compromised cellular energy metabolism. The proteomic data analysis further supported these findings, indicating impairments of several mitochondrial functional aspects and cell metabolism. During the differentiation of NSCs into neural progenitor cells, and eventually neurons, cells undergo an energetic transition from the dominance of glycolysis to relying on mitochondrial oxidative phosphorylation (OxPhos) — a more efficient mode of ATP generation^40^. As the process of neuronal differentiation increases energy demands, an impaired mitochondrial energy supply very likely explains the observed reduction in neurite outgrowth in differentiating NSCs lacking *KCNQ1*. In these KO cells, the metabolic shift from glycolysis to OxPhos may be delayed. Moreover, a possible ATP deficiency could affect phosphorylation of KCNQ1 directly and indirectly through PKC signalling^41^. Taken together, these results suggest that KCNQ1 plays an important role in regulating cellular energy metabolism.

Further, loss of *KCNQ1* in NSCs and neurons disrupts insulin signalling. This was initially observed in the enrichment analysis of the DEGs and DEPs in the homKO 1 dataset, which indicated a dysregulation of the insulin signalling, including downstream the RAS-MAPK and PI3K-AKT cascades. Furthermore, gene expression of the insulin receptor was significantly reduced in homKO 1 neurons relative to WT controls. These neurons showed a general enrichment of proteins regulated by insulin (insulin-responsive compartment). In particular, pathways related to insulin secretion and insulin receptor signalling were enriched, with additional evidence supporting the involvement of AKT and MAPK signalling. AKT and MAPK, both regulate cytoskeleton dynamics and promote neurite outgrowth^42^. As previously reported, both pathways respond to a wide range of stimuli, including growth factors, cytokines, nutrient levels, and cellular stress, enabling them to tightly regulate cellular energy homeostasis and mitochondrial function. Remarkably, AKT and MAPK signalling promote mitochondrial biogenesis, enhance mitochondrial fusion, and support OxPhos and ATP production^36,37^. Therefore, the upregulation of AKT and MAPK observed in *KCNQ1*-deficient NSCs by western blotting likely reflects a compensatory mechanism aimed at restoring mitochondrial biogenesis and function, which were found to be impaired.

KCNQ1 also interferes with the mode of action of the anti-diabetic drug metformin, likely through its effect on insulin signalling. Loss of *KCNQ1* impaired the response of NSCs to metformin in terms of neurite outgrowth. Metformin is known to modulate insulin signalling by activating AMPK, inhibiting mTOR, and directly inhibiting mitochondrial electron transport chain complex I^33^. Several studies have reported that metformin enhances proliferation and promotes neurite growth of NSCs^43,44^, which contrasts with our findings. In this study, metformin had no effect on NSC proliferation and instead reduced neurite outgrowth in WT and hetKO 1 cell lines, compared to their respective mock treatments. This reduction, however, was not observed in homozygous *KCNQ1*-KO NSCs, suggesting that the effect of metformin on neurite outgrowth depends on functional KCNQ1. Potential toxic effects of metformin were ruled out, as apoptosis rates were not elevated. The discrepancies with previous studies may be due to differences in experimental design and timeframes, such as acute treatment with a low dose of metformin (0.5 mM), and short-term observation over 48 hours.

In recent years metformin has received increasing attention for its potential neuroprotective and cognitive-enhancing properties in the context of intellectual disability (ID) and major neurodegenerative disorders like Alzheimeŕs disease^45^. Emerging evidence from both preclinical models and early-phase clinical studies has highlighted its potential therapeutic benefits in specific forms of ID, such as Fragile X syndrome^46,47^. As our findings indicate that KCNQ1 interferes with the pharmacological activity of metformin in NSCs, this may have relevance for individuals harboring deleterious genetic variants in *KCNQ1*, as such variants could impair metformin responsiveness and potentially diminish its cognitive-enhancing effects.

We discovered that KCNQ1 plays a role in neuronal function, and its loss led to a reduced frequency of spontaneous EPSC. This reduction is likely due to the delayed maturation of homKO 1 compared to WT neurons. HomKO 1 neurons exhibit lower expression of synaptic genes such as *SYP* and *DLG4*, increased expression of the cortical layer I marker *RELN*, and decreased expression of the layer II–IV marker *CUX2*. Additional evidence for impairments of neuronal differentiation was obtained at protein level pointing towards impairments in the layer formation of the cerebral cortex (GSEA) and cell proliferation and growth (IPA). Insulin signalling is critically important for early neuronal development both *in vivo* and *in vitro*, as it regulates NSC proliferation, neuronal differentiation, axonal and dendritic growth, synaptogenesis, and neuronal survival^48^. Thus, the combination of decreased insulin receptor expression and disrupted cellular energy metabolism very likely contributes to the reduced maturity phenotype of homKO 1 neurons and their reduced sEPSC frequency.

Taken together, this study is the first to demonstrate a functional role of KCNQ1 in NSCs and during neuronal differentiation. Our findings show that KCNQ1 influences key cellular processes, including cell adhesion, transcriptional regulation, insulin signalling, and mitochondrial function. Moreover, KCNQ1 affects neuronal activity and differentiation, likely through the modulation of insulin signalling pathways. Clinically, *KCNQ1* variants are linked to disorders of peripheral organs — such as long/short QT syndrome, atrial fibrillation, Jervell and Lange-Nielsen syndrome, Romano-Ward syndrome, and type 2 diabetes (DM2) — as well as neurological conditions including epilepsy and OCD ^8–18^. Some genetic variants were even directly linked to comorbidities, such as long QT syndrome with epilepsy. Thus, elucidating the role of KCNQ1 in human neuronal cells contributes to a better understanding of these disease comorbidities. Individuals with pathogenic *KCNQ1* variants, while initially diagnosed with cardiac disease, may also have an elevated risk for DM2, epilepsy, cognitive impairment, and psychiatric disorders^19,20^.

While this study provides important insights, some limitations should be taken into account when interpreting the results. Although many results reached statistical significance and were reproducible — for instance, the observed reduction in neurite outgrowth during NSC differentiation — the identified effect sizes were modest. This may be attributable to the inherently low expression levels of KCNQ1 in NSCs and neurons. The sample size was limited, as only one homozygous *KCNQ1*-KO iPSC clone was available, and most functional analyses in NSCs were conducted using this clone alongside a single heterozygous KO line and their respective isogenic controls. Different significance thresholds needed to be applied for transcriptomic and proteomic analyses. While transcriptomic data were evaluated at FDR < 0.05, a less stringent threshold (FDR < 0.10) was used for proteomic data due to the limited number of differentially expressed proteins. Further investigation will provide valuable insights into the extent of insulin signalling and mitochondrial dysfunction in neurons with KCNQ1 loss and deepen our understanding of the associated metabolic and energetic consequences.

Studying patient-specific *KCNQ1* variants in iPSC-derived cardiomyocytes, NSCs and neurons, holds promise for identifying common mechanisms of comorbidity and assessing individual patient risk. A comparative analysis of our findings in NSCs and neurons with mechanisms observed in cardiac cells will help to determine whether shared or distinct cellular impairments underlie the observed comorbidities.

Notably, our data shows that KCNQ1 has a strong effect on several molecular pathways in neurons and their progenitors — particularly those involving insulin signalling and mitochondrial function — which have also been implicated in the development of insulin resistance and related disease phenotypes^49^. Potassium homeostasis and insulin signalling are intricately connected influencing neuronal excitability and synaptic function^50^. Previous studies showed that dysregulation in brain K⁺ channels impairs neuronal insulin sensitivity, potentially contributing to disorders such as obesity and DM2^51^. Studies in mice even revealed that blocking of voltage-gated potassium channels ameliorated diabetic cognitive dysfunction^50^. Therefore, targeting potassium channels, including the Kv7.1 channel, may be a promising strategy for the treatment of comorbid cognitive impairments and diabetes.

## Materials and Methods

### CRISPR/Cas9 editing

Guide RNAs (gRNAs) flanking the exons 4 and 5 of *KCNQ1* (hg38, MANE transcript NM_000218.3) were designed using the CRISPR/Cas9 target online predictor CCTop^52^. Two gRNAs were used in combination to delete both exons (two different gRNA combinations, gRNA 1 + gRNA 2, gRNA 3 + gRNA 4) (Fig. 1a). Further details on the gRNAs are provided in the Supplementary materials and methods. To generate the KO cell lines, iPSCs were split with Accutase. TrueCut Cas9 protein v2 (A36497, Thermo Scientific) and gRNAs were introduced into the cells by nucleofection. Subsequently, per reaction 200.000 cells were seeded in a 24-well. When cells reached confluency, single cell cloning was performed by seeding 500 or 1000 cells on a 6 cm dish. Colonies were picked after 7 days and placed into a 96-well plate. After 5-7 days colonies reached 70-80% confluency and were further expanded. During this split, part of the cells was used for DNA isolation and KO-screening by PCR. One homozygous iPSC line (homKO 1) and three heterozygous KO clones (hetKO 1, hetKO 2, hetKO 3) were selected. Two unedited lines were used as isogenic WT controls (WT 1, WT 2). The KO was confirmed by PCR across the deleted region followed by Sanger sequencing to determine the exact position and size of the deletion (gRNA 1 + gRNA 2: 960 bp, gRNA 3 + gRNA 4: 1207 / 1219 bp) (Suppl. Fig. 1). Two heterozygous clones carry the deletion on the active allele (hetKO 1, hetKO 2), and one cell line on the inactive allele (hetKO 3) (Suppl. Fig. 1). Cell lines generated with the same gRNA were compared in the follow-up experiments (Fig. 1b).

### NSC induction

NSCs were generated by using the protocol of Yan et al.^31^. IPSCs were split with Accutase and 300.000 cells/6-well were seeded in mTeSR Plus medium. 24h after the split, medium was changed to PSC Neural Induction medium (A1647801, Thermo Fisher, 98% neurobasal medium, 2% neural induction supplement (NIS)). On day 7 of induction, cells were split with a high cell density (3 million cells/6-well) in neural expansion medium (NEM) (Neurobasal, Advanced DMEM/F12 (12634010, Thermo Fisher), PenStrep (15140122, Thermo Fisher), NIS) and ROCK inhibitor for 24h. After this split, these induced NSCs at passage 1 (P1) were further maintained in NEM up to P10 with minimal spontaneous differentiation (Suppl. Fig. 4a). Whenever cells reached 100% confluency (every 4-5 days), they were split with Accutase. ROCK inhibitor was used from P1 to P4. From P5 till P10 onwards cells were used for experiments.

### Neuronal differentiation

Neurons were generated by using the protocol of Yan et al.^31^ with some modifications. For differentiation, NSCs were first maturated for 6 days and then differentiated into cortical neurons for 6 weeks (Suppl. Fig. 5a, b). For the maturation, NSCs were seeded in 6-well format on Geltrex coating in NEM medium. After 24h, medium was changed to maturation medium (MM) (50% Neurobasal medium, 50% Advanced DMEM/F12, 1% Glutamax (35050061, Thermo Fisher), 1% N2 (17502048, Thermo Fisher), 0.5% B27 (17504044, Thermo Fisher) and 1% PenStrep) and after 6 days, cells were seeded for neuronal differentiation on Ornithin-(P4957, Sigma Aldrich) Laminin (L2020, Sigma Aldrich) coating, either in 6-well format or 24-well format with cover slips, depending on the follow-up experiment. Again, 24h after the split, medium was changed to neuronal differentiation medium (NDM) containing Neurobasal Plus, 2% B-27 plus (both A3653401, Thermo Fisher), 1% Glutamax, 0.1% ascorbic acid (A5960, Sigma Aldrich) and 1% PenStrep. We performed a MitomycinC (M0503, Sigma Aldrich) treatment when cells reached 90-100% confluency (usually on day 6 of differentiation, 10 µg MitomycinC for 10 min). Cells were always differentiated for 42 days in NDM and then used for analysis. This protocol was used for all experiments including NSCs and neurons.

An alternative protocol for neuronal differentiation was used to confirm a role of KCNQ1 in neurite outgrowth which was modified according to Qi et al^32^. Unlike the protocol from Yan et al.^31^, which involved the generation and expansion of neural stem cells (NSCs) prior to neuronal differentiation, the Qi et al. method induces rapid cortical neuron differentiation directly from iPSCs using small-molecule inhibitors, bypassing the NSC stage. Cells differentiated via the Qi protocol were assessed for neurite outgrowth after splitting on day 8 of differentiation. Further details are given in the Supplementary information.

### Neurite outgrowth and treatments

To measure neurite outgrowth, 120.000 NSCs were seeded in a 24-well on Ornithin-Laminin coating in NDM and immediately placed into the IncuCyte S3 Live-cell Analysis Instrument system (Sartorius). In case of treatments, NDM contained either JNJ303 (1 µM; 3899, Tocris), ML277 (1 µM or 2 µM; 4777, Tocris), or metformin (0.5 mM; D150959, Sigma Aldrich). For the mock control treatment, DMSO was used, as both JNJ303 and ML277 were diluted with DMSO. Water was used as mock control for metformin treatment, as it was diluted in water. One hour after cell seeding, the first measurement was conducted. Cells were afterwards imaged every 4 hours for 48 hours. Neurite length was determined with the Neurotrack software and normalized to cell body cluster area. Each measured time point was normalized to the first measurement (time point 0) which was set to 1. When comparing the KO to its respective WT control, each time point of the KO line was normalized to the respective time point of the WT control, which was also set to the value 1. The same procedure was used for treatment experiments, were mock treatment of each cell lines served as control and was set to 1, and treatment conditions were analysed in relation to mock treatment of the same cell line.

### Western blotting

Protein was isolated with Pierce RIPA Buffer (89900, Thermo Scientific) containing protease inhibitor cocktail (PIC) (S8820, Sigma Aldrich) and PhosSTOP (4906845001, Sigma Aldrich). After incubation for 5 min on ice, benzonase (70746-4, Sigma Aldrich) was added and incubated for 10 min on ice. Afterwards, samples were centrifuged at 13.000 rpm at 4°C for 45 min. The supernatant was transferred to a fresh tube, as the final protein lysate. Protein concentration was determined with a BCA Kit (23225, Thermo Scientific). Samples were denatured at 95°C for 5 min and loaded to a 4-12% Tris-Glycin gel (XP04120BOX, Thermo Scientific). Protein was transferred from the gel to a PVDF membrane with the iBlot 2 system (Invitrogen). Membranes were blocked in blocking buffer (Intercept Blocking Buffer by LI-COR (927-70001) + PBS) for 1h at RT shaking. Primary antibody was incubated in blocking buffer over night at 4°C under constant shaking. On the next day, membranes were washed 3× 10 min in PBS-T. Secondary antibody was incubated for 45 min at RT in blocking buffer and constant shaking, and afterwards removed by washing 3x in PBS-T. Membranes were imaged using the Odyssey Imaging System (LI-COR). All antibodies used are listed in the Supplementary information.

### Electrophysiology

#### General

Electrophysiological recordings were performed using a BX51 upright microscope (Olympus), equipped with differential interference contrast (DIC) and fluorescent capabilities. Electrical signals were recorded using a Multiclamp 700B amplifier (Axon Instruments) controlled by Clampe× 10.1 and Digidata 1440 digitizer (Molecular Devices). Recordings were done at 26 ± 1°C using a dual TC344B temperature control system (Sutter Instruments), with neurons continuously perfused with oxygenated (95% O2 and 5% CO2) artificial cerebro-spinal fluid containing 125 mM NaCl, 2.5 mM KCl, 1 mM MgCl_2_, 2 mM CaCl_2_, 25 mM glucose, 1.25 mM NaH_2_PO_4_, 0.4 mM ascorbic acid, 3 mM myo-inositol, 2 mM Na-pyruvate, and 25 mM NaHCO3, pH 7.4. Neurons were approached and patched under DIC, using 3.0 ± 1.0 MegaOhm pipettes (WPI), pulled with a PC10 puller (Narishige, Japan). For current clamp experiments, pipettes were loaded with an internal solution containing (in mM) 140 K-gluconate, 20 KCl, 10 HEPES, 0.05 BAPTA, 10 ATP-Mg^2+^, 0.3 GTP-Na^+^, 10 Na^+^-Phosphocreatine, osmolarity 312 mOsmol, pH 7.2 adjusted with KOH. For voltage clamp experiments, pipettes were loaded with an internal solution containing 125 mM Cs-gluconate, 20 mM KCl, 4 mM MgATP, 10 mM Na-phosphocreatine, 0.3 mM GTP, 0.5 mM EGTA, 2 mM QX314 (Hello Bio, #HB1030), and 10 mM HEPES, pH 7.2. Data analysis was done using Clampfit 10.5.

#### Intrinsic neuronal properties

To determine the basic electrophysiological properties of control and mutant iPSC-derived neurons, we performed whole cell current clamp recordings. In these experiments, cells were maintained at ∼-70 mV potentials by injecting small (< 30 pA) amounts of current through the recording pipette. Cells in which larger current injections were needed to keep the membrane potential at ∼-70 mV were discarded. To determine the rheobase, i.e. the minimal current needed to trigger an action potential (AP), cells were increasingly depolarized from their resting potential by injections of positive current (1 pA steps) for 500 ms. The AP properties (height, half-width, max speed) were extracted from spikes generated during rheobase measurements. To determine the input resistance and input-output function of the differentiated neurons, we used stepwise current injections through the recording pipette (500 ms pulses) from-100 to 480 pA, with 20 pA steps.

#### Spontaneous and miniature excitatory postsynaptic currents

To measure excitatory currents, we performed whole-cell voltage-clamp recordings at −70 mV holding potentials. Spontaneous excitatory currents were detected as downward deflections from the baseline. Miniature currents were recorded under identical conditions, but after addition of 0.5 μM tetrodotoxin to the bath, to block sodium channels and thus spike-dependent spontaneous excitatory transmission.

### RNA sequencing and data analysis

RNA isolation for RNA sequencing was carried out by GENEWIZ from Azenta Life Sciences. Frozen cell pellets of NSCs (2.5-3 million cells) from homKO 1 and WT 1 samples and cortical neurons from homKO 1, hetKO 1, hetKO 2, hetKO 3, WT 1, and WT 2 were provided.

#### RNA sequencing

Library preparation with poly(A) selection and 150-bp paired-end sequencing on an Illumina NovaSeq, as well as mapping sequence reads to the reference genome and extracting gene counts were performed by GENEWIZ (Germany). Specifically, sequence reads were trimmed to remove possible adapter sequences and nucleotides with poor quality using Trimmomatic v.0.36^53^. The trimmed reads were mapped to the Homo sapiens GRCm38 reference genome available on ENSEMBL using the STAR aligner v.2.5.2b^54^. Unique gene hit counts were calculated by using featureCounts from the Subread package v.1.5.2^55^. The hit counts were summarized, and only unique reads that fell within exon regions were counted. Furthermore, counts were normalized according to the library size and filtered, keeping only genes having count per million reads (cpm) >0.5 across 4 samples for further analysis, as previously suggested^56^. Exploratory data analysis included principal component analysis (PCA) and correlation analysis to assess variability between samples and hierarchical clustering with heatmap using R package PCAtools^57^. For the differential gene expression analysis, we used R package DESeq2^58^ and performed pairwise comparisons between: (a) NSCs homKO 1 and WT 1 samples, (b) neuronal hetKO 1 and WT samples, (c) neuronal hetKO 2 and WT samples, (d) neuronal hetKO 3 and WT samples, (e) neuronal homKO 1 and WT samples. When testing for differential expression in neuronal cells, we adjusted for the effect of type of gRNA and differentiation batch, as these two factors showed influence on gene expression based on PCA plot Volcano plots and heatmaps were generated using Rstudio with ggplot2 package^59^. Venn diagrams were generated by InteractiVenn^60^.

#### Ingenuity Pathway Analysis (IPA)

We used IPA (Q2-2024) to analyse differentially expressed genes for enriched canonical pathways and networks (with up- and down-regulated genes analysed together). The background gene list included all genes that were detected to be expressed in each of the cell lines. We applied the Benjamini-Hochberg procedure for multiple hypothesis correction and used a *P*-adjusted threshold (FDR) of < 0.05 to denote significant findings.

### Liquid chromatography mass spectrometry (LC-MS)

#### Preparation of beads for SP3 protein digestion

Following vortexing, 80 µl Promega MPSP beads (cat. No CS3325A04) and 720 µl of dH_2_O were combined in a low-binding Eppendorf tube. The tube was placed on a magnetic rack allowing beads to settle for about 2 min. After the supernatant was removed, beads were rinsed three times with 800 µl of dH_2_O. Beads were prepared freshly.

#### Protein reduction, alkylation, and digestion using SP3 method

10 µg of protein extracts were digested using the SP3 method with some modifications^61^. Each protein extract was mixed with reduction/alkylation buffer (100 mM TEAB pH 8.5, 1% SDS, 10 mM TCEP, 40 mM CAA) and incubated at 95°C for 5 min and then 25 min at 70°C. After the samples were cooled down to RT, 2 µl of pre-rinsed beads (see above) were mixed in. Finally, acetonitrile (ACN) was added to reach 80% ACN. The samples were then mixed thoroughly and incubated for 20 min on a shaking incubator. Next, samples were shortly centrifuged, placed on a magnetic rack, and the supernatant was carefully removed. Beads were washed with 80% v/v EtOH for a total of three washes, and once with 80% v/v ACN. Beads were reconstituted in 100 µl of 100 mM TEAB, pH 8.5. Trypsin was added in a ratio of 1:50, followed by overnight incubation at 37°C. Samples were acidified using TFA to reach pH < 2. The supernatant was transferred to fresh vials. Digested proteins were desalted on self-assembled C18 Empore® extraction discs (3M, Maplewood, MN) STAGE tips^62^.

#### LC-MS analysis and data processing

Samples were suspended in 0.1% TFA and an equivalent of 2.5 µg of peptides was analysed using an Ultimate 3000 liquid chromatography system coupled to an Orbitrap QE HF (Thermo Fisher) as described earlier^63^. Mobile phase solutions were prepared as follows, solvent A: 0.1% formic acid / 1% acetonitrile, solvent B: 0.1% formic acid, 89.9% acetonitrile. Shortly, peptides were separated in a 120 min linear gradient started from 3% solvent B and increased to 23% B over 100 min and then to 38% B over 20 min, followed by washout with 95% B. The mass spectrometer was operated in data-dependent acquisition mode, automatically switching between MS and MS2. MS spectra (m/z 400–1600) were acquired in the Orbitrap at 60,000 (m/z 400) resolution, and MS2 spectra were generated for up to 15 precursors with normalized collision energy of 27 and isolation width of 1.4 m/z.

#### LC-MS data analysis

The MS/MS spectra were searched against the H. sapiens (UP000005640) and a customized contaminant databases using Proteome Discoverer 2.5 with Sequest HT^64^. The fragment ion mass tolerance was set to 0.02 Da, and the parent ion mass tolerance to 5 ppm. Trypsin was specified as an enzyme. The following variable modifications were allowed: Oxidation (M), Deamidation (N, Q), Acetylation (N-terminus), Met-loss (M), and a combination of Met-loss and acetylation (N-terminus), whereas Carbamidomethylation (C) was set as a fixed modification. Peptide quantification was done using a precursor ion quantifier node with the Top N Average (n = 3) method set for protein abundance calculation. Data preprocessing was performed in R (v4.3.1). Protein identifiers were made unique using the “make_unique()” function from the DEP2 package^65^. Normalization was performed using the variance stabilizing normalization (VSN) method implemented in DEP2, adapted from the vsn package^66^. Missing values were imputed using the MinProb method (q = 0.01). Visualization was performed using PCA, volcano plots, k-means clustered heatmaps, and correlation matrices. Gene Set Enrichment Analysis (GSEA) was conducted using the clusterProfiler (v4.10.0) package^67–70^. Enrichment was performed with pvalueCutoff = 0.05 and without *P*-value adjustment (pAdjustMethod = “none”) unless otherwise stated. Visualization of enrichment results was carried out using enrichplot (v1.22.0) and DOSE (v3.28.1)^71^.

#### Ingenuity Pathway Analysis (IPA)

We used IPA (Q3-2025) to analyse differentially expressed proteins for enriched canonical pathways (with up- and down-regulated proteins analysed together). The background protein list included all proteins that were detected to be expressed in each of the cell lines. We used a non-adjusted *P* value threshold of < 0.05 to denote significant findings (for differentially expressed proteins and enriched pathways).

### Mitochondria copy number and deletion

To measure mitochondrial copy number, DNA was isolated with the Quick-DNA Miniprep Plus Kit (D4068, Zymo Research) and 8 ng of DNA was used to quantify mitochondrial DNA (mtDNA) encoding the genes NADH dehydrogenase 2 (*ND2*), cytochrome C oxidase 1 (*CO1*) and beta-2 microglobulin (*β2M*) on genomic DNA with the QuantStudio 3 Real-Time PCR System. We calculated mitochondrial DNA copy numbers using the mtDNA/nDNA (nuclear DNA) ratio. To measure mitochondria deletion, the quantity of the mitochondrial genes *ND2* and *CO1* were analysed in relation to NADH dehydrogenase 4 (*ND4*), because *ND4* is known to have a high susceptibility to deletions, while *ND2 and CO1* have not. Primer sequences are available in the supplementary information.

### Statistical Analysis

To analyse qPCR results (*KCNQ1*, *KCNQ1OT1*, NSC transcriptome follow-up), area under curve of neurite outgrowth, western blot results, mitochondria copy number and deletion, as well as apoptosis and proliferation, either KO lines were normalized to their respective isogenic WT control (homKO 1 vs. WT 1, hetKO 1 vs. WT 2) or each cell line was normalized to its respective mock treatment which was set to the value 1 or 100. For analysis, a one-way ANOVA followed by a Bonferroni-adjusted post hoc test was used. To analyse the time response of neurite outgrowth, a two-way ANOVA was performed considering the factors cell line or treatment and time followed by a Bonferroni-adjusted post hoc test. To analyse neuronal activity (voltage responses upon injections of hyperpolarizing current injections, spiking activity upon injections of depolarizing current injections), a two-way ANOVA was used with the factors cell line and injected current followed by a Bonferroni-adjusted post hoc test. To analyse AP properties, rheobase, and EPSC properties, an unpaired, two-tailed Student’s *t*-test was performed. For the analysis of qPCR results to characterize the neurons, a multiple unpaired two-tailed Student’s *t*-test followed by a Holm-Sídák-adjusted post hoc test. For nCounter analysis, multiple unpaired two-tailed Student’s *t*-test were used with two-stage step-up method of Benjamini, Krieger and Yekutieli (FDR = 0.05) to correct for multiple testing. Statistical analysis was performed using GraphPad Prism 10.3.1 (GraphPad Software, San Diego, CA, USA).

For the transcriptomic data analysis, *P*-values were adjusted for multiple testing by DESeq2 using the Benjamini-Hochberg procedure with independent filtering disabled. Genes having an adjusted *P* < 0.05 were considered differentially expressed.

For the proteomic data analysis, differential expression analysis was carried out using linear modelling via the “test_diff()” function in DEP2, which internally applies empirical Bayes statistics based on the limma package^72^. Adjusted *P*-values were calculated using the Benjamini-Hochberg false discovery rate (FDR) correction. Proteins were considered significantly differentially abundant using thresholds of adjusted *P*-value < 0.1 and absolute log2 fold-change > 0.58.

**iPSC culturing, immunocytochemistry, karyotyping, PCR, RT-PCR, qPCR, sequencing, trilineage differentiation, analysis of proliferation and apoptosis, and nCounter gene expression profiling were performed using standard protocols further described in the Supplementary Information as Supplementary materials and methods.**

## Supporting information

Supplementary Table 2 - Differential expressed proteins

Supplementary Table 1 - Differential expressed genes

## Author contribution

DS performed the majority of the experiments, designed the figures, carried out the data analysis, contributed to the data interpretation and writing of the manuscript,

JW, GP, BEG carried out transcriptomic data analysis,

MS, JCG, BEG carried out proteomic data analysis,

CA carried out patch clamp recording analysis,

SA performed the mitochondria DNA copy number and deletion analysis and contributed to neurite outgrowth analysis

MH contributed to RNA sequencing data analysis and performed the western blot experiments

RR performed the nCounter analysis,

CL contributed to western blot experiments and reviewed the manuscript

LB performed the TMRM staining and data analysis

KV contributed to neuronal induction and neurite outgrowth experiments,

KB performed the karyotyping of all iPSC lines

DM contributed to the data interpretation of the Incucyte experiments

ML carried out the mass spectrometry analysis

AA, JMR contributed to conceptualization

DS, BEG, JW, CA, GP, MS, RR, KV, KB, DM, GAR, ML, AA, JMR, JCG and BF reviewed and edited the paper

SB designed the study, contributed to the data analysis, interpreted the data and wrote the manuscript

All co-authors commented on the final manuscript

## Acknowledgements

DS, JW, GP, KV, JR, BF, SB were supported by funding from the European Union’s Horizon 2020 research and innovation program under Grant agreement no. 847879 (PRIME, Prevention and Remediation of Insulin Multimorbidity in Europe). DS and SB were also supported by the Medical Faculty of the Ruprecht-Karls-University Heidelberg. SA was supported by the ERASMUS+ program, DM was supported by the Chica and Heinz Schaller Foundation. BF received relevant funding from the Dutch Ministry of Education, Culture and Science of the government of The Netherlands for the NWO Gravitation programme GUTS (grant 024.005.011). BF has received educational speaking fees and travel support from Medice. JCG was supported by the Ad Astra Programme of University College Dublin and has received funding from the Irish Research Council (grant no. GOIPG/2024/5062).

We acknowledge the technical support of the Core Facility for Mass Spectrometry and Proteomics of the Center for Molecular Biology (ZMBH) of Heidelberg University. We thank Ute Bach and Marcin Luzarowski for their support with mass spectrometry analysis. The Core Facility for Mass Spectrometry and Proteomics is funded by the ZMBH and partially funded by the CellNetworks Core Technology Platform (CCTP) of Heidelberg University. The CCTP is funded in part by the Federal Ministry of Education and Research (BMBF) and the Ministry of Science Baden Württemberg within the framework of the Excellence Strategy of the Federal and State Governments of Germany. We acknowledge Alexandra Köppel for her technical assistance in performing the karyotyping.

## Conflicts of interest

All authors declare no conflicts of interest.

## Data availability

The processed and raw (FASTQ) gene expression data for the NSCs and neurons is available through the gene expression omnibus (GSE number to be allocated). The proteomic data for the NSCs and neurons will be available on the PRoteomics IDEntifications Database (PRIDE). Proteomic data are available via ProteomeXchange with identifier PXD068563.

## Notes

### Competing Interest Statement

The authors have declared no competing interest.

