## Supplementary information - Supplementary figures & Supplementary materials and methods for "KCNQ1 regulates human neuronal development through mitochondrial and insulin signalling pathways"

### Supplementary materials and methods

**Supplementary figures**

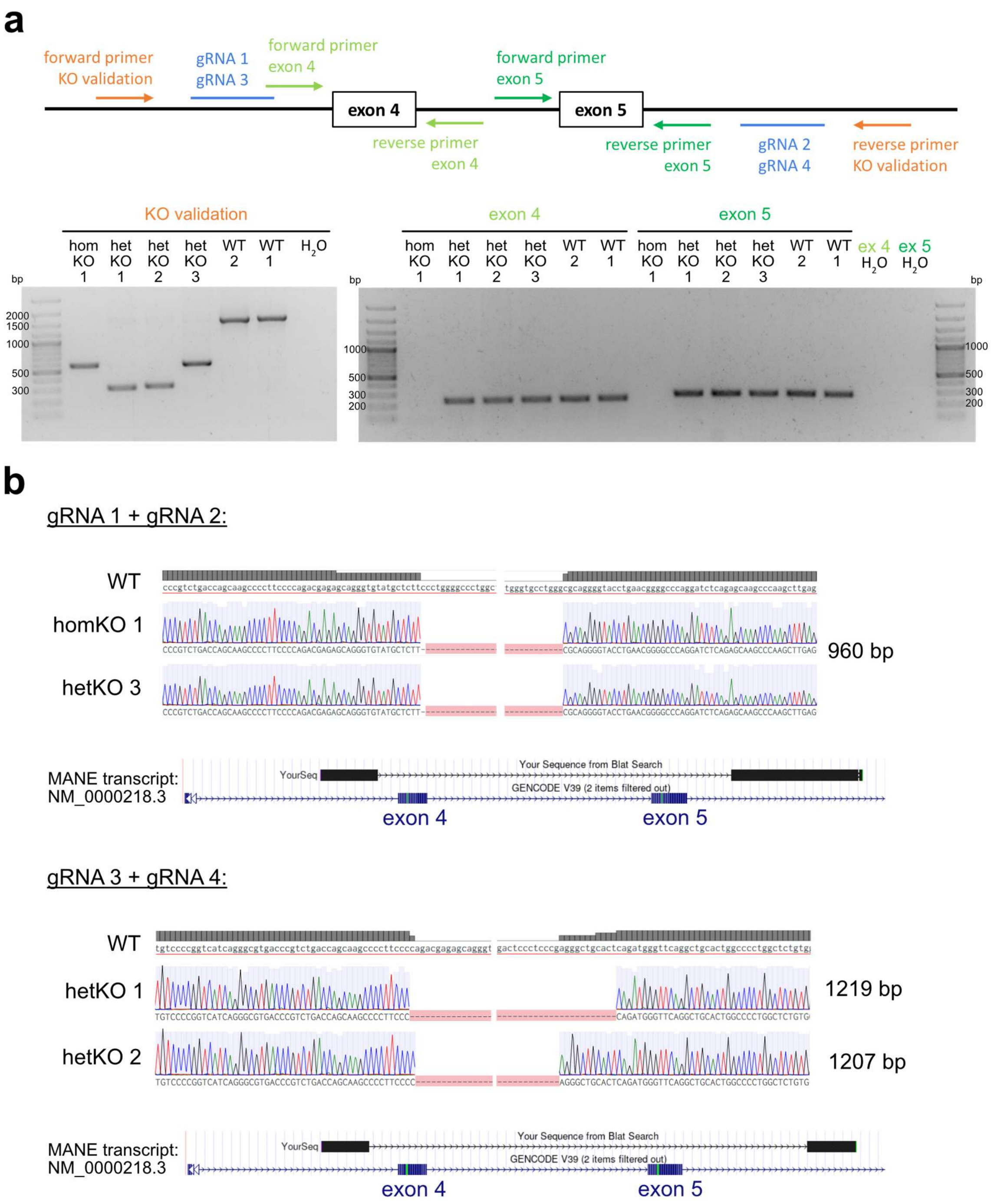


**Supplementary Figure 1. Validation of *KCNQ1*-KOs in iPSCs. a** PCR was used to screen and validate the *KCNQ1*-KO in the iPSC lines. The upper scheme shows where primers bind in relation to exon 4 and 5 and the gRNAs. For a first KO validation, a primer pair flanking the gRNAs upstream and downstream were used. Without a deletion the PCR product results in 1520 bp (WT cell lines). With a deletion, the PCR product for gRNA 1 + gRNA 2 (homKO 1, hetKO 3) results in a product size of 558 bp; and gRNA 3 + gRNA4 (hetKO 1, hetKO 2) results in a product size of 312 bp. To determine whether a KO is homozygous or heterozygous, we additionally performed a PCR with specific primers for either exon 4 or 5. The homozygous KO does not show any band, whereas all heterozygous KOs and WT lines show a band for both exons, as expected. Band sizes: exon 4: 235 bp, exon 5: 268 bp. **b** Sanger sequencing results of KO cell lines. For each gRNA combination the upper picture shows sequencing alignment of KO compared to WT sequence to show the exact cutting side of each gRNA and the deletion size. The lower picture visualizes the deletion position on the *KCNQ1* gene by comparing the sequence to the MANE *KCNQ1* transcript (UCSC browser, hg38). Chromosome positions of deletions: homKO 1/hetKO 3: 2.571.269-2.572.229, hetKO 1: 2.571.424-2.572.461, hetKO 2: 2.571.243-2.572.450.

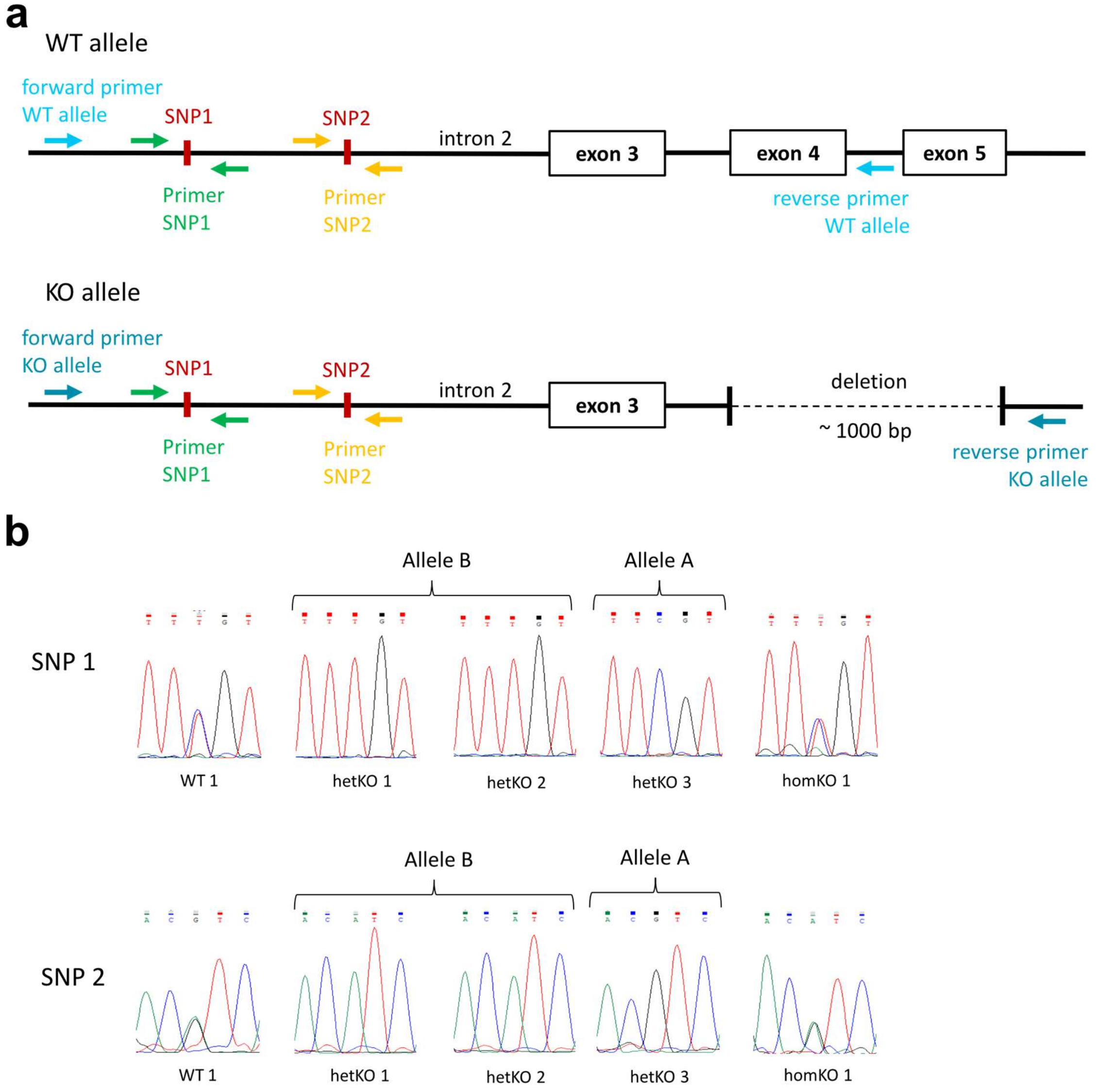


**Supplementary Figure 2. Assignment of deletion to allele of the heterozygous KO cell lines.** To distinguish which allele was edited in the heterozygous *KCNQ1*-KO iPSC lines, two heterozygous SNPs in intron 2 (rs179484, rs179486) of *KCNQ1* were genotyped by Sanger sequencing. **a** Primer design for PCR and sequencing. For the specific amplification of the WT allele, the reverse primer was set within the region of the deletion. To amplify the KO allele, the reverse primer was set downstream of the deletion. Two specific primer pairs were designed to amplify and sequence the region of the two SNPs. **b** Representative sequencing results of the PCR to detect the WT allele (for WT and heterozygous KO cell lines) or KO allele (for homozygous KO cell line). In both WT lines and in the homozygous KO, the SNPs are heterozygous, proving that both alleles are amplified (in WT lines only WT alleles, in homKO 1 only KO alleles). In all heterozygous cell lines only one SNP variant per allele is visible, proving that with each PCR only one allele is amplified, either WT or KO allele. SNP 1 (rs179484) displays a shift from C to T with C as the major and T as the minor allele. SNP 2 (rs179486) shifts from G to A with G as the major and A as the minor allele. The major allele was defined as allele A, the minor allele was defined as allele B. HetKO 1 and 2 carry the deletion on allele A, hetKO 3 carries the deletion on allele B.

**a**

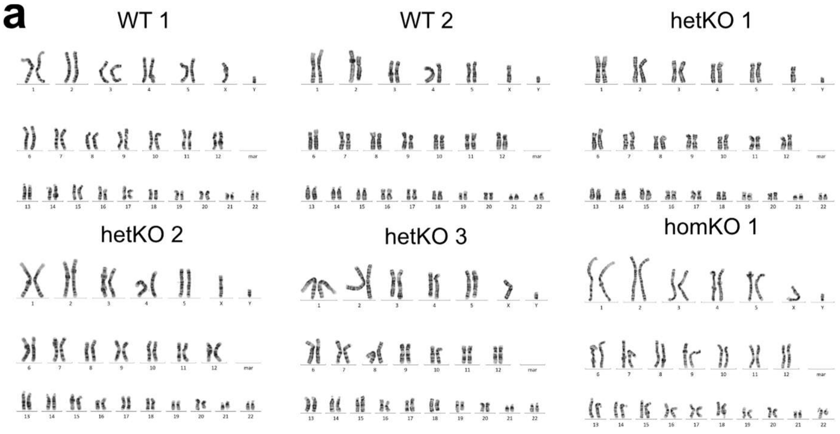


**b**

| gRNA | Predicted Gene | Predicted Position | Off-target Editing |
|---|---|---|---|
| **WT 1, homKO 1, and hetKO 3** | | | |
| gRNA 1 | *RPS19* | Exonic | no |
| | *ELAV4* | Exonic | no |
| | *NES* | Exonic | no |
| | *CLDN9* | Intronic | no |
| gRNA 2 | *PCBD1* | Exonic | no |
| **WT 2, hetKO 1, and hetKO 2** | | | |
| gRNA 3 | *FAM20C* | Exonic | no |
| | *ADAM15* | Intergenic | no |
| | *ELOCP20* | Intergenic | no |
| | *MCF2L2* | Intronic | no |
| | *LARGE* | Intronic | no |
| | *PKHD1* | Intronic | no |
| gRNA 4 | *RAD51B* | Exonic | no |
| | *PRR5L* | Exonic | no |
| | *CCDC57* | Exonic | no |
| | *THEM6* | Intronic | no |

**c**

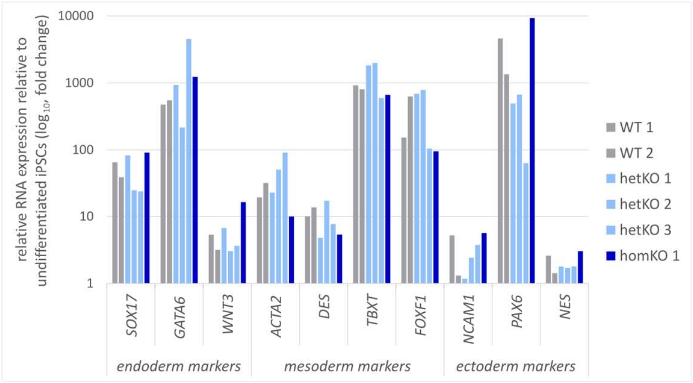


**d**

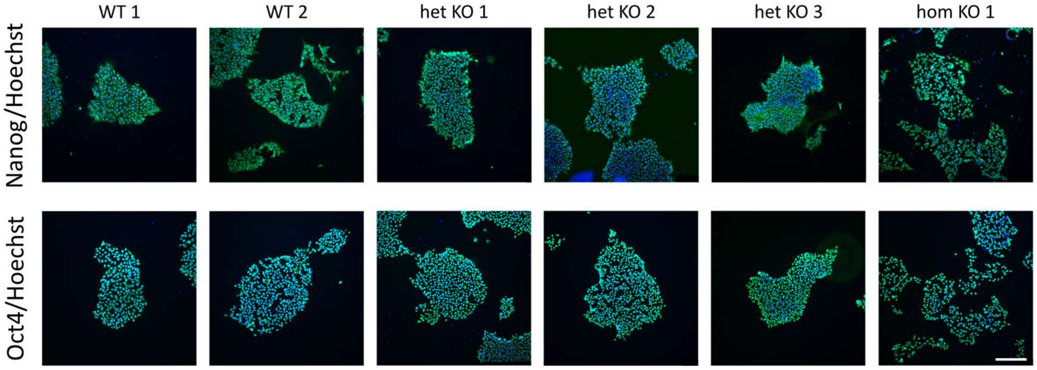


**Supplementary Figure 3. Quality assessment of generated iPSC lines. a** Karyotype images of all iPSC lines. **b** Sequencing of predicted putative off-target sites revealed no evidence for unspecific editing. **c** Relative RNA expression of endoderm, mesoderm, and ectoderm markers after trilineage differentiation of iPSCs. All generated cell lines show a higher expression of the germ layer specific markers relative to their respective undifferentiated iPSCs. **d** Immunofluorescence images of pluripotency markers. All iPSC lines express both pluripotency markers, Nanog or OCT4 (green); nuclei staining: Hoechst (blue). Scalebar = 100 µm. *RPS19*: ribosomal protein S19, *ELAV4*: ELAV like RNA binding protein 4, *NES*: nestin, CLDN9: claudin9, *PCBD1*: pterin-4 alpha-carbinolamine dehydratase 1, *FAM20C*: FAM20C Golgi Associated Secretory Pathway Kinase, *ADAM15*: ADAM metallopeptidase domain 15, *ELOCP20*: elongin C pseudogene 20, *MCF2L2*: MCF.2 cell line derived transforming sequence-like 2, *PKHD1*: PKHD1 ciliary IPT domain containing fibrocystin/polyductin, *RAD51B*: RAD51 paralog B, *PRR5L*: proline rich 5 like, *CCDC57*: coiled-coil domain containing 57, *THEM6*: thioesterase superfamily member 6, *SOX17*: SRY-box transcription factor 17, *GATA6*: GATA binding protein 6, *WNT3*: Wnt family member 3, *ACTA2*: actin alpha 2, smooth muscle, *DES*: desmin, *TBXT*: T-box transcription factor T, *FOXF1*: forkhead box F1, *NCAM1*: neural cell adhesion molecule 1, *PAX6*: paired box 6.

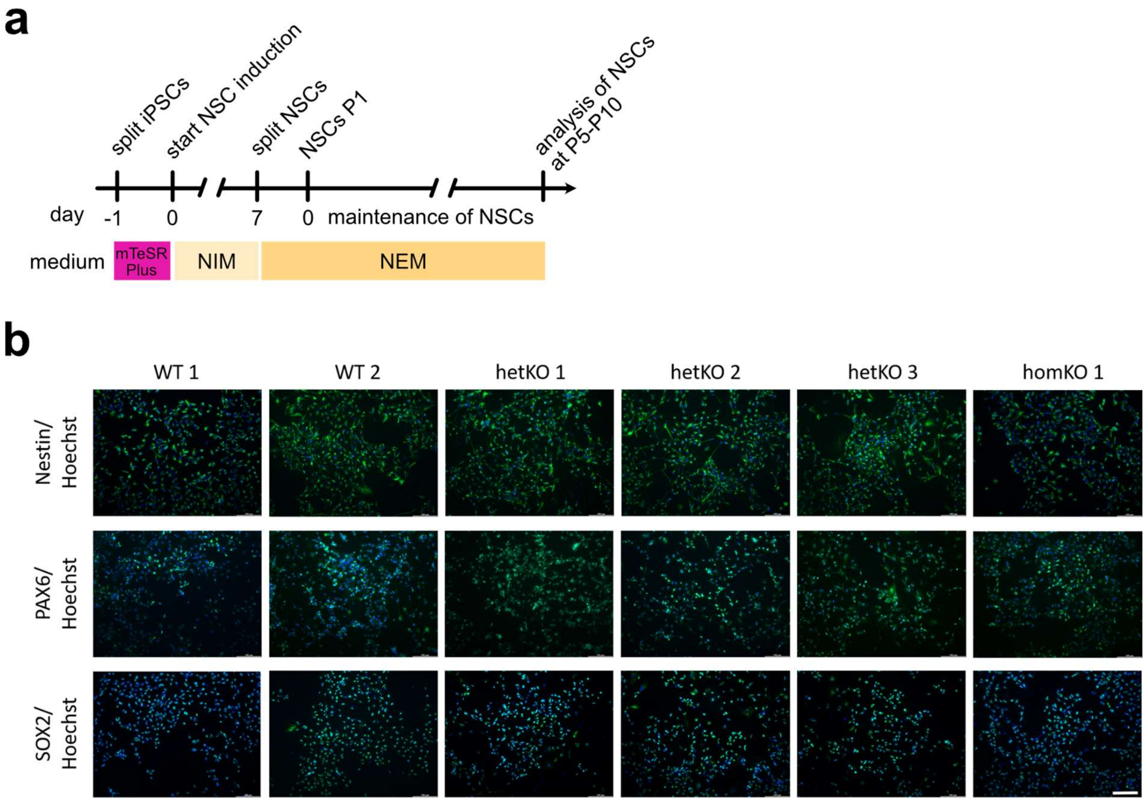


**Supplementary Figure 4. Characterization of induced NSCs. a** Scheme of NSC induction from iPSCs according to Yan et al.[1]. NIM: neural induction medium, NEM: neural expansion medium. **b** Immunofluorescence images of NSC markers in green (Nestin, PAX6, SOX2) and nucleus staining in blue (Hoechst). All cell lines express all three NSC marker. Scalebar = 100 µm.

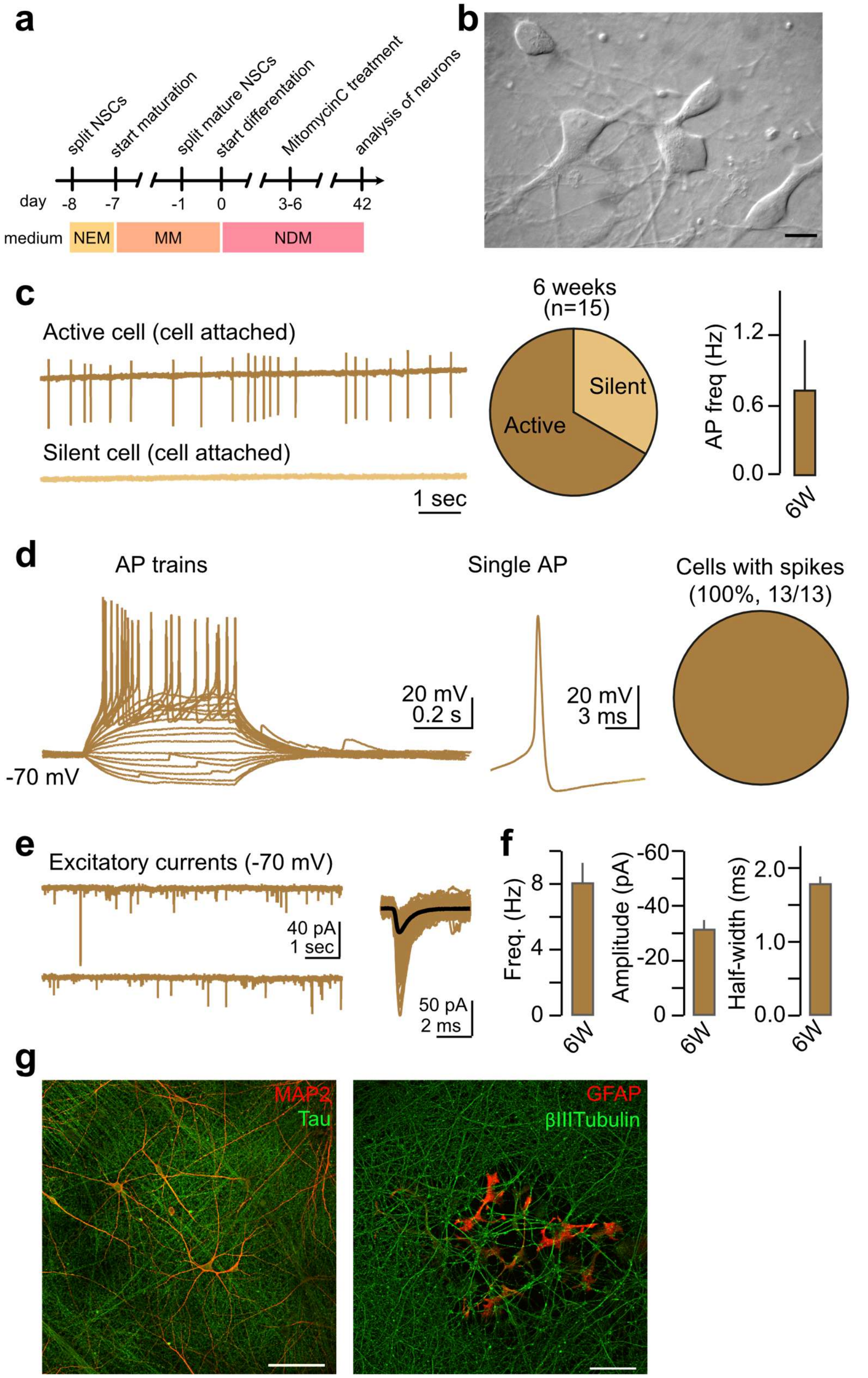


**Supplementary Figure 5. Generation and basic characterization of differentiated forebrain neurons.** **a** Scheme of differentiation protocol according to Yan et al.[1]. NEM: neural expansion medium, MM: maturation medium, NDM: neuronal differentiation medium. **b** Differential-infrared contrast image of 6-week-old neurons generated as indicated in (a). Scale bar: 20 µm. **c** Cell-attached recordings of 6-week-old neurons. Under physiological recording conditions, most neurons fired spontaneous spikes at ~0.7 Hz frequencies. **d** Whole-cell current clamp recordings of 6-week-old neurons. All recorded cells fired action potentials upon positive current injections. **e** Whole-cell voltage clamp recordings of 6-

week-old neurons, which displayed prominent EPSCs under physiological conditions. **f** Summary graphs for EPSC frequency (left), amplitude (middle), and half-width (right). Summary graphs in c (right) and f represent mean ± SEM. Number of experiments (c:15 cells; f: 16 cells). AP: action potential. **g** Representative immunofluorescence microscopy images with neuronal (MAP2, Tau, and β3-Tubulin) and glial (GFAP) markers after 6 weeks of differentiation. MAP2: microtuble-associated protein 2; GFAP: glial fibrillary acid protein. Scale bar: 100 µm.

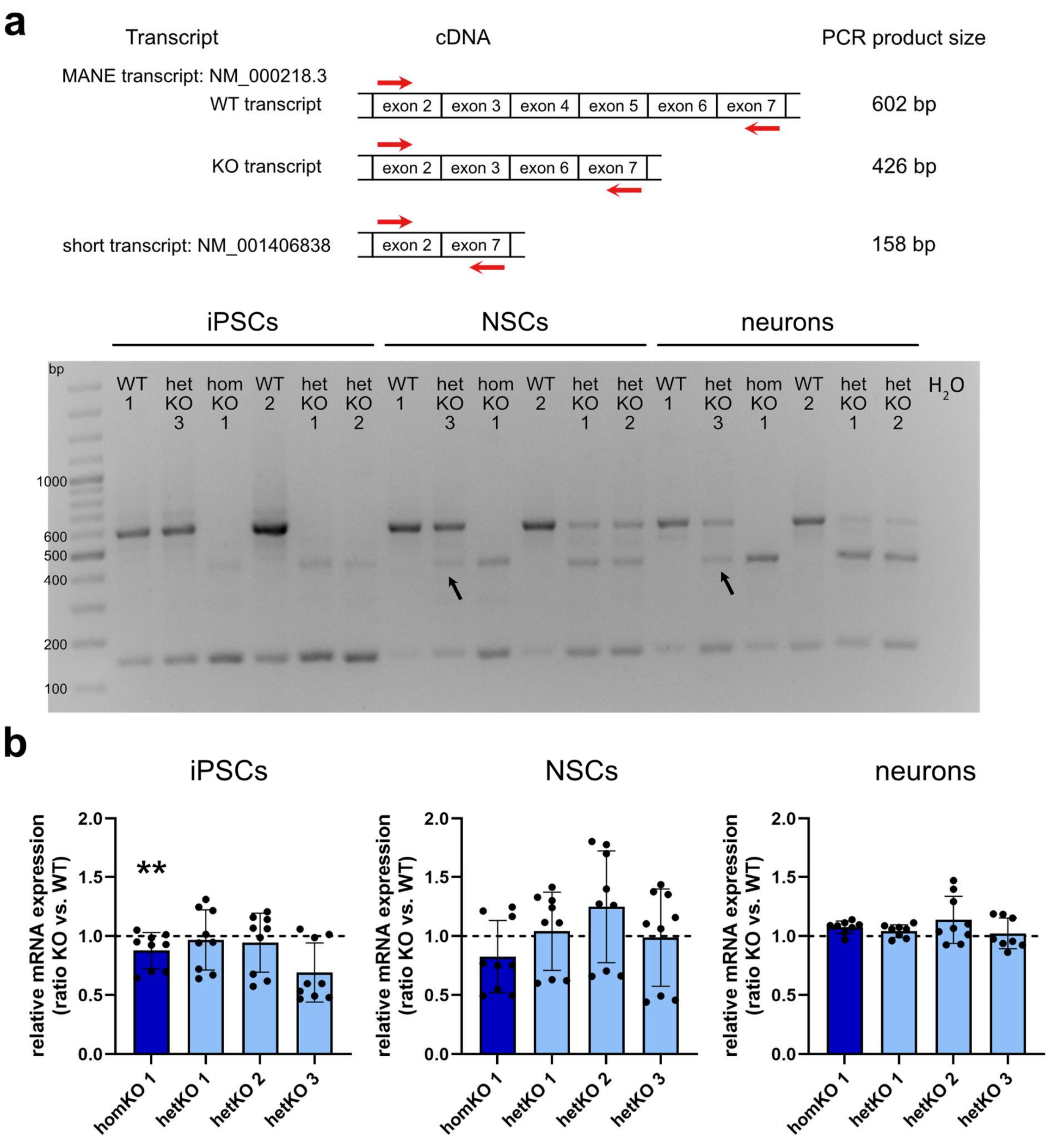


**Supplementary Figure 6. Expression analysis of *KCNQ1* isoforms and long non-coding RNA *KCNQ1OT1*.** Different *KCNQ1* isoforms have been annotated in the UCSC Human Genome Browser (hg 38) with the MANE (Matched Annotation from the NCBI and EMBL-EBI) transcript, containing exons 2 to 7, and a more recently published short transcript lacking exons 3 to 6 and 8. To date, this isoform has only been reported in RNA sequencing data, and its translation into a functional protein remains unconfirmed. If translated, the resulting protein would lack transmembrane segments 2 to 4 (part of the voltage-sensing domain) and segment 5 (part of the pore domain), making it unlikely to form a functional potassium channel. To elucidate imprinting differences between cell types, we performed PCR on cDNA (RT-PCR) to identify transcription of WT and KO alleles in iPSCs, NSCs, and neurons for the different genotypes. **a** Top: Illustration of the different *KCNQ1* transcripts and the selected primers targeting exon 2 (forward) and exon 7 (reverse) which result in different product sizes based on the number of exons included in the transcript. Bottom: PCR results showing expression of the MANE-WT (band size 602 bp) and -KO transcript (band size 426 bp), and of the short transcript (band size of 158 bp). In iPSCs, WT 1, WT 2, and hetKO 3 (carrying the deletion on the inactive allele) show the WT band.

HomKO 1, hetKO 1, and hetKO 2 (both hetKO carrying the deletion on the active allele) show the KO band. In NSCs, both WT lines show the WT band whereas all three heterozygous KO lines show both bands, WT and KO. The homKO 1 NSCs show expression of only the MANE-KO transcript. In neurons, the WT cell lines show the WT band and all heterozygous KO lines show WT and KO expression. The homKO 1 neurons show the KO band of the MANE transcript. All cell lines and cell types express the short *KCNQ1* transcript NM_001406838.1 lacking exon 3-6 (band size: 158 bp). Even though hetKO 3 carries the KO on the inactive allele, imprinting patterns change and both alleles become transcribed in NSCs and neurons (indicted by arrows). **b** Relative *KCNQ1OT1* expression in iPSCs, NSCs, and neurons. In iPSCs only a difference between WT 1 and hetKO 3 was identified ($P$=0.003, rest $P$>0.05) with almost no effect size. In NSCs and neurons no difference was found (all $P$>0.05). One-Way ANOVA with Bonferroni-adjusted post hoc test, mean ± SD, testing WT 1 vs. hetKO 3 and homKO 1, and WT 2 vs. hetKO 1 and hetKO 2, n=3. KO values are normalized to their respective WT control, which are set to the value 1 and indicated by the dotted line.

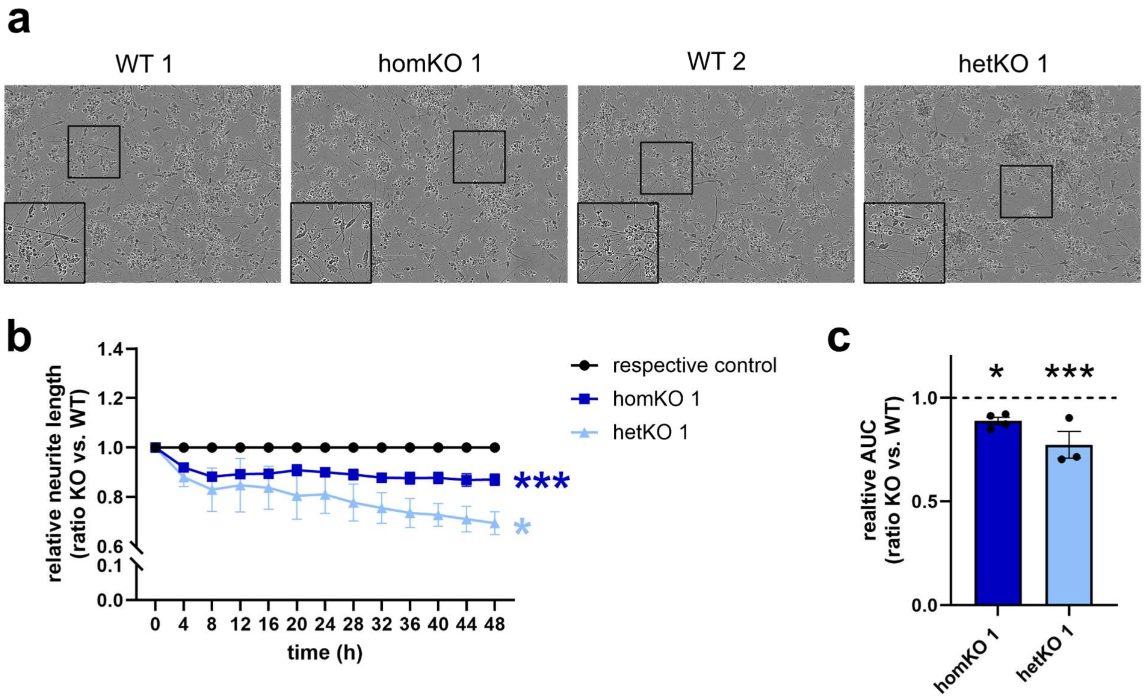


**Supplementary Figure 7. Neurite outgrowth in *KCNQ1*-KO and WT cell lines.** Cells were differentiated with an alternative protocol modified according to Qi et. al.[2] which is described in detail in the Supplementary materials and methods. **a** Representative pictures of all cell lines at time point 48h. **b, c** Both KO lines have a reduced neurite length compared to their isogenic control. homKO 1 vs. WT 1: n=4, hetKO 1 vs. WT 2: n=3. **b** Two-way ANOVA (time x cell line) with Bonferroni-adjusted post hoc test, mean ± SEM. Asterisks indicate significances of ANOVA for factor cell line: homKO 1: *P*=0.0004, F= 49.45; hetKO 1: *P*=0.013, F=14.2. Post hoc test: homKO 1: 8h: *P*=0.0250, 12h: *P*=0.0058, 32h: *P*=0.0258, rest all *P*>0.05; hetKO 1: all *P*>0.05. **c** Calculation of the area under curve (AUC), followed by a one-way ANOVA with Bonferroni-adjusted post hoc test, mean ± SEM, *P*=0.0008, F=13.11, post hoc test: homKO 1: *P*=0.0301, hetKO 1: *P*=0.0008.

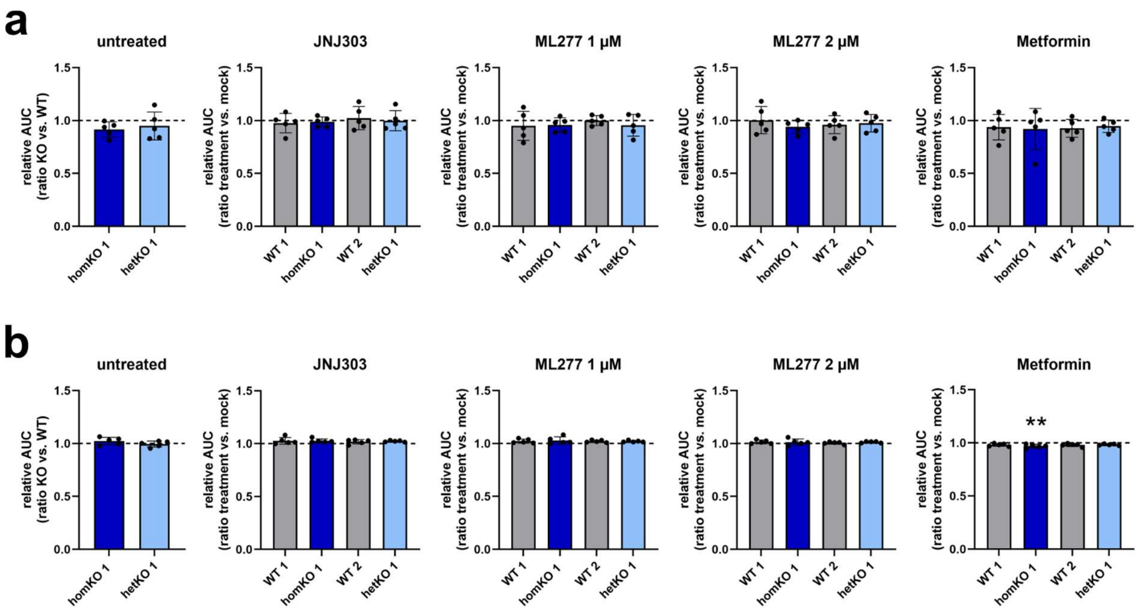


**Supplementary Figure 8. Analysis of cell proliferation and apoptosis.** Proliferation and apoptosis were either measured untreated or treated with KCNQ1 antagonist JNJ303 (1 µM), KCNQ1 agonist ML277 (1 µM, 2 µM) or Metformin (1 mM) with the IncuCyte system. **a** Apoptosis analysis revealed no differences. **b** Proliferation analysis showed only for treatment with Metformin a significantly reduced proliferation in homKO 1 (*P*=0.004, rest all *P*>0.05) with almost no effect size. One-way ANOVA with Bonferroni-adjusted post hoc test, mean ± SD n=5. Untreated: KO values are normalized to their respective WT control; treated: values are normalized to mock treatment of the same cell line. Either WT (untreated) or mock treatment (treated) are set to the value 1 and indicated by the dotted line.

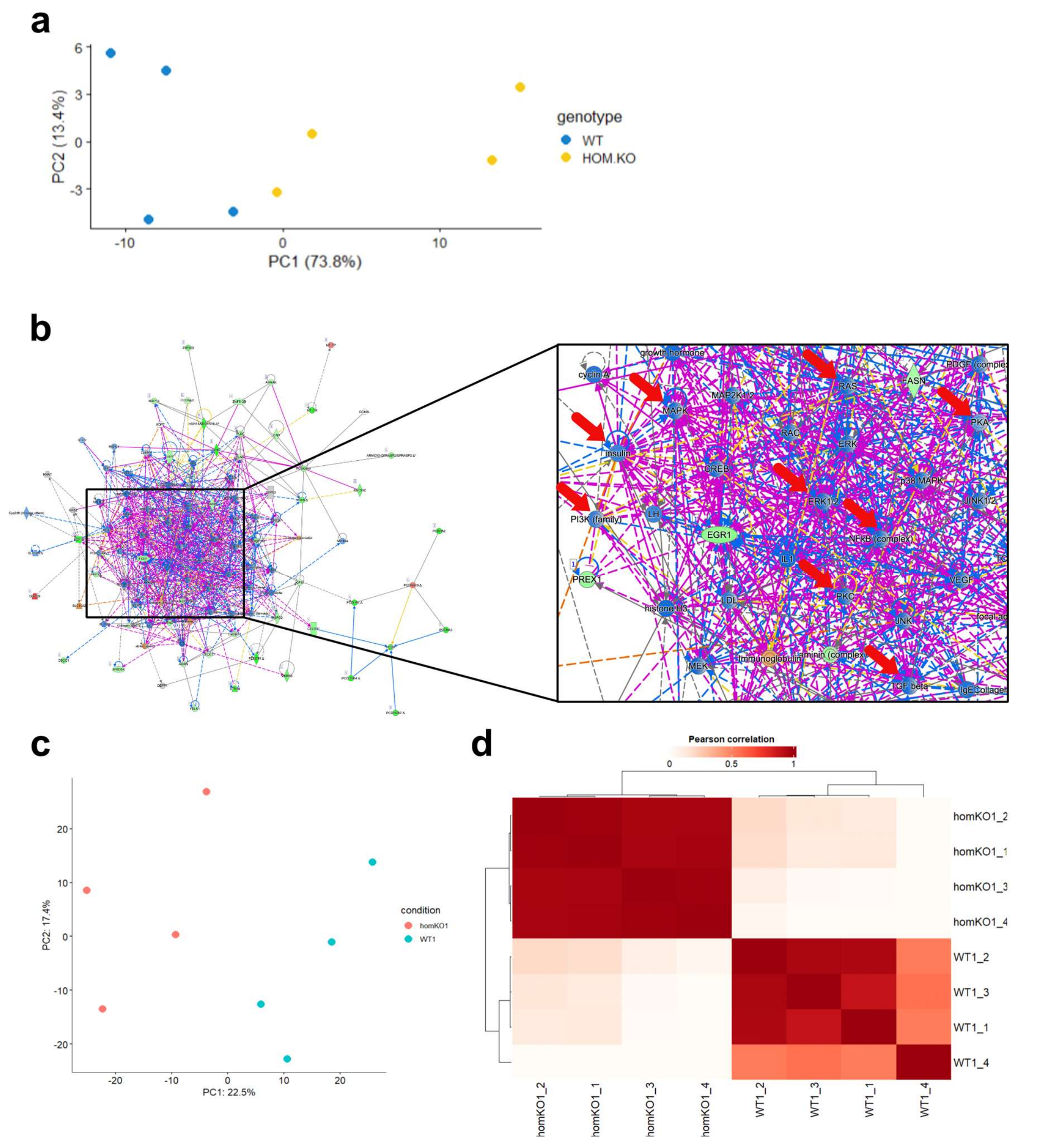


**Supplementary Figure 9. Analysis of RNA sequencing data and mass spectrometry data from WT 1 and homKO 1 NSCs. a** PCA of RNA sequencing data. PC1 and PC2 account for 73% and 13.4% of the variance in the data, respectively. Samples cluster according to the genotype conditions separating along PC1. Color-coding by condition reveals distinct clustering, suggesting that the genotype is a major driver of variance. **b** Network enrichment analysis of the transcriptomic data in NSCs (IPA). The three most significantly enriched networks for all DEGs were merged. The network enrichment analysis points towards an involvement of insulin signalling (including downstream molecules RAS, MAPK, ERK, PI3K), TGFβ, NFκB, PKC and PKA signalling (indicated by red arrows). Genes: green: downregulated in expression, red: upregulated in expression, blue: predicted to be inhibited, orange: predicted to be activated, grey: differentially expressed but not significantly, white: not present; arrows: purple: overlap between networks, orange: predicted positive effect, blue: predicted negative effect, yellow: inconsistent/unpredicted effect, grey: no predicted effect. **c** PCA of proteomic data from WT 1 and homKO 1 NSCs. PC1 and PC2 explain 22.5% and 17.4% of the data variance. According to the separation of genotypes in PC1, the genotype seems to be the main cause of variance between groups. **d** Pearson correlation analysis revealed clustering of the NSCs according to the different genotypes.

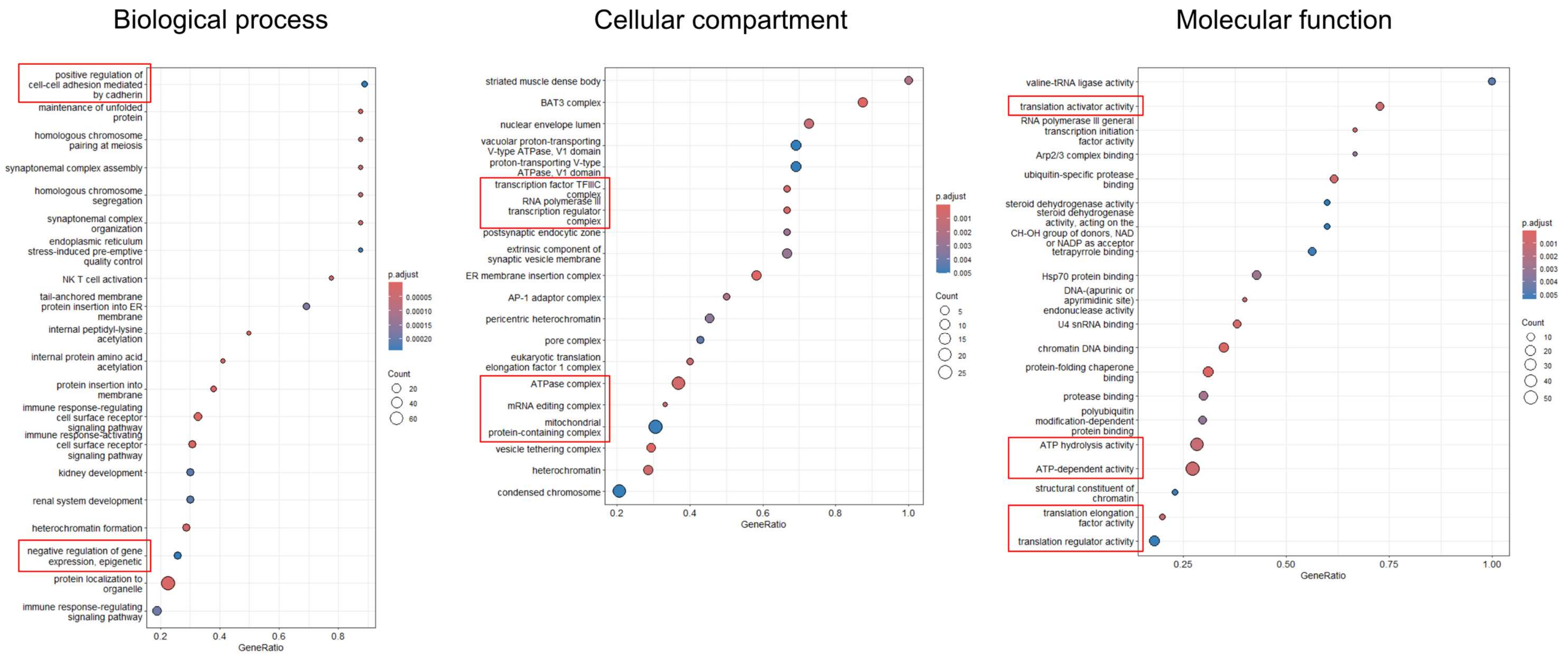


**Supplementary Figure 10. GSEA comparing homKO 1 to WT 1 in NSCs.** All nominal significant proteins with their log2fold changes were considered for the analysis. The top 20 affected biological processes, cellular compartments, and molecular functions are shown. Most interesting findings were labelled in red boxes and are summarized in the main Table 1.

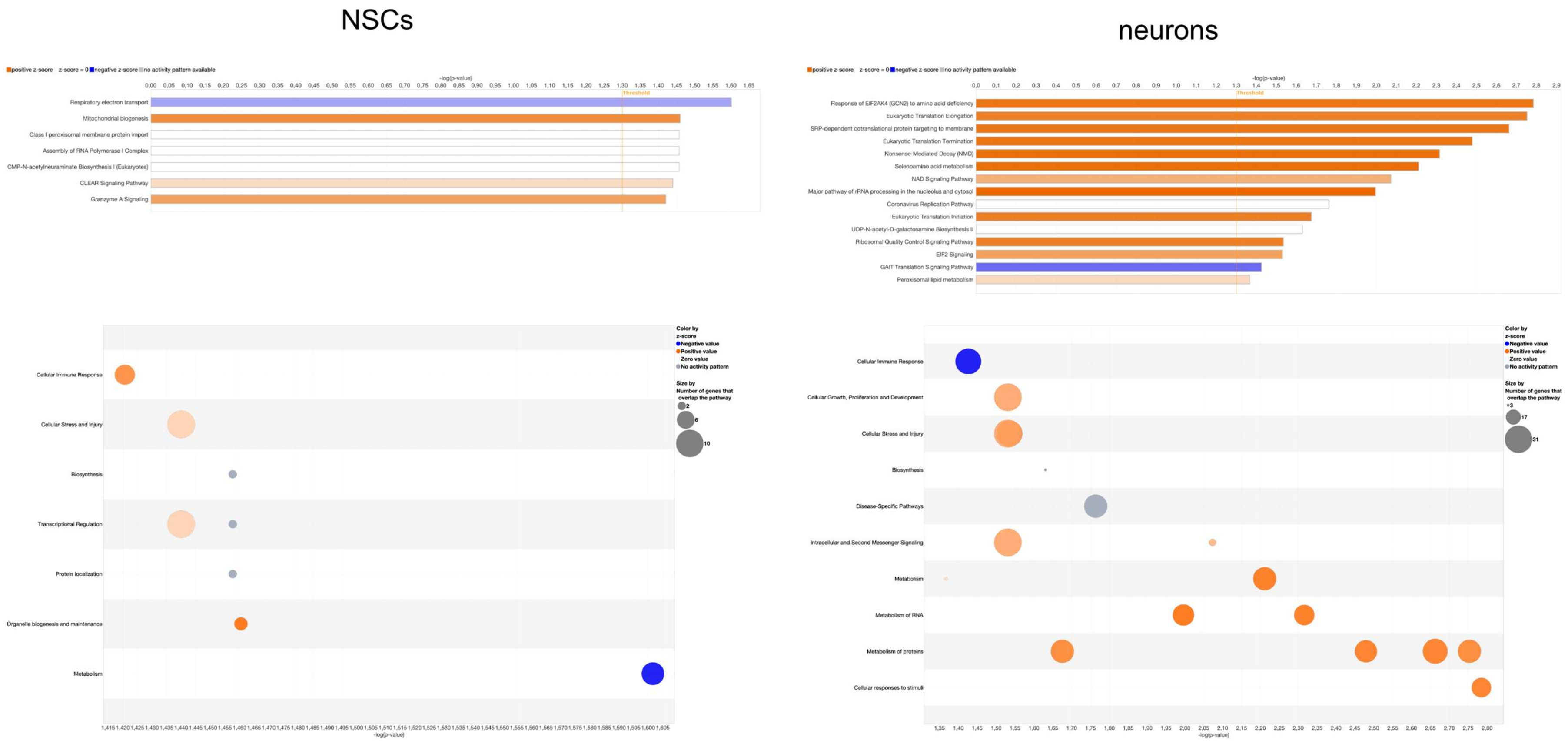


**Supplementary Figure 11. IPA canonical pathway enrichment analyses of proteomic data in NSCs and neurons comparing homKO 1 to WT 1.** All nominally significant proteins were considered for the analysis together with their log2fold changes. The bar graph at the top shows the significantly enriched canonical pathways (i.e., with nominally significant *P*-value), with orange (positive z-scores) and blue (negative z-cores) indicating predicted activation and inhibition of the pathway, respectively. The bubble blot at the bottom provides a holistic view of all identified cellular mechanisms.

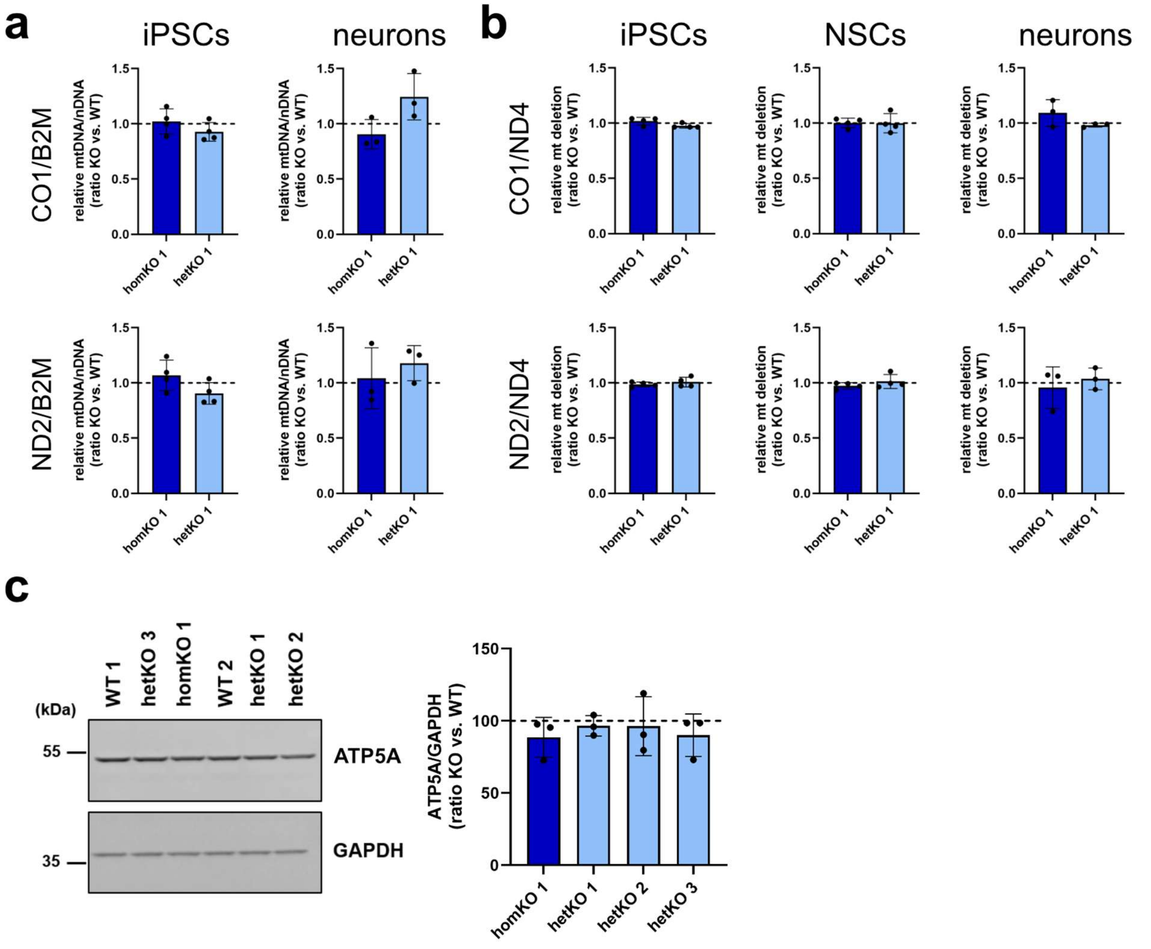


**Supplementary Figure 12: Analysis of mitochondria in iPSCs, NSCs, and neurons. a** Quantification of mitochondrial DNA in iPSCs and neurons. No difference was observed in any cell line or cell type. iPSCs: n=4, neurons: n=3. **b** Quantification of mitochondrial deletion in iPSCs, NSCs, and neurons showed no difference for both KO lines in all three cell types. iPSCs: n=4, neurons: n=3. **c** Western blot analysis of ATP5A expression in neurons (left: representative membrane image, right: quantification). ATP5A shows similar expression levels for all cell lines. n=3. **a-c** One-way ANOVA with Bonferroni-adjusted post hoc test, mean ± SD, all $P$>0.05.

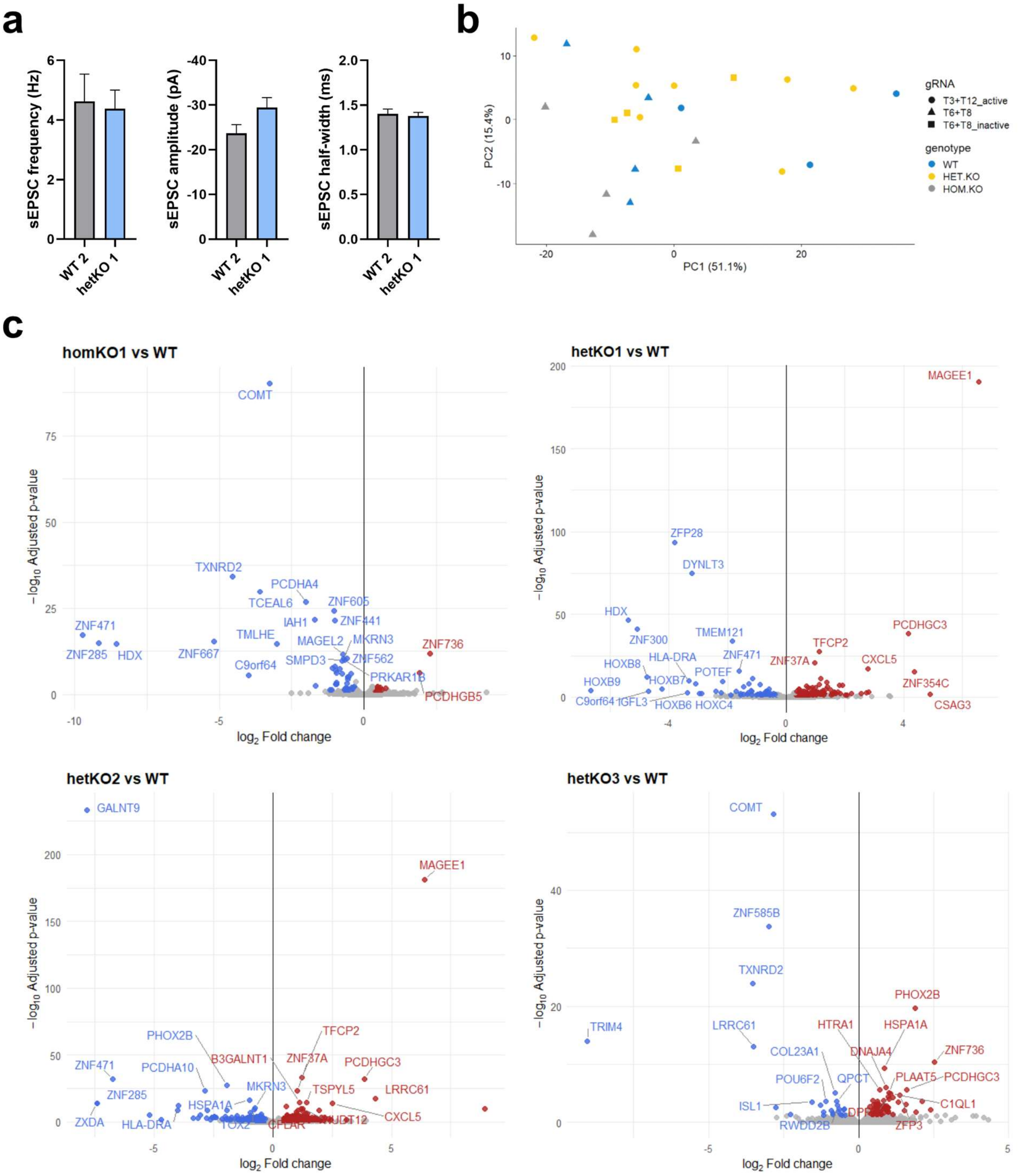


**Supplementary Figure 13. Electrophysiology measurements of hetKO 1 and WT 2, and transcriptomic analysis in *KCNQ1*-KO neurons. a** Measurements of EPSC frequency, amplitude, and half-width of hetKO 1 and WT 2 does not show any differences between WT and KO (all $P>0.05$). Unpaired two-tailed Student's *t*-test, mean ± SEM. **b** PCA of RNA sequencing data from WT 1, WT 2, hetKO 1, hetKO 2, hetKO 3, and homKO 1 neurons did not reveal distinct clustering among the different cell lines. PC1 and PC2 explain 51.1 % and 15.4% of the variance, respectively, but this variation does not align with the experimental groups. Samples from different genotypes are interspersed across the PCA plot, with only slight separation for homKO 1. The overlapping distribution suggests either minimal transcriptional differences across conditions or that relevant variation lies outside the first two components. **c** Volcano plots showing differentially expressed protein coding genes in neurons. Genes with FDR < 0.05 and log2fold change >1.3 are shown (red: significantly upregulated genes, blue: significantly downregulated genes).

**a**

| Comparison | Total | Up | Down |
|---|---|---|---|
| homKO 1 vs WT | 64 | 14 | 50 |
| hetKO 1 vs WT | 189 | 125 | 64 |
| hetKO 2 vs WT | 377 | 226 | 151 |
| hetKO 3 vs WT | 100 | 74 | 26 |

**b**

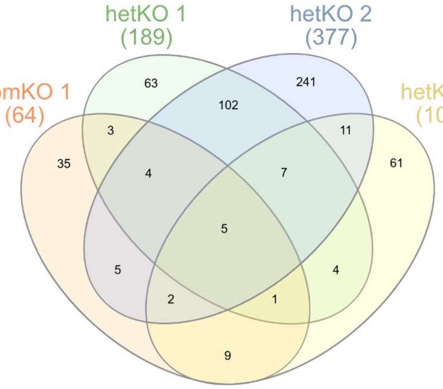


**c**

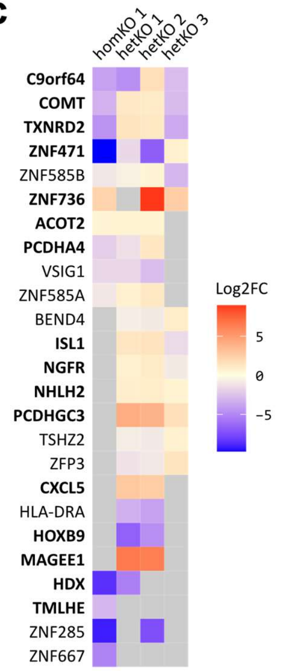


**d**

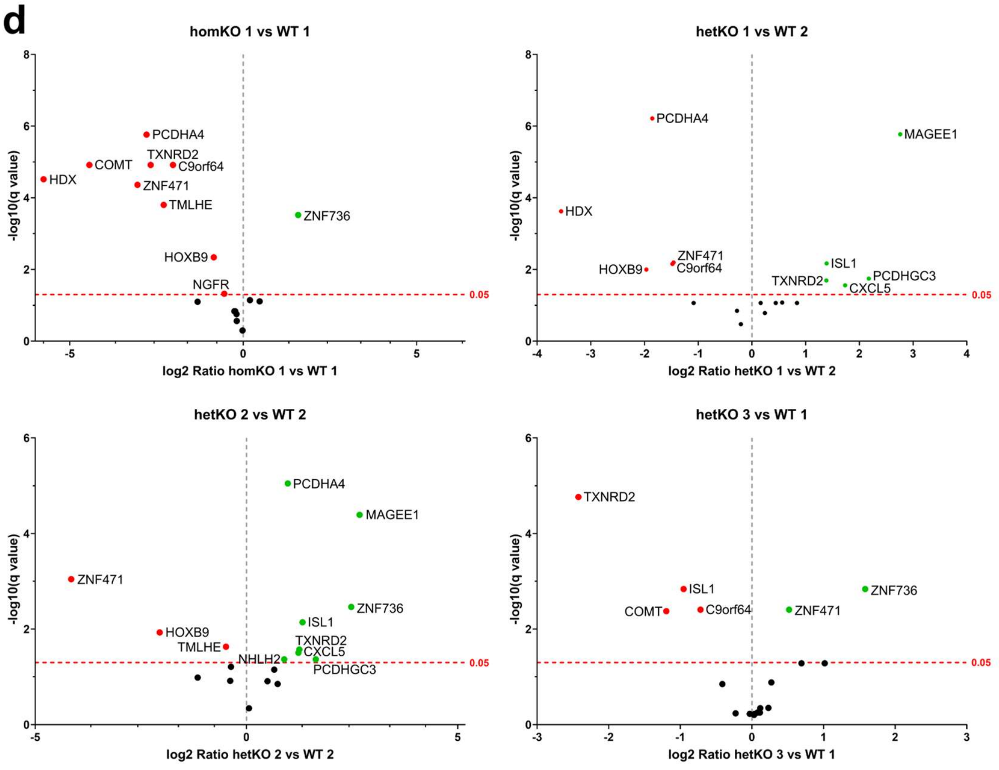


**Supplementary Figure 14. Differential expressed protein coding genes in *KCNQ1*-KO neurons. a** Table with the number of significantly protein-coding DEGs (FDR < 0.05) identified by whole transcriptome analysis. **b** Venn diagram of significantly differentially expressed protein coding genes in neurons. **c** Heatmap of selected protein coding genes. Five protein-coding genes (*C9orf64*, *COMT*, *TXNRD2*, *ZNF471*, and *ZNF585B*) were consistently differentially expressed across all four *KCNQ1*-KO neuronal lines compared to WT controls. *C9orf64* encodes a protein involved in tRNA-guanine transglycosylation, while *COMT* encodes catechol-O-methyltransferase. *TXNRD2* regulates mitochondrial redox homeostasis and reactive oxygen species levels. The zinc finger genes (*ZNF471, ZNF585B*) encode transcriptional regulators. Moreover, in three KO cell lines overlapping differential expression was seen for genes with a role in cell adhesion (*PCDHA4*, *PCDHGC3*), neurodevelopment, differentiation, neuronal survival (*NGFR*, *NHLH2*), insulin regulation (*ISL1*), and transcriptional regulation (*ZFP3*, *ZNF585A*, six different *HOX* (e.g. *HOXB9*) genes, and *MAGEE1* which enhances ubiquitine-ligase function. Genes labelled in bold were further followed-up by nCounter analysis. Like in the NSCs, *KCNQ1*, other KCNQ family members (*KCNQ2–5*), and key KCNQ1 interaction partners were not found differentially expressed in any of the neuronal samples analysed. **d** Gene expression analysis with nCounter. Volcano plots showing differential expression of the followed-up genes.

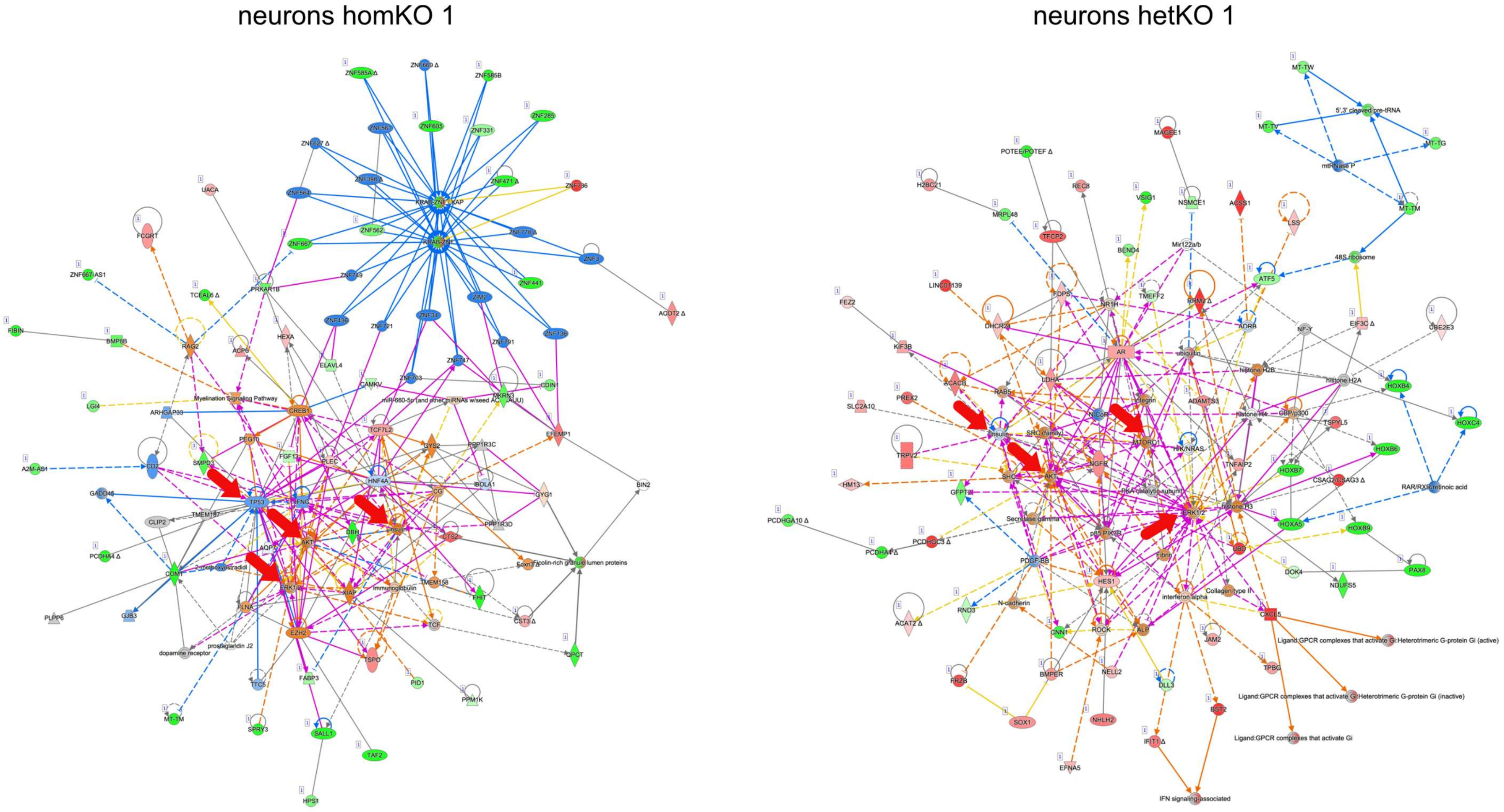


**Supplementary Figure 15. Network enrichment analyses of transcriptomic data in homKO 1 and hetKO 1 neurons.** The three most significantly enriched networks for all DEGs were merged (IPA). The network enrichment analyses point towards an involvement of insulin signalling (insulin, AKT, ERK) in homKO 1 and hetKO 1 neurons and TP53 in homKO 1 neurons (indicated by red arrows). Genes: green: downregulated in expression, red: upregulated in expression, blue: predicted to be inhibited, orange: predicted to be activated, grey: differentially expressed but not significantly, white: not present; arrows: purple: overlap between networks, orange: predicted positive effect, blue: predicted negative effect, yellow: inconsistent/unpredicted effect, grey: no predicted effect.

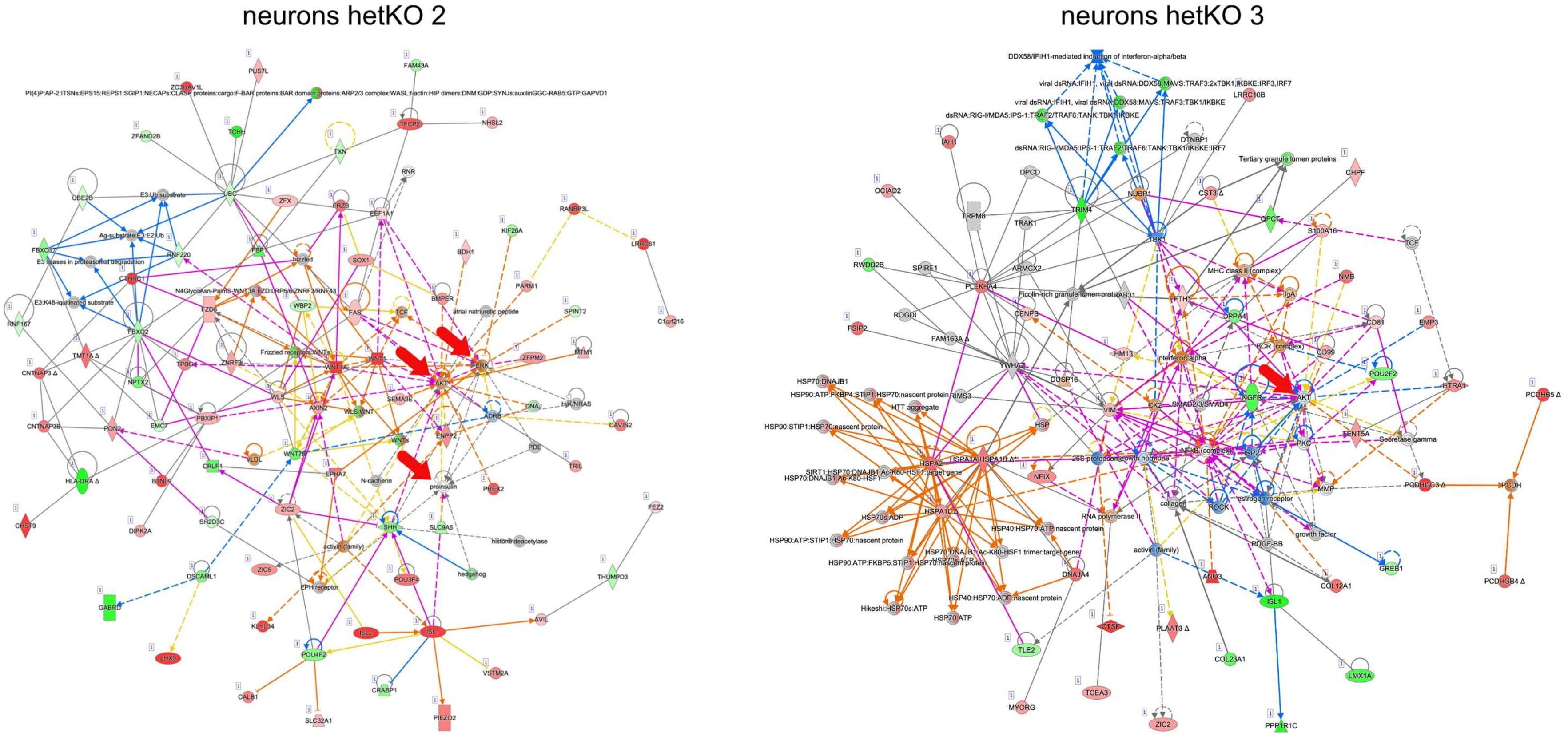


**Supplementary Figure 16. Network enrichment analysis of transcriptomic data in hetKO 2 and hetKO 3 neurons.** The three most significantly enriched networks for all DEGs were merged (IPA). The network enrichment analyses point towards an involvement of insulin signaling (insulin, AKT, ERK) in hetKO 2 neurons, which is comparable to what we observed in homKO 1 and hetKO 1 neurons. For hetKO 3, only AKT was found as a gene overlapping with the analyses for the other KO cell lines (indicated by red arrows). Genes: green: downregulated in expression, red: upregulated in expression, blue: predicted to be inhibited, orange: predicted to be activated, grey: differentially expressed but not significantly, white: not present; arrows: purple: overlap between networks, orange: predicted positive effect, blue: predicted negative effect, yellow: inconsistent/unpredicted effect, grey: no predicted effect.

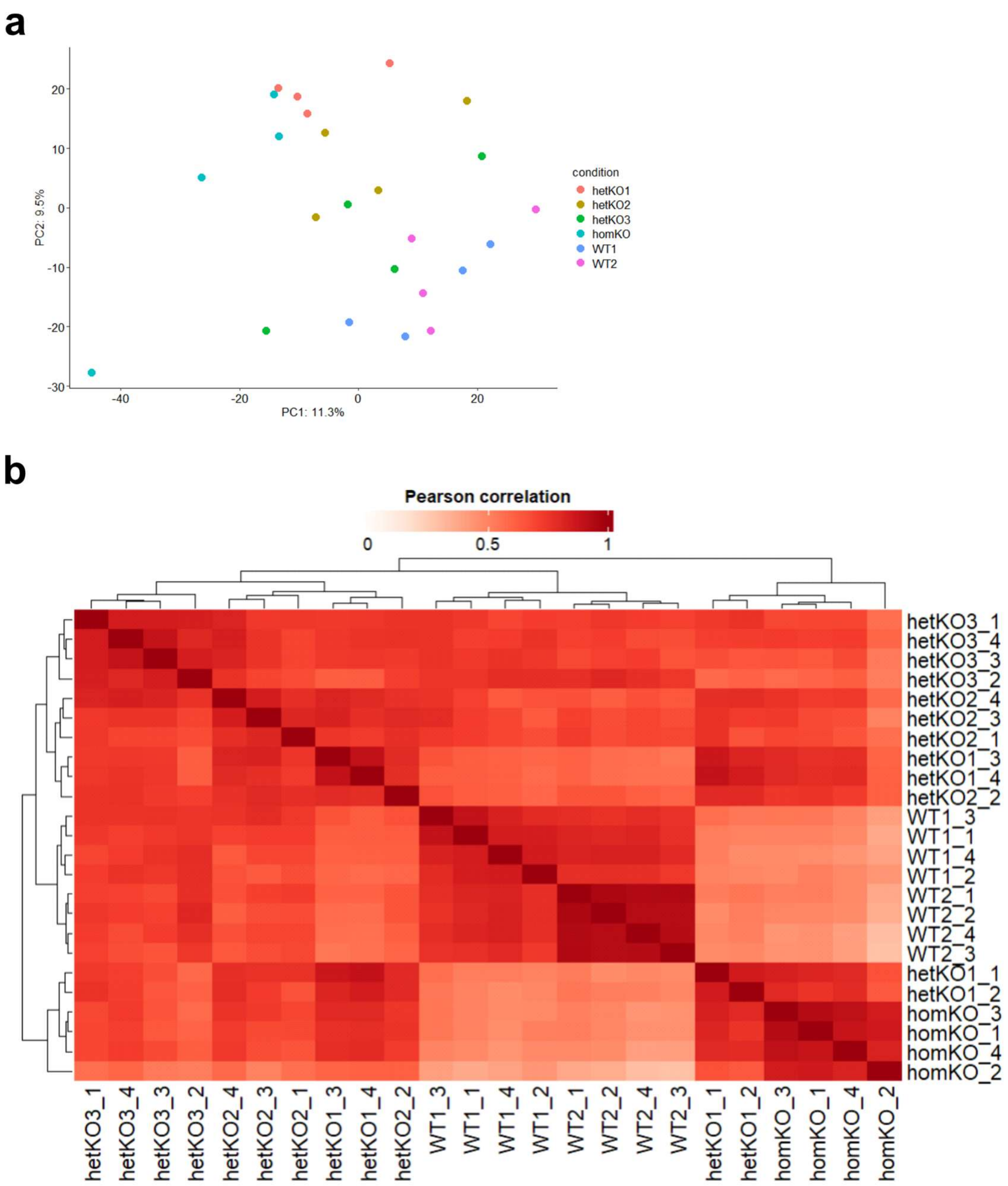


**Supplementary Figure 17. Quality assessment of proteomic data from neuronal cell lines. a** PCA of proteomic data from WT 1, WT 2, homKO 1, hetKO 1, hetKO 2, and hetKO 3 neurons. PC1 and PC2 explain 11.3% and 9.5% of variance. In the PCA plot WT 1 and WT 2 are separated from homKO 1, hetKO 1, and hetKO 2. HetKO 3 resides between the WT and the other KO lines. **b** Heatmap displaying pairwise Pearson correlation coefficients (based on $log_2$-transformed protein intensities) across all biological replicates with the dendrogram reflecting hierarchical clustering based on correlation distances. The correlation matrix highlights clear grouping by genotype. One replicate of hetKO 1 clusters with homKO 1, suggesting shared proteomic features likely due to the deleted allele in hetKO 1 being on the active allele. Three replicates of hetKO 1 group closely with hetKO 2, indicating strong similarity in protein expression profiles. HetKO 3 forms a distinct sub-cluster but remains closer to other hetKO samples than to WT samples. WT and all hetKO replicates exhibit higher inter-sample similarity compared to homKO 1, which shows the most divergent proteomic profile, indicating the strongest impact of full *KCNQ1* deletion on protein expression.

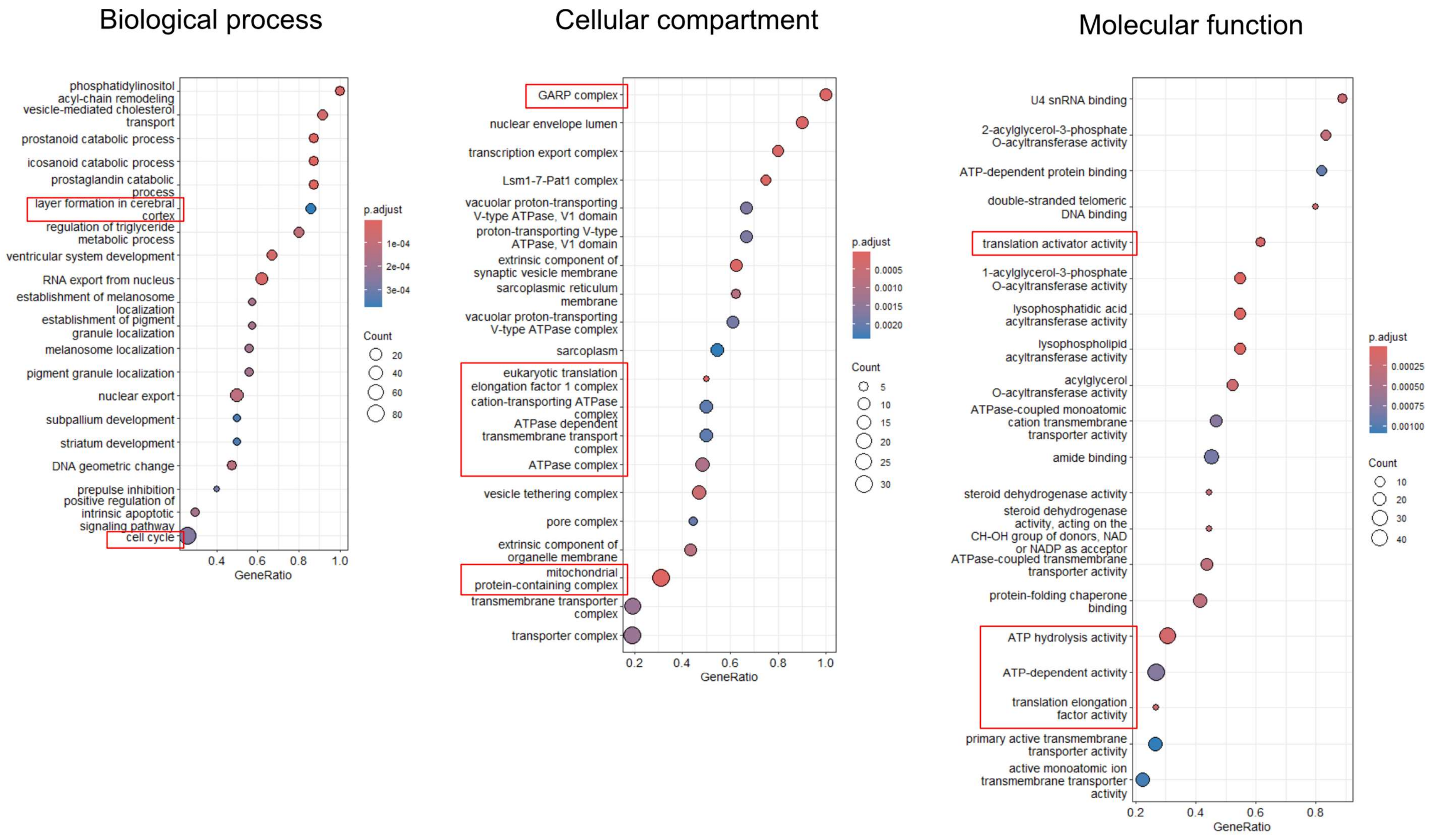


**Supplementary Figure 18. GSEA comparing homKO 1 and WT 1 neurons.** All nominal significant proteins and their log2fold changes were taken for the analysis. The top 20 affected biological processes, cellular compartments, and molecular functions are shown. Most interesting findings were labelled in red boxes and are summarized in the main Table 1.

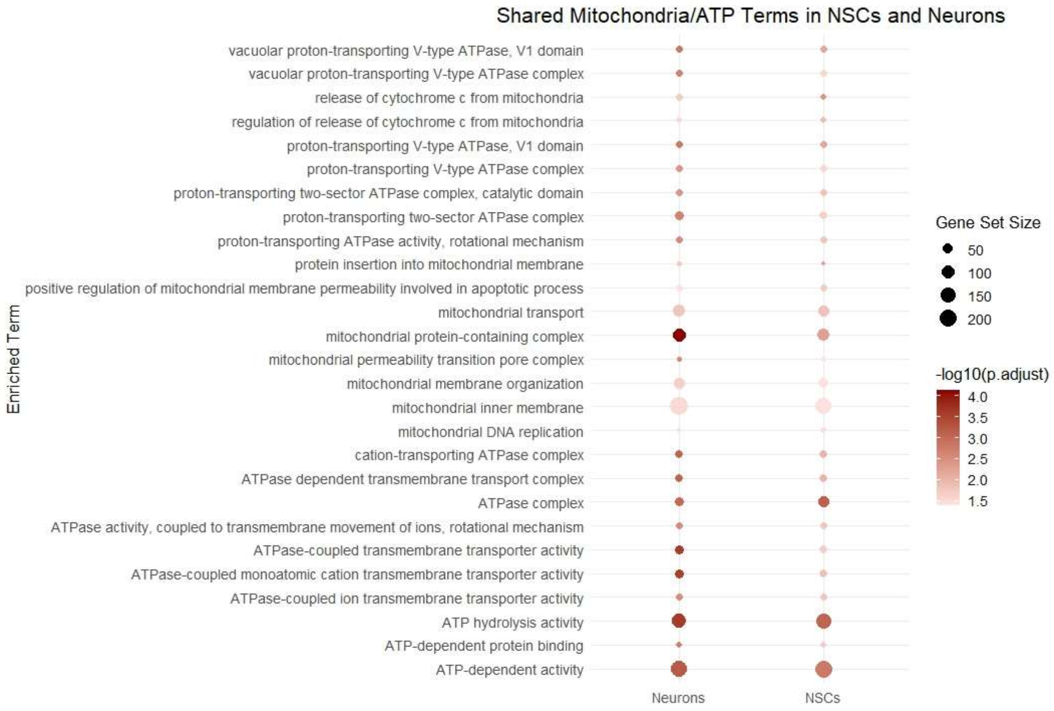


**Supplementary Figure 19. GSEA focusing on mitochondria and ATP-related GO terms in homKO 1 / WT 1 NSCs and neurons.** Several mitochondrial (mt) functional aspects showed enrichment of downregulated proteins in both cell types, including cytochrome c release, protein insertion into the mt membrane, mt transport, regulation of mt membrane permeability in apoptosis, and mt membrane organization (see also Suppl. Table 2 Tab 3 and 5). Mt DNA replication was found enriched among the upregulated proteins. Overlapping cellular compartments (significant GO terms) are: mt protein-containing complex, the inner membrane, and the permeability transition pore complex. Several ATP-related processes were found enriched among the upregulated proteins such as ATPase activity, ATP-dependent activity, and among the downregulated proteins such as ATP-dependent protein binding.

### Supplementary Materials + Methods

#### IPSC culturing

IPSCs were cultured in mTeSR Plus medium (100-0276, StemCell Technologies), seeded on Geltrex (A1413302, Thermo Fisher) and either split with Gentle Cell Dissociation Reagent (GCDR; clump splitting; 100-0485, StemCell Technologies) or Accutase (single cell splitting; A1110501, Thermo Fisher), depending on the experiment. For single cell splitting with Accutase, Rho kinase (ROCK) inhibitor (Y27632; HY-10583, Hycultec) was used for 24h after the split to increase cell survival. Cells were split when reaching 70-80% confluency (every 4-5 days).

#### Reverse transcription and quantitative PCR (qPCR)

RNA was isolated from cells with the Quick-RNA Miniprep Kit (R1057, Zymo Research). RNA was subsequently transcribed into cDNA by using the VILO Kit (11754050, Thermo Fisher) and cDNA was diluted 1:8. For qPCR we used qPCRBIO SYGreen Mix Lo-Rox (PB.2011-51, NIPPON Genetics). Gene expression was quantified according to the standard curve method and normalized to the reference genes GAPDH and SDHA. Primers were designed with NCBI Primer blast. Primer sequences can be found in Supplementary information.

#### Immunocytochemistry

100.000 cells were seeded on coverslips in a 24-well format and after growing 2-3 days fixed with 4% PFA for 15 min at room temperature (RT). For immunocytochemistry, cells were permeabilized with 0,1% Triton X-100 in PBS for 6-7 min at 4°C. Cells were blocked for 1h at RT in blocking buffer (10% NGS in PBS). Primary antibody was incubated over night at 4°C. On the next day, cells were washed 3x with PBS, incubated with secondary antibody 1:500 for 1h at RT, and washed again with 3x PBS. If a Hoechst (H3570, Life Technologies) staining was performed, cells were incubated for 5 min with 1:5000 Hoechst in PBS, then washed again 3x with PBS. Afterwards, cells were washed 3x with water and mounted on a slide with Aqua-Poly/Mount.

#### Karyotyping

Chromosome analysis was performed according to standard cytogenetic methods by treatment with trypsin, followed by Giemsa staining, in the Cytogenetics Laboratory of the Institute of Human Genetics (Heidelberg, Germany). Karyotype description was done according to the International System for Cytogenetic Nomenclature (2014). At least 10 metaphases were analysed for each clone.

#### PCR and Sequencing

DNA was isolated by using the Quick-DNA Miniprep Plus Kit (D4068, Zymo Research). PCR was performed with HotStarTaq polymerase (203205, Quiagen). Primers were design by using NCBI Primer blast and can be found in Supplementary information. Samples were sequenced by Azenta Life Science.

#### RT-PCR

To analyse the different *KCNQ1* transcripts (Suppl. Fig. 6), a regular PCR was performed (see section 'PCR and Sequencing'). Instead of using DNA, transcribed cDNA was used as PCR input to only amplify expressed transcripts on mRNA and not DNA level. Primers were design by using NCBI Primer blast and can be found in Supplementary information.

#### Trilineage Differentiation and analysis

To differentiate iPSCs into all three germ layers, a Trilineage Differentiation Kit (05230, StemCell Technologies) was used. After harvesting the cells, RNA was isolated and transcribed into cDNA. This cDNA was used to measure specific marker genes for each germ layer in iPSCs and the respective differentiation. Expression in differentiated cells was normalized to iPSC expression.

**Analysis of proliferation and apoptosis**

To assess the proliferation and apoptosis rate of NSCs, cells were split with Accutase and 25.000 cells were seeded in a 96-Well on Geltrex coating in NEM. 16h after seeding the cells, they were treated with two different dyes simultaneously: Nuclight Rapid Red Dye (4717, Sartorius) to examine proliferation and Annexin V Dye (4642, Sartorius) to examine apoptosis. Additionally, cells were treated either with JNJ303 (1 µM), ML277 (1 µM, 2 µM), Metformin (0.5 mM) or DMSO/water (mock). Immediately after treatment, cells were placed into the IncuCyte system and the recording was started within 30 min. Cells were recoded for 48h every 2h. To analyse the apoptosis rate, values were normalized to the number of cells (determined by Nuclight Red Dye). Proliferation and apoptosis rate were normalized to timepoint 0h and each timepoint was normalized to its corresponding timepoint of either respective WT control or respective mock treatment which was set to the value 1.

**Neuronal differentiation protocol 2**

This differentiation protocol was only used for one set of neurite outgrowth measurements. Neurons were generated according to the protocol by Qi et al.[2], adapted by Dr. Magdalena Laugsch and colleagues. IPSCs were seeded in a 6-well on Geltrex with 250.000 cells in mTeSR Plus medium. 24h after splitting, medium was changed to differentiation medium including different small molecules. Every second day, media with small molecules was changed (see details in table below). On day 8, cells reached an NSC-like state and were split in a 24-well on Ornithin-Geltrex coating with 380.000 cells in NDM + PD0325901 + SU5402 + DAPT + Chir99021. Cells were placed into the IncuCyte system immediately after splitting and measured for 48h.

**Overview of medium composition and small molecules for day 0-7 of protocol 2.**

| Day of differentiation | Material | Concentration |
|---|---|---|
| Day 0 + Day 1 | DMEM F12 + Glutamax | 84% |
| | Knockout Serum Replacement | 15% |
| | MEM-NAC | 1% |
| | β-Mercaptoethanol | 100 µM |
| | LDN193189 | 250 nM |
| | SB431542 | 10 µM |
| | XAV939 | 5 µM |
| Day 2 + Day 3 | DMEM F12 + Glutamax | 84% |
| | Knockout Serum Replacement | 15% |
| | MEM-NAC | 1% |
| | β-Mercaptoethanol | 100 µM |
| | LDN193189 | 250 nM |
| | SB431542 | 10 µM |
| | XAV939 | 5 µM |
| | PD0325901 | 1 µM |
| | SU5402 | 5 µM |
| | DAPT | 10 µM |
| Day 4 + Day 5 | DMEM F12 + Glutamax | 72% |
| | Knockout Serum Replacement | 10% |
| | MEM-NAC | 1% |

| | | |
|---|---|---|
| | β-Mercaptoethanol | 66.6 µM |
| | Neurobasal | 16.60% |
| | Glutamax | 0.16% |
| | N2 | 0.33% |
| | B27 | 0.66% |
| | LDN193189 | 250 nM |
| | SB431542 | 10 µM |
| | XAV939 | 5 µM |
| | PD0325901 | 1 µM |
| | SU5402 | 5 µM |
| | DAPT | 10 µM |
| Day 6 + Day 7 | DMEM F12 + Glutamax | 51% |
| | Knockout Serum Replacement | 5% |
| | MEM-NAC | 1% |
| | β-Mercaptoethanol | 33.3 µM |
| | Neurobasal | 33.3% |
| | Glutamax | 0.33% |
| | N2 | 0.66% |
| | B27 | 1.33% |
| | PD0325901 | 1 µM |
| | SU5402 | 5 µM |
| | DAPT | 10 µM |

**Electrophysiology**

Spontaneous action potential firing. To determine if *in vitro* networks of induced human neurons were spontaneously active at 6 weeks, we used loose cell-attached recordings. For this, a low resistance seal (~0.5 GΩ) between the recording pipette and the cell membrane was established, and then spontaneous spikes were recorded in voltage-clamp mode. No current injection was applied through the recording pipette. Spontaneous activity was typically recorded for 2 min.

**nCounter® gene expression profiling**

To validate the RNA Sequencing results, 16 DEGs were selected for follow-up analysis using nCounter direct RNA quantification (selected genes were highlighted in bold in Suppl. Fig. 5c). Genes were selected based on the following criteria: differential expression in all or at least three *KCNQ1*-KO lines, or differential expression in two cell lines with highest and lowest log2fold changes. Genes with very similar or unclear function were excluded (Suppl. Table 1 Tab 7).

The same batch of RNA as for RNA sequencing was used. After quantity and quality control of RNA samples, an nCounter® target gene expression analysis was performed at the nCounter Core Facility (Institute of Human Genetics, Heidelberg University Hospital, Germany, https://t1p.de/nCounter-Core-Facility) using a customized nCounter® Elements TagSet (https://nanostring.com/products/ncounter-assays-panels/ncounter-custom-solutions/elements-tagsets/) containing 16 target genes and 6 reference genes (TagSet probe design, see table below). Briefly, for each hybridization reaction, 50 ng of total RNA was used to hybridize the probe set at 65 °C. Up to 7 µl of total RNA samples were combined with 2 µl nCounter customized TagSet, 5 µl hybridization buffer, and 0.5 µl probe A plus 0.5 µl probe B for a total reaction volume of 15 µl. Samples were incubated for 20h, cooled to 4 °C and then purified, immobilized on a cartridge, and measured using the nCounter® SPRINT profiler (NanoString a Bruker Company, Seattle, WA, USA, www.nanostring.com).

Normalization and analysis of the raw data was performed using the internal positive controls and the stably expressed reference genes with the freely available nSolverTM Analysis Software (version 4.0, https://www.nanostring.com/products/analysis-software/nsolver). Stably expressed reference genes were selected based on the Normfinder method (v0.953, MS Excel Add-in66). To determine the background and detection limit, the threshold was set as the geomean of negative controls (NEG_A-F, included in the assay by default) + 2*standard deviation. Gene numbers below this threshold were considered as background, those above as specific signal. Genes exceeding this threshold in at least 50% of the samples within a group were considered to be present in the respective group. Genes not present within all groups were removed from the downstream analysis.

**Transcriptome and proteome data analysis**

Ingenuity pathway analysis was used to perform network enrichment analysis for all significantly differential expressed genes (FDR < 0.05) and for the nominal significant differentially expressed proteins. The three most significant networks were merged and shown in the figures.

**Primer table**

| Experiment | Target | Sequence |
|---|---|---|
| Genome editing | gRNA 1 (chr11: 2,571,242-2,571,261; GRCh38/hg38) | GCAGGGTGTATGCTCTTCC |
| | gRNA 2 (chr11: 2,571,251-2,571,270; GRCh38/hg38) | GTTCAGGTACCCCTGCGCCC |
| | gRNA 3 (chr11: 2,572,227-2,572,246; GRCh38/hg38) | ATACACCCTGCTCTCGTCTG |
| | gRNA 4 (2,572,450-2,572,469; GRCh38/hg38) | CCCATCTGAGTGCAGCCCTC |
| | KCNQ1 intron 2 F | CCCTTTCTGGCCACTTGCA |
| KO screening | KCNQ1 intron 5 R | TTGCCCAGCTCTCTTTCCTG |
| | KCNQ1 exon 4 F | GTGTATGCTCTTCCCTGGGG |
| | KCNQ1 exon 4 R | GTGGATGGGGCGTGAGAC |
| | KCNQ1 exon 5 F | GGACACCCATGCCATCGG |
| | KCNQ1 exon 5 R | CAAGCTGTCCTAGTGTGGGC |
| | KCNQ1 WT allele F | GTTGAGCACTGACTGGAGCA |
| Assignment of deletion and allele | KCNQ1 WT allele R | GCTGGGTATATGTCCATCCCG |
| | KCNQ1 KO allele F | CGATGACCAAACAAAGGCGG |
| | KCNQ1 KO allele R | CCTCACAACTCCGGTAAGTGG |
| | SNP 1 F | GTTGAGCACTGACTGGAGCA |
| | SNP 1 R | CGGACAGGTGCATACTGGAG |
| | SNP 2 F | CCCTGCCATCTCCAGTATGC |
| | SNP 2 R | CACCATCCGGCATGTGTTTG |
| | RPS19 F | GGCCTTCCCAGGTCAAACAG |
| Off-target analysis – gRNA 1 | RPS19 R | AAGGACCATTCAGCCTGTGC |
| | ELAV4 F | AGCACTCTCCCACGTCCA |
| | ELAV4 R | ATGCACTTAGTGGGTTCAGCA |
| | NES F | TTCCAGGATCGGGGTGTACG |
| | NES R | AGTTTGGAGGCAAAGAGGGT |
| | CLDN9 F | TGAATGTGGAACCGGTGAGTC |
| | CLDN9 R | GGGGCTCAGATTTCACAGGG |
| | PCBD1 F | TTCCAGGATCGGGGTGTACG |
| Off-target analysis – gRNA 2 | PCBD1 R | AGTTTGGAGGCAAAGAGGGT |

| | | |
|---|---|---|
| | RAD51B F | AAAGCTGGTGAAAGTCAGGG |
| Off-target analysis – gRNA 3 | AC145676.2 R | GGCGTCTGTTCAGTCAGGTC |
| | ADAM15 F | CAAGCAACACTCTGCGGACCTGC |
| | ADAM15 R | ACAGCTTGACAGATTGCGCC |
| | TCEB1P20 F | AGGGAGTGCTGGAGCTAGTA |
| | TCEB1P20 R | CAAGTACTGCACGGCTTGTC |
| | MCF2L2 F | AGGGAGTGCTGGAGCTAGTA |
| | MCF2L2 R | CAAGTACTGCACGGCTTGTC |
| | LARGE F | TTGCTCTTCCGTATCTACCACA |
| | LARGE R | GAGGCCAGAGAGCTTTCAAGT |
| | PKHD1 F | TGCCTACTCCCAAGCTCAGAA |
| | PKHD1 R | TGCAACTTTGTCCCTCGATGT |
| | SOX17 F | ACGCCGAGTTGAGCAAGA |
| Off-target analysis – gRNA 4 | RAD51B R | AGTTTGGAGGCAAAGAGGGT |
| | PRR5L F | AAACGTGTCACCAGCCCG |
| | PRR5L R | ACCCACCACTCACCTCCTT |
| | THEM6 F | CGTAGCTTAGCTGGGTGTCC |
| | THEM6 R | GTCAGGATGCCAATGCCACT |
| | AC145676.2 F | TCAGTGTGGGGTCAACAAGC |
| Trilineage differentiation | SOX17 R | TCTGCCTCCTCCACGAAG |
| | GATA6 F | ACCACCTTATGGCGCAGAAA |
| | GATA6 R | ATAGCAAGTGGTCTGGGCAC |
| | WNT3 F | ATCTACGACGTGCACACCTG |
| | WNT3 R | TGCTTCCCATGAGACTTCGC |
| | ACTA2 F | CCTATCCCCGGGACTAAGAC |
| | ACTA2 R | AGGCAGTGCTGTCCTCTTCT |
| | TBXT F | GCTGTGACAGGTACCCAACC |
| | TBXT R | CATGCAGGTGAGTTGTCAGAA |
| | FOXF1 F | CAGCCTCTCCACGCACTC |
| | FOXF1 R | CCTTTCGGTCACACATGCT |
| | DESa F | GTGAAGATGGCCCTGGATGT |
| | DESa R | CGGCTGGTTTCTCGGAAGTT |
| | NCAM1 F | AGACGCAGCCAGTCCAAG |
| | NCAM1 R | TGCTTGATCAGGTTCACTTTAATAGA |
| | PAX6 F | TCAGAGCCCCATATTCGAGC |
| | PAX6 R | CAAAGACACCACCGAGCTGA |
| | NES F | CAGCGTTGGAACAGAGGTTGG |
| | NES R | TGGCACAGGTGTCTCAAGGGTAG |
| | MAP2 F | CGAAGCGCCAATGGATTCC |
| Characterization neurons | MAP2 R | TGAACTATCCTTGCAGACACCT |
| | SYP F | CCAATCAGATGTAGTCTGGTCAGT |
| | SYP R | AGGGGTGGAGACCTAGGGTA |
| | DLG4 F | TCACAACCTCTTATTCCCAGCA |
| | DLG4 R | CATGGCTGTGGGGTAGTAGTCG |
| | GRIA2 F | ACTGACACCCCACATCGAC |
| | GRIA2 R | TCGAAAACTGGGAGCAGAAA |
| | GABRB3 F | TGAGCTCCCGCAGTTCTC |
| | GABRB3 R | CAGTGACAGTCGAGGATAGGC |
| | RELN F | GCCAAAGGACTTCACACAAGC |
| | RELN R | CATGTAATTTTGTTTGCGAGTGC |
| | CUX2 F | AAGGAGATCGAGTCGCAGAA |

| | | |
|---|---|---|
| | CUX2 R | CTCCAGGATGCTCTTGATGG |
| | POU3F2 F | TTGTGTTGCCCCTTCTTCGT |
| | POU3F2 R | TTGCCTTCGATAAAGCGGGT |
| | FOPX2 F | GCAGCAGAGATGGAAGATCA |
| | FOXP2 R | AGTTGTCTTGCTGCCTGGAG |
| | GFAP F | GCACGCAGTATGAGGCAATG |
| | GFAP R | TAGTCGTTGGCTTCGTGCTT |
| | IR F | CGAGAAGACCATCGACTCGG |
| | IR R | GACACCAGAGCGTAGGATCG |
| | IGF1R F | CTCCTGTTTCTCTCCGCCG |
| | IGF1R R | CCTGGCCCGCAGATTTCTC |
| | KCNQ1 F | ATCTCCTTCTTTGCGCTCCC |
| Expression of KCNQ1 | KCNQ1 R | CCTCCATGCGGTCTGAATGA |
| | KCNQ1OT1 F | CTGGCCCTGGAGATTATTCA |
| Expression of KCNQ1OT1 | KCNQ1OT1 R | AGCCCTAAAATGGAGGAATAGG |
| | GAPDH F | CTGGGCTACACTGAGCACC |
| Housekeeping genes | GAPDH R | AAGTGGTCGTTGAGGGCAATG |
| | SDHA F | TGGGAACAAGAGGGCATCTG |
| | SDHA R | CCACCACTGCATCAAATTCATG |
| | ND4 F | GCTCACTCACCCACCACATT |
| Mitochondria copy number and deletion | ND4 R | TCGGGGTTGAGGGATAGGAG |
| | ND2 F | ATCCACCCTCCTCTCCCTAG |
| | ND2 R | GGTGGGGATGATGAGGCTAT |
| | CO1 F | GCCCACTTCCACTATGTCCT |
| | CO1 R | GCGTAGGTTTGGTCTAGGGT |
| | β2M F | TCCACCTCTTGATGGGGCTA |
| | β2M R | GAGTACCAGGCCACCTTGAC |
| | COMT F | CCCAGCTGAAGAAGAAGTATGAT |
| RNA seq follow-up | COMT R | CCGCAGCAGGCCACATT |
| | PCDHA4 F | GCCCCAGTTTACCTGACTCT |
| | PCDHA4 R | GAGGCAGAGTAACGCCAGTC |
| | ZNF471 F | GAGCCTTGGGAGATGACGAG |
| | ZNF471 R | TCATACATGAACTGCTTCAGAGGT |
| | HDX F | CTGTGCCTTGGATTACAGGGT |
| | HDX R | AAGGGTGGCTAAGTCTCGGT |
| | TMLHE F | GGATCGGCACACTGACACTA |
| | TMLHE R | CAGTGTCCTGCCACCAGTTC |

**Antibody table**

| **Antibody** | **Company** | **Cat. No** | **Concentration** |
|---|---|---|---|
| Anti-AKT | Cell Signaling Technologies | 9272 | 1:1000 |
| Anti-p44/42 MAPK (ERK1/2) | Cell Signaling Technologies | 4696 | 1:1000 |
| Anti-Phospho-p44/42 MAPK (Erk1/2) (Thr202/Tyr204) | Cell Signaling Technologies | 4370 | 1:1000 |
| Anti-ATP5A | Abcam | Ab14748 | 1:1000 |

| | | | |
|---|---|---|---|
| Anti-GAPDH | NovusBio | NB300-328SS | 1:15000 |
| Anti-Rabbit, IRDye800CW | LI-COR | 926-32213 | 1:15000 |
| Anti-Mouse, IRDye800CW | LI-COR | 926-32212 | 1:15000 |
| Anti-Rabbit, IRDye680CW | LI-COR | 926-32223 | 1:15000 |
| Anti-Mouse, IRDye680CW | LI-COR | 926-32222 | 1:15000 |
| Anti-Nanog | NovusBio | NB100-588 | 1:100 |
| Anti-OCT4 | Abcam | ab181557 | 1:200 |
| Anti-Nestin | Abcam | ab22035 | 1:400 |
| Anti-PAX6 | BioLedgend | 901301 | 1:100 |
| Anti-SOX2 | Abcam | ab97959 | 1:100 |
| Anti-mouse Alexa Fluor 594 | Invitrogen | A11032 | 1:500 |
| Anti-rabbit Alexa Fluor 488 | Invitrogen | A11008 | 1:500 |

**TagSet probe design for nCounter analysis**

| Gene | Accession number | Sequence probe A | Sequence probe B |
|---|---|---|---|
| C9orf64 | NM_032307.3 | GAAAAGAGCCTCCAAACTTC TCCAGCAGAATTTTCCCGGTT TCATTGAGACCAATTTGGTTT TACTCCCCTCGATTATGCGGAGT | CGAAAGCCATGACCTCCGATCACTCTGC ATTAACTTCTGCGCACTATTCTCACTTTC TCGGACGCAGTTGA |
| COMT | NM_000754.3 | GCAGCAGGGGAATGGCAGTTGA GAAGGATGTACTAAGTCGAGTT ATATCTCTTTCGGGTTATATCTAT CATTTACTTGACACCCT | CGAAAGCCATGACCTCCGATCACTCCGT TAGCGTCCGTCAAGGGGAGCTTTGAAT GTTTGGTGCCCAAGTCAAGG |
| TXNRD2 | NM_006440.3 | GCAACGCCGCAAACCGTGTGCT CGTCAACAAAGCTGGCTTTGAT GTTAAACAACAGCCACTTTTTTT CCAAATTTTGCAAGAGCC | CGAAAGCCATGACCTCCGATCACTCTAG CAATGATGATGTGATCGGCTGACAGCA GAATCTCTTTCCCACCTTTG |
| ZNF471 | NM_020813.2 | GCATCTTGTCTGAGGACATTTG CACATTCTTTACATTGTTAAGA CCGCTCCACCGTGTGGACGGC AACTCAGAGATAACGCATAT | CGAAAGCCATGACCTCCGATCACTCGGT GAGGAAGAGCTGGCTGCCTGAAATGAA CTCACTAATGAGCAAGGTGT |
| ZNF736 | NM_001170905.3 | TACCACCACCAGCGCTCTTTG GTTGATGGAGTTTTGAAACC TCTTTGGGACCTGGAGTTTAT GTATTGCCAACGAGTTTGTCTTT | CGAAAGCCATGACCTCCGATCACTCCCG TTTCATTCCTTCTCACTGGAAAGTACTTC CCAATGAGGGTCTCCAGC |
| ACOT2 | NM_006821.4 | GACGCTCGAAGGACAGCAGG AGATGAGGCCATTTTGAATTC | CGAAAGCCATGACCTCCGATCACTCAGG AACTGCGCCGAACTCTTCAGGCTCCATTG |

| | | | |
|---|---|---|---|
| | | AGACCTGAGCAGATAAGGTTG TTATTGTGGAGGATGTTACTACA | GTACAGCCGG |
| PCDHA4 | NM_031500.2 | ATAACTCTACAATGGCCAGAA AGTGGGAGCTGTCCCTTATCA ATGCCCTCCTTCCTTCCTGTGT TCCAGCTACAAACTTAGAAAC | CGAAAGCCATGACCTCCGATCACTCACT TGAATTCCAAATCTGGGACATTATCGT TGTTGTCTTCTACTTCCACA |
| ISL1 | NM_002202.2 | CTTTCCAAGGTGGCTGGTAAC TTTGTACTTCCACTGGGTTAGC CTGTAAGCATAAAATTGGTTTT GCCTTTCAGCAATTCAACTT | CGAAAGCCATGACCTCCGATCACTCCTG AAAAGCAGGCTGATCTATGTCACTCTG CAAGGCGAAGTCGCTCAGTA |
| NGFR | NM_002507.3 | CGAAGTCAATTCCTTCTTGCCG CATTCCCACACTGGCTGGTCAA GACTTGCATGAGGACCC GCAAATTCCT | CGAAAGCCATGACCTCCGATCACTCGT CTAGAGCTGGGAGAAATCCCCACAGG TCACAGT |
| NHLH2 | NM_005599.3 | CGCCCAGGGACTCCGGATCCGA GTGCGCCGAGCTGGGATGGTCC GAATCTCTTTCGTTGGGACGCTT GAAGCGCAAGTAGAAAAC | CGAAAGCCATGACCTCCGATCACTCTC CAGGTCCGACACGCTGCCGAGCACCT TGGTGTCCGTGC |
| PCDHGC3 | NM_032403.3 | CGCCGAGTCCGACATGGCTTTC TGGATGCCGTTGATTTGCCCAG CAGACCTGCAATATCAAAGTTA TAAGCGCGT | CGAAAGCCATGACCTCCGATCACTCTGA GATTTCGCTGAGAAACTTTCAGCGGGT TAGCGCTTGGGCGCTGGG |
| CXCL5 | NM_002994.3 | TTGGGATGAACTCCTTGCGTGGT CTGTAAACAAACGCAACGCAGCT CTCTCCTGCCAATGCACTCGATCT TGTCATTTTTTTGCG | CGAAAGCCATGACCTCCGATCACTCCC TTGGAGCACTGTGGGCCTATGGCGAA CACTTGCAGATTACTGATCATT |
| HOXB9 | NM_024017.5 | AGACAGCACTGGCTTTGCAGTCG TCACATAACTAAGAGTGAGATGG GGAACGATTGCTGCATTCCGCTC AACGCTTGAGGAAGTA | CGAAAGCCATGACCTCCGATCACTCTA AAGGACTTGGAAGAGAGACTCCCTTC CCCATTCACTTGAATACATCCC |
| MAGEE1 | NM_020932.2 | GAATCCATAAGCAGAAATAACTG CACCAGCTTCCTTGCCTTGCTCTC CAACTGAGGCTGTTAAAGCTGTA GCAACTCTTCCACGA | CGAAAGCCATGACCTCCGATCACTC CTCGGCCAATGTAGTACAGAATTCC TTTCTTTGGTATAGGCAGCTTAGTT |
| HDX | NM_001177478.1 | TGTACTTTGGTTCTCAGCTCG AAGGGAACTGCCACTTTCATA AAATCTCGCTAGGACGCAAAT CACTTGAAGAAGTGAAAGCGAG | CGAAAGCCATGACCTCCGATCACT CATATTCACCATTTGTGAATTTGGCA TATTTCTTCCTGGTCCGGGCAAGGT |
| TMLHE | XM_011531182.4 | TTCTAAGAAGCTCTGGCAATC TACCGATGGAACTTGGGCTTG CTGGTAGACCACGCGATGACG TTCGTCAAGAGTCGCATAATCT | CGAAAGCCATGACCTCCGATCACTC ATTCCATAGAGCAGAAAGTTTTGCA GAAACTTCTTCAGTCCCTCGTTGGT |
| GAPDH (HKG) | NM_001256799.1 | GCTCCTGGAAGATGGTGATGGG ATTTCCATTGATGACAAGCTTCC CGTTCCCTCAAGACCTAAGCGAC AGCGTGACCTTGTTTCA | CGAAAGCCATGACCTCCGATCACT CCGCCAGCATCGCCCCACTTGATT TTGGAGGGATCTC |
| HPRT1 (HKG) | NM_000194.1 | TGAGCACACAGAGGGCTACA ATGTGATGGCCTCCCATCTCC TTCATCACACATCCTCTTCTTT TCTTGGTGTTGAGAAGATGCTC | CGAAAGCCATGACCTCCGATCACTC CAGTGCTTTGATGTAATCCAGCAGG TCAGCAAAGAATTTATAGCCCCCCT |
| HSPD1 (HKG) | NM_002156.4 | TCAACATCTTCAGCGATTATG ACCAAAGGCTTACGGTGAGCA | CGAAAGCCATGACCTCCGATCACTC GAAGACCAACCTTTAGCCTATTCAA |

| | | | |
|---|---|---|---|
| | | TTGGCAATCACAATTCTGCGGG TTAGCAGGAAGGTTAGGGAAC | GACGAGTGTACTTAGAGCTTCTCCA |
| PSMB4 (HKG) | NM_002796.2 | CGGATCCATGAAGGAATCGGGA GTGGACGGAATGCGGTAAAACT GTCCTGCTGTTGAGATTATTGAG CTTCATCATGACCAGAAG | CGAAAGCCATGACCTCCGATCACTCTT CTGGGTCCGCGTGATTGGACCTCTGTA AAGTGCAGACGC |
| SDHA (HKG) | NM_004168.1 | TAAACCCTGCCTCAGAAAGGCCA AATGCAGCTCGCAAGCCTGCCAA AGACGCCTATCTTCCAGTTTGATC GGGAAACT | CGAAAGCCATGACCTCCGATCACTCTG CAACAGTGTGTGACCTGGTAGGAAACA GCTTGGTAACACATGCTGTAT |
| TBP (HKG) | NM_001172085.1 | GCACGAAGTGCAATGGTCTTTAG GTCAAGTTTACAACCAAGATTCAC TGTCGAACCTAACTCCTCG CTACATTCCTATTGTTTTC | CGAAAGCCATGACCTCCGA TCACTCTCCTCATGATTACC GCAGCAAACCGCTTGGGAT TATATTCGGCGTTTCGG |

### References Supplementary


1 Yan, Y. *et al.* Efficient and rapid derivation of primitive neural stem cells and generation of brain subtype neurons from human pluripotent stem cells. *Stem Cells Transl Med* **2**, 862-870 (2013). https://doi.org:10.5966/sctm.2013-0080

2 Qi, Y. *et al.* Combined small-molecule inhibition accelerates the derivation of functional cortical neurons from human pluripotent stem cells. *Nat Biotechnol* **35**, 154-163 (2017). https://doi.org:10.1038/nbt.3777
